# ALDH1A3-acetaldehyde metabolism potentiates transcriptional heterogeneity in melanoma

**DOI:** 10.1101/2023.10.28.564337

**Authors:** Yuting Lu, Jana Travnickova, Mihaly Badonyi, Florian Rambow, Andrea Coates, Zaid Khan, Jair Marques, Laura Murphy, Pablo Garcia-Martinez, Richard Marais, Pakavarin Louphrasitthiphol, Alex H. Y. Chan, Christopher J. Schofield, Alex Von Kriegsheim, Joseph A Marsh, Valeria Pavet, Owen J Sansom, Robert Illingworth, E. Elizabeth Patton

**Affiliations:** MRC Human Genetics Unit, Institute of Genetics and Cancer, The University of Edinburgh, EH4 2XU, United Kingdom; Edinburgh Cancer Research, CRUK Scotland Centre, Institute of Genetics and Cancer, The University of Edinburgh, EH4 2XR, United Kingdom; Department of Applied Computational Cancer Research, Institute for AI in Medicine (IKIM), University Hospital Essen, 45131 Essen, Germany; University of Duisburg-Essen, 45141, Essen, Germany; Cancer Research UK Manchester Institute, The University of Manchester, Alderley Park, SK10 4TG, United Kingdom; The Dermatology Centre, Northern Care Alliance NHS Foundation Trust, Salford, M6 8HD, United Kingdom; Oncodrug Ltd, Alderley Park, Macclesfield, SK10 4TG, United Kingdom; Ludwig Institute for Cancer Research, Nuffield Department of Clinical Medicine, University of Oxford, Headington, Oxford OX3 7DQ, United Kingdom; Cancer Research UK Beatson Institute, CRUK Scotland Centre, Garscube Estate, Switchback Road, Bearsden Glasgow, G61 1BD, United Kingdom; School of Cancer Sciences, University of Glasgow, Glasgow, G12 0ZD, United Kingdom; Centre for Regenerative Medicine, Institute for Regeneration and Repair, The University of Edinburgh, Edinburgh BioQuarter, EH16 4UU, United Kingdom; Department of Chemistry and the Ineos Oxford Institute for Antimicrobial Research, University of Oxford, 12 Mansfield Road, Oxford, OX1 5JJ, United Kingdom

**Author notes:** Corresponding author: EEP.

## Abstract

Cancer cellular heterogeneity and therapy resistance arise substantially from metabolic and transcriptional adaptations, but how these are interconnected is poorly understood. Here, we show that in melanoma, the cancer stem cell marker aldehyde dehydrogenase 1A3 (ALDH1A3) forms an enzymatic partnership with acetyl-CoA synthetase 2 (ACSS2) in the nucleus to couple high glucose metabolic flux with acetyl-histone H3 modification of neural crest lineage and glucose metabolism genes. Importantly, we show acetaldehyde is a metabolite source for acetyl-histone H3 modification in an ALDH1A3, dependent manner providing a physiologic function for this highly volatile and toxic metabolite. In a zebrafish model of melanoma residual disease, a subpopulation of ALDH1-high cells emerges following BRAF inhibitor treatment and targeting these with an ALDH1 suicide inhibitor, nifuroxazide, delays or prevents BRAF inhibitor drug-resistant relapse. Our work reveals that the ALDH1A3-ACSS2 couple directly coordinates nuclear acetaldehyde-acetyl-CoA metabolism with specific chromatin-based gene regulation and represents a potential therapeutic vulnerability in melanoma.

**Highlights:**

- ALDH1A3-high melanomas are in a high glucose metabolic flux and neural crest stem cell dual state.
- Nuclear ALDH1A3 partners with ACSS2 to promote selective acetyl-histone H3.
- Acetaldehyde is an acetyl source for ALDH1A3 dependent histone H3 acetylation.
- ALDH1A3 is a master regulator and pharmaceutical target for melanoma heterogeneity.

## Introduction

The perennial challenge in the clinical development of cancer therapies is that cancer cells frequently co-opt non-genetic mechanisms to switch between dynamic cellular states^1^. This cellular plasticity enables cancer cells to adapt and thrive under environmental pressures such as immune surveillance, nutrient deprivation, or therapy^2^. Across tumour types, numerous studies have exposed that hijacking of common developmental (foetal) lineage programs and substantial cell state heterogeneity often underlie tumour progression and drug resistance^3–10^. While several molecular mechanisms that can give rise to cancer heterogeneity phenotypes have been described, to make transformative progress towards curtailing tumour state transitions and enhancing treatment efficacy, a deeper understanding of the interplay between epigenetic, transcriptional, and metabolic plasticity in cancer cell biology is needed.

For many patients with advanced melanoma, systemic targeted and immune therapies have greatly improved prognosis^11–15^. However, a vexing obstacle towards eradicating melanoma cells and achieving long-term survival is that melanoma cell subpopulations undergo phenotypic transitions into dedifferentiated stem-like states, leading to innate or acquired drug resistance and tumour recurrence. In such subpopulations, a transcriptional state resembling that of neural crest stem cells (NCSC) and characterized by low activity of Melanocyte Inducing Transcription Factor (MITF) emerges^10,16–18^. This state then becomes enriched after therapy, which is predictive for patient outcomes ^6–8,10,16,17,19–24^.

In addition to transcriptional states, metabolic heterogeneity has recently come to the forefront as a mechanism influencing tumour cell survival and biology^25^. Generally considered to arise from competition for nutrients between cancer cells and tumour-associated cells in the microenvironment, metabolic heterogeneity plays a role in this phenotypic plasticity, which enables melanoma and other cancer cells to survive therapy in specific tissue and organ environments^26–33^. While the developmental neural crest state is known to be highly sensitive to metabolic deficiencies^34–38^, we lack an understanding of how metabolic states are co-opted by and coordinated with neural crest programs in melanoma.

In this study, we discover that the pan-cancer stem cell marker ALDH1A3 is a central regulator of both metabolic and stem cell transcriptional and phenotypic states in melanoma. Using melanoma cell lines and patient sample-derived low passage cells, we show that ALDH1A3 forms an enzymatic partnership with ACSS2 to consolidate a high glucose metabolic flux with a NCSC transcriptional state instructed by Transcription Factor AP-2 Beta (TFAP2B). Mechanistically, we show that ALDH1A3 activity promotes selective acetylation of histone H3, and link this to transcription of genes regulating NCSC and glucose metabolism. Critically, by tracing acetaldehyde to acetyl-histone H3, we demonstrate its role as a source for acetylated histones dependent on ALDH1A3. Our findings uncover an actionable, high dimensional metabolic-transcriptional framework that controls melanoma stem cell plasticity.

## Results

### ALDH1A3^High^ melanomas are enriched for neural crest stem cell (NCSC) and glucose metabolic states

When patients with melanoma become resistant to MAPK inhibitor therapy, their tumour cells can upregulate expression of the stem cell marker and aldehyde dehydrogenase enzyme ALDH1 **(Figure S1A)**^39–41^. Further, in experimental mouse melanoma models, we find *Aldh1a3* expression is tightly associated with dedifferentiated, neural crest and stem cell states, which have been reported to fuel cancer growth in a cellular hierarchy^42,43^ (**Figure S1B, C**). From these observations, we hypothesised that the ALDH-high metabolic activity of cancer cells is the consequence of a changed transcriptional state that contributes to melanoma stemness and plasticity.

To investigate this, we considered that ALDH-high activity (ALDH^High^) is heterogeneous in human cell lines (**Figure S1D)**, and that the predominant ALDH activity in melanoma cell line A375 is due to ALDH1A3^39^. Thus, we sorted A375 melanoma cells for the highest and lowest ALDH activity, termed ALDH^High^ and ALDH^Low^, using the flourescent substrate Aldefluor (an amino acetaldehyde) (**Figure 1A**). As demonstrated previously, these sorted A375 cell populations have differing phenotypic potential, with ALDH^High^ cells having increased tumour-initiating potential^39,41^. When we validated ALDH1A3 expression by immunocytochemistry (ICC), we were intrigued to see that ALDH1A3 was predominantly expressed in the cytosol in ALDH^Low^ cells but was enriched in the nucleus of sorted ALDH^High^ cells (**Figure 1B**).

**Figure 1:**
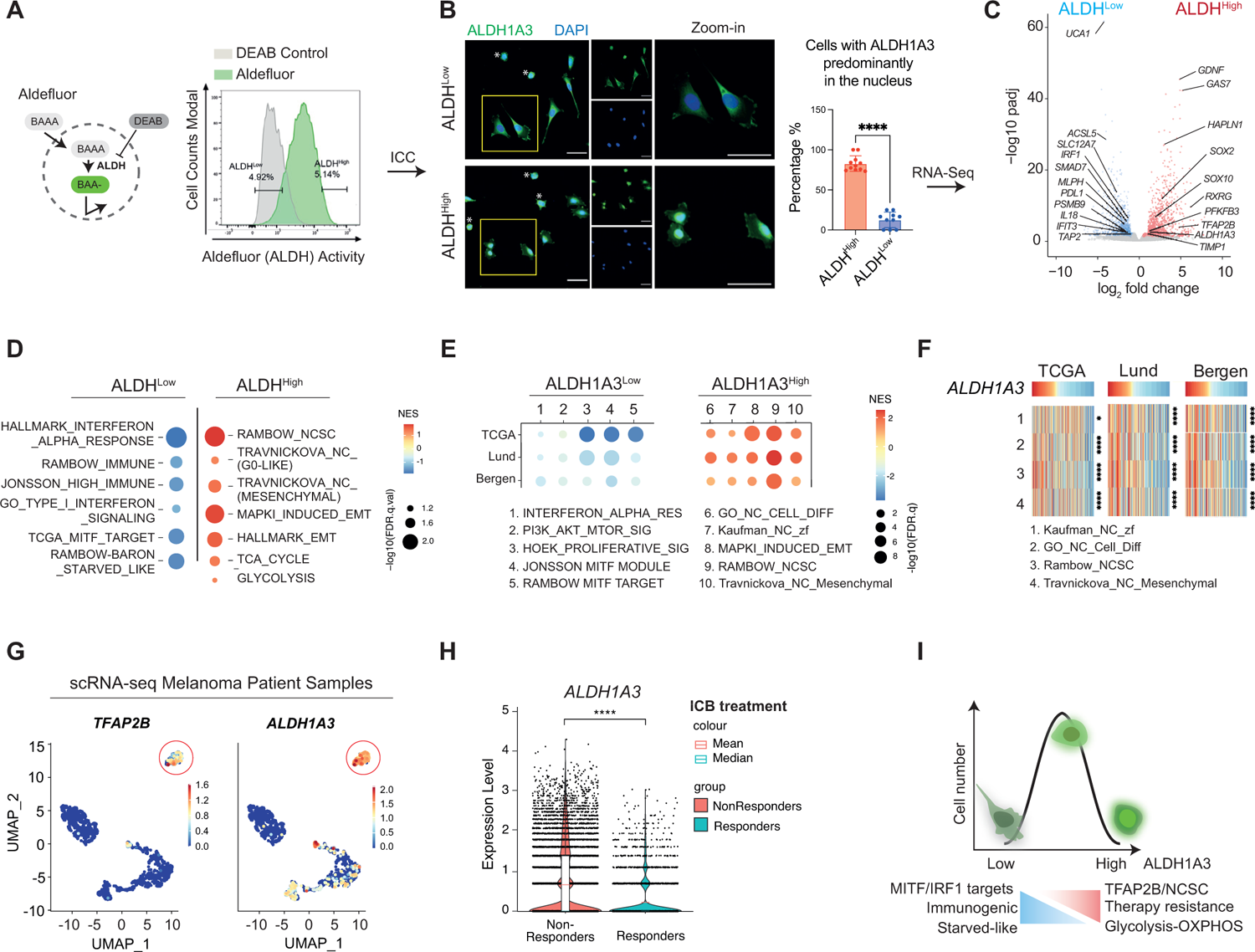
ALDH1A3^High^ melanomas are enriched for NCSC and glucose metabolic states. **A.** ALDH^High^ and ALDH^Low^ melanoma cell subpopulations. The Aldefluor assay quantifies ALDH activity in live cells by measuring the accumulated fluorescence from converted BODIPY-aminoacetaldehyde (BAAA) to BODIPY-aminoacetate (BAA-). Highest and lowest Aldefluor activity, termed ALDH^High^ and ALDH^Low^ subpopulations, were isolated by FACS. Negative control: DEAB, a pan-ALDH inhibitor. **B.** ALDH1A3 cellular localisation and levels. Immunocytochemistry (ICC) staining by fluorescence antibody labelling ALDH1A3 in A375 sorted ALDH^High^ and ALDH^Low^ cells. Scale bar = 50 µm. Quantification of cells enriched with nuclear ALDH1A3: N=2 biological repeat experiments, quantified image fields n=11 for ALDH^High^ and n= 9 for ALDH^Low^, mean±s.d.; non-paired Kolmogorov-Smirnov test, ****p<0.0001. Cells showing minimal cytoplastic content outside DAPI-stained regions (indicated with an asterisk*) were determined as unfit following FACS and thus excluded from quantification. **C.** Differential gene expression in ALDH^High^ and ALDH^Low^ melanoma cells. Volcano plot showing differentially expressed genes in ALDH^High^ (red) and ALDH^Low^ (blue) cells. DESeq2-analysis was performed using RNA-seq data of 3 biological replicants and cut-off thresholds using fold-change >1 and p.adj < 0.05. See also **Table S1**. **D, E.** Gene set enriched analysis in melanoma cells and patient samples. Dot plot of pathway analysis showing enriched terms **(D)** in ALDH^High^ and ALDH^Low^ cells and **(E)** in ALDH1A3^High^ and ALDH1A3^Low^ patient samples. Dot sizes represent -log_10_ FDR q-value (weighted Kolmogorov– Smirnov test) and colours indicate normalised enrichment score (NES). Patient groups are defined in **Figure S1G**. See also **Table S1, S2**. **F.** Heatmap of NCSC gene sets variation analysis (GVSA) score from patient samples ranked by the *ALDH1A3* level. Melanoma patient samples from TCGA, Lund, and Bergen cohorts unanimously showed positive correlation between ALDH1A3 and NCSC gene sets as annotated. *p<0.05, ****p<0.0001, Spearman’s rank correlation critical probability exact (p) value. **G.** *ALDH1A3* and *TFAP2B* scRNA-seq cluster in metastatic patient samples. UMAP feature plot showing expression level of *ALDH1A3* and *TFAP2B* in scRNA-seq of reanalysed patient samples of metastatic melanomas^51^. Red circle highlights an *ALDH1A3* and *TFAP2B* cluster. Gene expression is visualised by a colour index from blue (low) to red (high). **H.** ALDH1A3 levels are significantly higher in ICB non-responders. scRNA-Seq data from approximately 14, 2000 malignant cells from Pozniak *et al*., 2024^106^ were interrogated for *ALDH1A3* expression and early respons to ICB. **I.** Schematic of new model of ALDH1A3 activity. High levels of ALDH1A3 Aldefluor activity are associated with nuclear localization and a high glucose metabolism and TFAP2B-NC stem cell dual state. Low levels of ALDH1A3 Aldefluor activity are associated with a differentiated, immunogenic, and starved-like state.

We then conducted comparative RNA-sequencing (RNA-seq) of sorted ALDH^High^ and ALDH^Low^ cells and identified high expression of genes associated with dedifferentiated, neural crest and stem cell states (including *SOX2, SOX10, GAS7, GDNF, RXRG, TFAP2B*) in ALDH^High^ cells (**Figure 1C, D; Figure S1E Table S1, S2)**. These genes were also identified as the top marker genes associated with a less-differentiated state as measured by CytoTRACE in murine melanoma **(Figure S1F).** This molecular profile resembles a cell state present in drug-resistant melanomas that is predictive of poor outcomes for patients treated with both targeted and immunotherapy^7,8,44^.

In contrast, ALDH^Low^ cells were enriched for Interferon Regulatory Factor 1 (IRF1) target genes (*IFIT3, IL18, PDL1*), proinflammatory genes (*TAP2, PSMB9, SMAD7*) as well as MITF target pigmentation (differentiation) genes (*MLPH)* (**Figure 1C, D; Table S1)**. Metabolic transcriptional signatures were also highly distinct in ALDH^High^ and ALDH^Low^ cell populations, with glycolysis pathway genes (*PDHX, PFKFB3*) enriched in ALDH^High^ cells whereas starvation response-related fatty acid metabolism genes (*ACSL5, SLC12A7*) were enriched in ALDH^Low^ cells (**Figure 1D**). Taken together, these data show that the ALDH1A3^High^ cell state has features of both NCSC development and high glucose metabolism, while ALDH1A3^Low^ has proinflammatory and differentiation features^18,34,45,46^.

Next, we asked if the ALDH1A3-enriched cell state is present in human patient samples. To answer this question, we ranked patient samples from available datasets based on *ALDH1A3* RNA levels, and then selected *ALDH1A3-high* and *ALDH1A3-low* patients from each cohort; top and bottom 10% for TCGA^47^ and Lund (population cohort of primary and metastatic melanomas)^48^, and top and bottom 25% for Bergen dataset (Stage IV melanomas)^49^ **(Figure S1G)**. Gene expression-based melanoma subtype consensus has been previously established from these patient cohorts which successfully stratified patient prognosis independent of oncogenic genotype^50^. In our case, for the *ALDH1A3-high* patient samples, we found that the NCSC state as well as mesenchymal state (epithelial to mesenchymal transition program) were enriched (**Figure 1E, F; Figure S1H; Table S1, S2).** Analysis of a fourth independent patient sample dataset revealed a direct positive correlation between *ALDH1A3* expression and NCSC transcriptional signatures **(Figure S1I).**

We were particularly intrigued to see a strong association between *ALDH1A3* and *TFAP2B* expression across melanoma cell and mouse models, and human patient datasets, given that we have recently shown *tfap2b* marks an adult multipotent melanocyte stem cell population in zebrafish^46^ (**Figure 1C; Figure S1E-F)**. The *ALDH1A3-TFAP2B* association was isoform-specific: across the entire ALDH family, *TFAP2B* was only positively correlated with *ALDH1A3* **(Figure S1J)**. In contrast, *TFAP2A*, a related family member expressed in proliferative melanoma that shares near identical binding motif with TFAP2B **(Figure S1K)**, was negatively correlated with *ALDH1A3* **(Figure S1L)**. Given these data, we re-analysed scRNA-seq data from 15 melanoma patient samples^51^ and found that *ALDH1A3* and *TFAP2B* are co-expressed in a distinct cell cluster present in five out of 15 samples of varied mutation subtypes, indicating that there are enough cells in this state to form a cluster, and that this cluster does not simply come from a single patient or genotype (**Figure 1G, Figure S1M)**. Thus, in five independent melanoma patient datasets *ALDH1A3* expression correlates with *TFAP2-NCSC* expression.

In contrast to *ALDH-high* melanomas, and in agreement with our RNA-seq analysis (**Figure 1D),** *ALDH1A3-low* patient samples were enriched in proinflammatory gene signatures and MITF pigmentation target genes (**Figure 1E, Figure S1H, L)**. Proinflammation associated gene expression signatures have predictive value as biomarkers of the immune checkpoint blockade (ICB) response, which has dramatically improved patient outcomes for advanced stage melanoma^52^. We addressed how *ALDH1A3* expression levels correlate with patient response to ICB. We analysed scRNA-seq of >14,000 cells from treatment-naive stage III/IV malignant melanoma patients receiving ICB (nivolumab or ipilimumab and nivolumab) and found ALDH1A3-high expression associated with “non-responders” while ALDH1A3-low expression was associated with “responders” to ICB (**Figure 1H**).

Collectively, our data thus far support a model in which high ALDH1A3 metabolic activity segregates with transcriptional activation of *TFAP2B-NCSC*-driven developmental stem cell pathways, high expression of glucose metabolism genes, and resistance to both targeted and ICB therapy, while low ALDH1A3 metabolic states have features of melanocyte differentiation, low glucose metabolism and robust immunogenicity (**Figure 1I**).

### TFAP2B promotes stemness and dedifferentiation in ALDH1A3^High^ melanoma cells

We next asked whether ALDH-associated melanoma stem cell phenotypes are directly regulated by TFAP2B. To do this, we selected two widely used cutaneous melanoma cell lines, A375 and C089, in which *ALDH1A3* and *TFAP2B* are expressed at high levels in A375, while both genes are expressed at a relatively moderate level in C089 (**Figure 2A**). Both cell lines bear wildtype p53 and BRAF(V600E) mutations^39,53^. Next, we engineered these two melanoma cell lines to knock out or over-express *ALDH1A3* **(Figure S2A)**. By western blot analysis, we validated that loss of ALDH1A3 led to reduced TFAP2B protein levels, while ALDH1A3 overexpression upregulated TFAP2B (**Figure 2A, B)**. Using the Aldefluor assay, we validated *ALDH1A3* knockout (KO) and overexpression (OE) has significantly shifted ALDH activity toward low and high states respectively (**Figure 2C, D)**. Consistently, RT-qPCR analysis in both cell lines revealed that loss of *ALDH1A3* led to reduced *TFAP2B-NCSC* gene expression, concomitant with increased expression of *IRF1* and *MITF* target genes, while this response was reversed in cells that overexpressed *ALDH1A3* (**Figure 2E**). Taken together, these data support that ALDH1A3 is a direct regulator of the *TFAP2B-NCSC* state.

**Figure 2:**
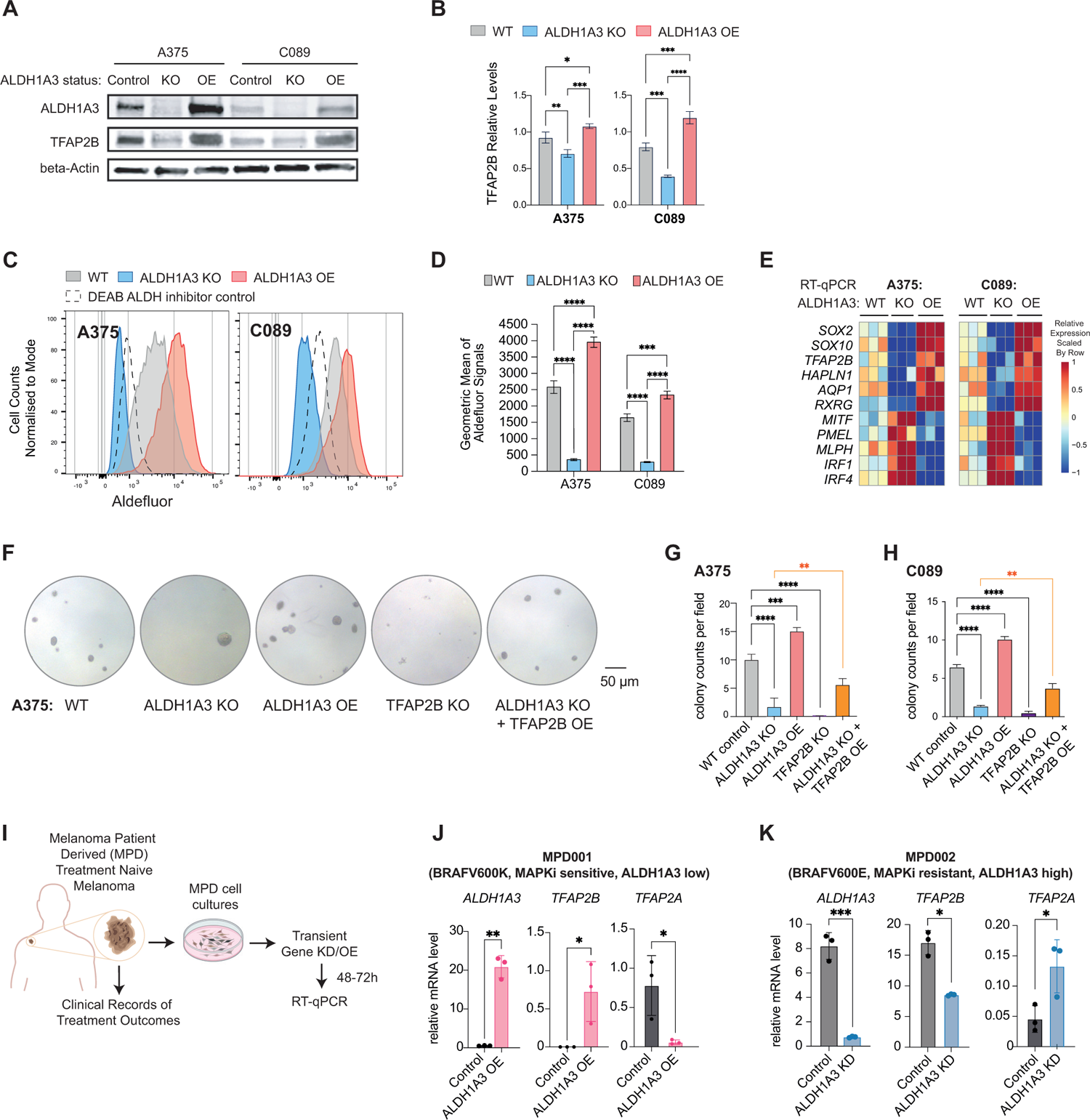
TFAP2B promotes stemness and dedifferentiation in ALDH1A3^High^ melanoma cells. **A, B.** TFAP2B protein levels are dependent on ALDH1A3. Western blot and quantification of ALDH1A3, TFAP2B protein expression in ALDH1A3 control, KO and OE cells. n=3, mean±s.d.; one-way ANOVA with Tukey’s test for multiple comparisons. **C, D.** ALDH1A3 melanoma cell models. Aldelfuor activity and quantification in *ALDH1A3* knockout (KO), over-expression (OE), and vehicle control cells (A375 and C089 cells). n=3, mean±s.d.; one-way ANOVA with Tukey’s test correction for multiple comparisons. **E.** The ALDH1A3-neural crest stem cell (NCSC) gene signature. RT-qPCR results of NCSC signature genes in control versus *ALDH1A3* KO and OE cells (A375 and C089 cells, 3 bio-replicants with 3 technical replicates each). **F, G, H.** *TFAP2B* rescues *ALDH1A3* activity in colony assays. Representative images and quantification of colonies formed by A375 cell control, *ALDH1A3* KO, *ALDH1A3* OE, *TFAP2B* KO, and combined *ALDH1A3* KO with *TFAP2B* OE conditions. Quantification on both A375 and C089 cells across each condition group, n=3 per condition, mean±s.d. one-way ANOVA with Tukey’s test for multiple comparisons. **I,** Schematics of the establishment of low passage melanoma patient derived (MPD) cells and experimental design for molecular profiling of ALDH1A3 target genes. **J, K.** *ALDH1A3* overexpression promotes *TFAP2B* expression in low-passage patient cells. MPD 001 and 002 cells are ALDH1A3-low, MAPKi sensitive and ALDH1A3-high, MAPKi resistant, respectively. RT-qPCR of *ALDH1A3, TFAP2A, TFAP2B* expression in control cells and cells over-expressing *ALDH1A3*. n=3, mean±s.d.; Multiple non-paired t-test corrected with Holm-Sidak’s method. *P<0.05; **P<0.01; ***P<0.001; ****P<0.0001.

Next, to test the phenotypic response to altering *ALDH1A3* and *TFAP2B*, we found that loss of *ALDH1A3* led to decreased colony formation while *ALDH1A3* overexpression increased colony formation in both A375 and C089 cell models (**Figure 2F-H**), consistent with the pan-cancer effect of ALDH activity in cancer stem-like states^39,40,54^. Remarkably, loss of *TFAP2B* via CRISPR-Cas9 gene knockout in melanoma cells phenocopied *ALDH1A3* deficiency in colony assays (**Figure 2F-H**). We were unable to assess the impact of *TFAP2B* overexpression alone in colony assays due to neuronal-like differentiation, consistent with TFAP2B being a powerful transcriptional regulator of the neural crest. However, in *ALDH1A3* KO cells, overexpression of *TFAP2B* partially rescued the stemness deficiency, leading to enhanced colony formation capacity in both *ALDH1A3* and *TFAP2B* KO cells (**Figure 2F-H, Figure S2B).** These data indicate that TFAP2B is a downstream mediator of ALDH1A3 that sustains cancer stemness in melanoma.

Next, we set out to directly test if ALDH1A3 mediates *TFAP2B* expression in patient samples. We obtained low-passage human melanoma cells with associated clinical treatment response data, termed melanoma patient-derived (MPD) cells. With this new resource, we employed MPD001 and MPD002 cells from patients diagnosed with BRAFV600-mutant melanoma, with relatively low and very high levels of ALDH1A3, respectively (**Figure 2I; Figure S2C**). The clinical records show that the patient’s disease for MPD001 donor was sensitive initially but then continued to progress on immune therapy (ipilimumab) as well as targeted therapies (vemurafenib), while the patient’s disease for MPD002 was innately resistant to vemurafenib as well as ipilimumab. When we over-expressed *ALDH1A3* in the MPD001 cells, we found that *TFAP2B* gene expression was upregulated whereas expression of the melanocyte differentiation gene *TFAP2A* was downregulated (**Figure 2J**). Moreover, these changes upon *ALDH1A3* overexpression segregated with an observable loss of pigmentation, indicative of melanocyte de-differentiation into a stem cell-like state **(Figure S2D)**. In contrast, knockdown (KD) of *ALDH1A3* in the MPD002 cells led to decreased *TFAP2B* expression, whereas *TFAP2A* was increased in expression (**Figure 2K**). Thus, in human melanoma cell lines and patient cell cultures, *ALDH1A3* expression is, in and of itself, sufficient to promote *TFAP2B* gene expression.

### ALDH1A3^High^ cells use glucose while ALDH1A3^Low^ cells rely on acetate for Acetyl-CoA production

Next, we sought to explore relationships between ALDH1A3 function and the metabolic states we identified by RNA-seq in cells and in patients (**Figure 1**). To this end, we traced ^13^C_6_-labelled glucose in two A375-based melanoma cell models: (1) WT control cells (that express high levels of *ALDH1A3*) and *ALDH1A3* knockout cells (*ALDH1A3* KO) for 24 hours labelling, and (2) non-engineered cells labelled for 12 hours, and then sorted for ALDH1A3^High^ and ALDH1A3^Low^ activity (**Figure 3A, Table S3)**. We did not find any difference in ^13^C_6_-glucose uptake between cells having different levels of *ALDH1A3* (**Figure 3B**), despite the increased expression of glycolysis genes in ALDH1A3^High^ cells determined by RNA-seq analysis. Instead, we observed significantly higher glucose-derived carbon flux in cells with high ALDH1A3 activity, as evidenced by an increase in ^13^C-labelled carbon in pyruvate and TCA cycle intermediates, including citric acid, α-ketoglutaric acid, succinic acid, and malic acid (**Figure 3C-D**). Conversely, in *ALDH1A3* KO and ALDH1A3^Low^ cells, glucose-derived carbon was converted into ketone bodies such as aceto-acetate, a metabolite synthesized from acetyl-CoA, reflecting a starvation-like metabolic state (**Figure 3E**). Despite the lower glycolytic flux, *ALDH1A3* KO cells produced higher levels of lactate both in cells and secreted into the culture media (**Figure 3F**). These data indicate that ALDH1A3^High^ cells are primed to execute glycolysis and oxidative phosphorylation (OXPHOS), whereas ALDH1A3^Low^ cells exist in a starvation-like state and secrete lactate. Consequently, we reasoned these states exert differential metabolic and epigenetic effects on surrounding cancer and stromal cells ^55,56^.

**Figure 3:**
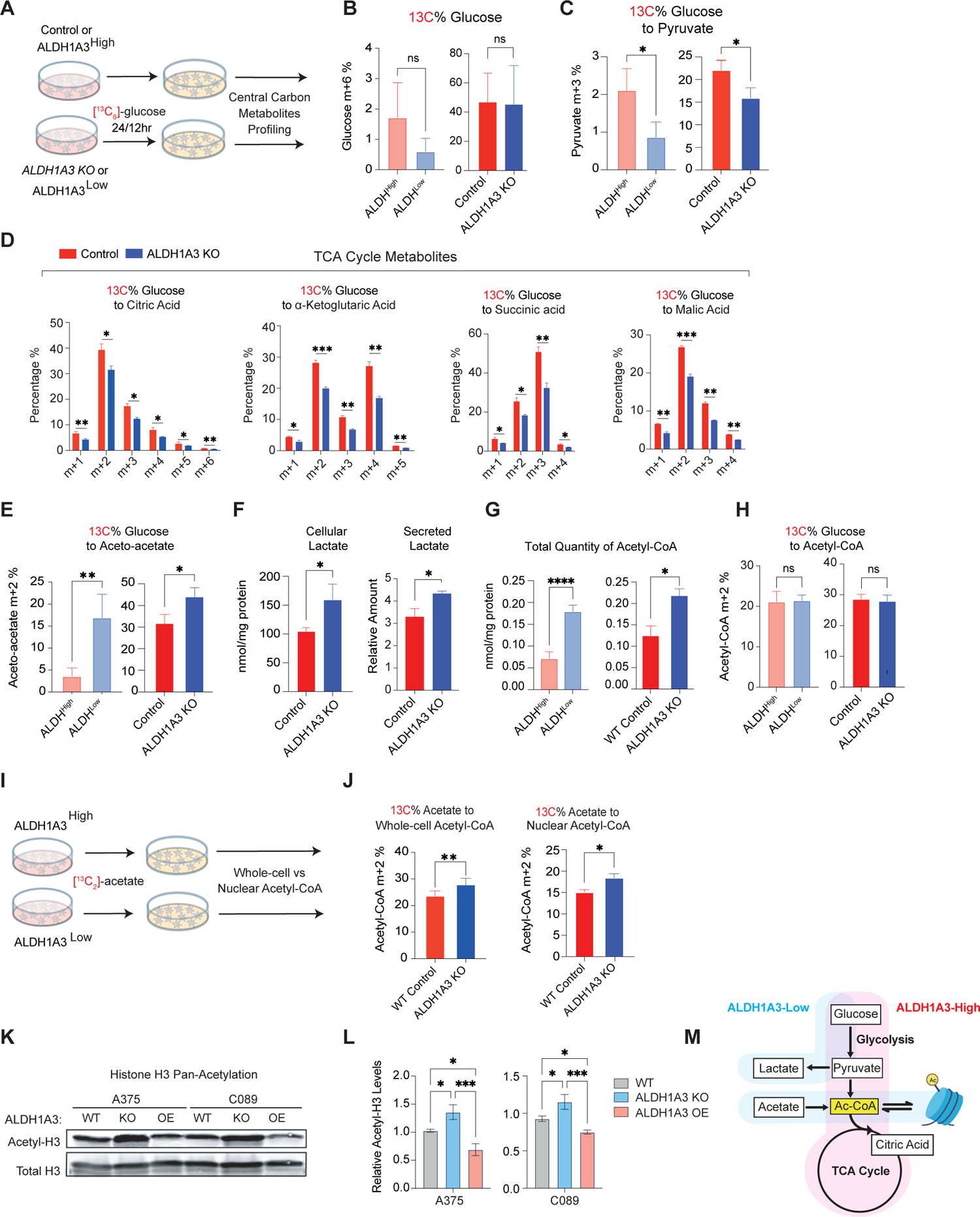
ALDH1A3^High^ cells use glucose while ALDH1A3^Low^ cells rely on acetate for Acetyl-CoA production. **A-E.** Schematic of ^13^C_6_ glucose tracing experiment. Central carbon metabolites were profiled by UPLC-MRM/MS in panel (**B**) Glucose (**C**) Pyruvate (**D**) TCA cycle metabolites: citrate, alpha-ketoglutarate, succinic acid, malic acid, and (**E**) aceto-acetate. (n=3 for WT vs ALDH1A3 knockout cells; n=5 for ALDH^High^ vs ALDH^Low^ cells. Multiple paired t-test corrected with Holm-Sidak’s method.) **F**. *ALDH1A3* KO cells generate and secrete lactate. Intracellular lactate measured by targeted UPLC-MRM/MS. (n=3, Multiple paired t-test corrected with Holm-Sidak’s method). Secreted lactate levels were measured by a colorimetric lactate assay kit and normalised to standards. (n=3, unpaired student t-test.) **G, H**. *ALDH1A3* KO cells generate more acetyl-CoA but not from glucose. Intracellular glucose w/o 13C_6_-labelling and **(H)** total acetyl-CoA measured by targeted UPLC-MRM/MS. (n=3, Multiple paired t-test corrected with Holm-Sidak’s method. **I, J.** Schematic of ^13^C_2_ acetate tracing experiment. ^13^C_2_ acetate tracing experiment designed for ^13^C-incoporation profiling by HPLC in (**L**) whole-cell and nuclear Acetyl-CoA. n=5, two-way ANOVA corrected with Holm-Sidak’s method.) **K, L.** Western blot analyses of pan-acetyl-histone H3 (Acetyl-K9+K14+K18+K23+K27), and total histone H3 protein levels in lysates of A375 and C089 cell lines with vehicle control (WT), *ALDH1A3* knockout (KO) or *ALDH1A3* overexpression (OE) respectively, with total histone H3 probed as loading control and **(L)** quantification (n=3, mean±s.d.; one-way ANOVA with Tukey’s correction for multiple comparisons.) **M.** Schematic of ALDH1A3 and metabolic states. ALDH1A3 acts through two different sources of Acetyl (Ac)-CoA production: high glucose flux generates high levels of pyruvate, leading to Ac-CoA in ALDH^High^ cells, while ALDH^Low^ cells preferentially uptake acetate as an Ac-CoA source. *P<0.05; **P<0.01; ***P<0.001; ****P<0.0001.

Acetyl-CoA, an acetyl group donor in biochemical reactions, is generated from pyruvate through glycolysis or through oxidation of long-chain fatty acids or certain amino acids, and its availability is known to shape both metabolic processes as well as epigenetic regulation^57^. Unexpectedly, we found that total levels of acetyl-CoA increased in ALDH1A3^Low^ cells despite decreased glycolysis and reduced TCA cycle flux (**Figure 3G**). This increase in the total levels of acetyl-CoA was not derived from a glucose source, as indicated by ^13^C_6_ glucose tracing (**Figure 3H**). Given that cancer cells under metabolic stress can support as much as half of their lipid synthesis by using acetate as a substrate^58^, we hypothesized that ALDH1A3^Low^ melanoma cells supplement their carbon source by switching from glucose to acetate. When we traced ^13^C_2_-acetate incorporation into acetyl-CoA (**Figure 3I),** we found that *ALDH1A3* KO cells had a higher percentage of acetate derived acetyl-CoA relative to control cells, both within the whole cell and in the nuclear fraction (**Figure 3J**).

To address the potential link between metabolic processes and chromatin modification, we found that the increase in acetyl-CoA in *ALDH1A3* KO and sorted ALDH^Low^ Cells was also concomitant with a significant increase in acetyl-histone H3 (∼30%), as assessed by western blot and mass spectrometry analysis of histones **(Figure K, L; Figure S3A)**. Consistently, overexpression of ALDH1A3 led to a ∼15% decrease in acetyl-histone H3 **(Figure3 K, L; Figure S3A)**. Thus, we conclude that cells with high levels of ALDH1A3 rapidly metabolise glucose to generate pyruvate, while cells with low ALDH1A3 activity preferentially use acetate as a carbon source for acetyl-CoA (**Figure 3M**).

### Pyruvate-derived acetaldehyde serves as an acetyl source for histone H3-acetylation

In human cells, pyruvate dehydrogenase (PDH) can generate acetaldehyde, an ALDH substrate, from pyruvate^59^. We investigated if this pyruvate-derived acetaldehyde pool could serve as a source of acetyl-groups for histone H3 modification in melanoma. To answer this question, we traced ^13^C_2_-labelled acetaldehyde to histones and performed mass spectrometric analyses (**Figure 4A**). Indeed, in our two independent melanoma cell lines (control vs *ALDH1A3* KO), we detected higher levels of ^13^C-labelled acetylated histone H3 proteins in cells with high ALDH1A3 activity, with the incorporation at H3K14ac and H3K23ac especially responsive to ALDH1A3 levels (**Figure 4B, left panel).** To our knowledge, this is the first demonstration that acetaldehyde can be a direct source for acetylation of histone H3.

**Figure 4:**
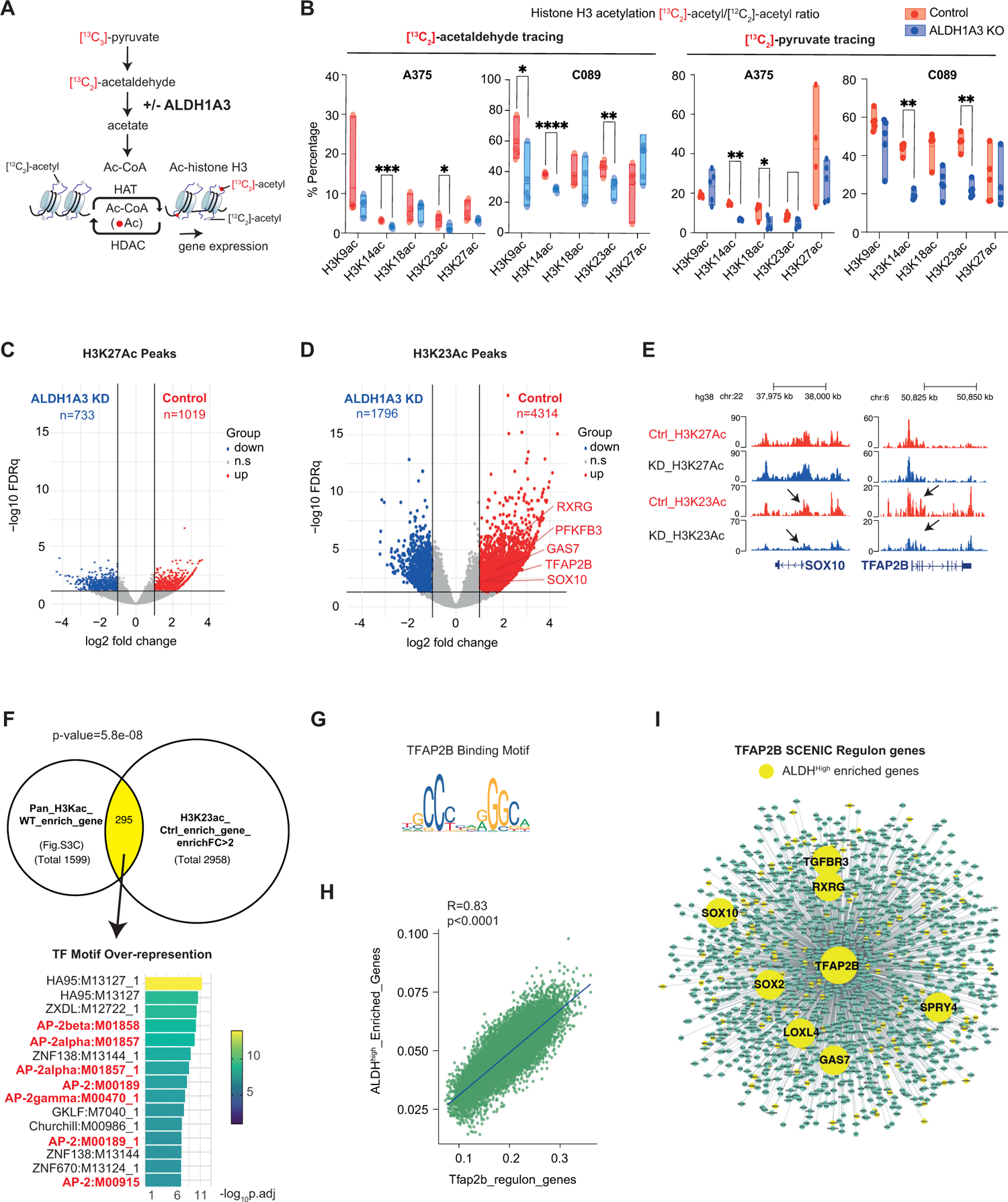
Pyruvate-derived acetaldehyde serves as an acetyl source for histone H3-acetylation. **A.** Schematic summarising the acetyl group transfer from pyruvate to histone mediated by ALDH1A3 metabolising acetaldehyde, with the experiment design that ^13^C_2_ - acetaldehyde and/or ^13^C_3_ pyruvate derived acetyl groups can be traced to histone acetylation. **B.** Pyruvate derived acetaldehyde is an acetyl source across multiple lysine residues on Histone H3. ^13^C_2_-acetyl groups were traced from acetaldehyde into lysine acetylation at K9, K14, K18, K23, and K27, measured by HPLC-MS/MS. *ALDH1A3* knockout cells treated with ^13^C2-acetaldehyde showed less ^13^C_2_-acetyl groups at K9, K14 and K23 residues. Similar differential patterns of ^13^C_2_-acetyl groups incorporation at histone H3 were observed when cells were treated with ^13^C_3_ sodium pyruvate. n=5 per cell line; P values by two-way ANOVA with Sidak’s multiple comparisons test. **C-D.** Volcano plot of differentially acetylated histone H3K27 peaks **(C)** and H3K23 peaks **(D)** from CUT&TAG in MPD002 control and ALDH1A3 KD cells (highlighted by red and blue respectively, fold change >2, FDR.q < 0.05). Representative NCSC gene related peaks are annotated in the control enriched H3K23 enriched peaks. **E.** Gene tracks of acetylated histone H3K27 and H3K23 peaks in MPD002 vehicle control versus *ALDH1A3* KD cells at representative NCSC genes. No significant changes are present in H3K27, while the significant change in H3K23 acetylation is observed. **F.** Venn diagram of overlapping genes between (upper left) genes enriched in A375 control cells versus *ALDH1A3* knockout cells mapped from total histone H3 acetylation and (upper right) MPD002 control cells versus ALDH1A3 knockdown cells mapped from histone H3K23 acetylation. P values by Fisher’s exact test. Transcription factor binding motif over-representation analysis of the Venn diagram overlapping genes (lower panel) showed significant enrichment of AP-2 binding motif, with enrichment score by g:Profiler (version e111_eg58_p18_30541362) with g:SCS multiple testing correction method applying statistical significance threshold of 0.05^111^. **G.** Binding motif of human TFAP2B from JASPAR database. **H.** Scatter plot shows the correlation of ALDH^High^ enriched gene signature and Tfap2b regulon activities (AUCell score) in murine melanoma cells (*NRAS^Q61K/°^;Ink4a^−/−^*)^43^. **I.** SCENIC-inferred Tfap2b regulatory network using a murine mouse scRNA-seq dataset^43^, with Tfap2b target genes in green and ALDH^High^ enriched Tfap2b target genes highlighted in yellow. Tfap2b regulon genes were mapped to human homologs to allow comparison and visualization. See also **Table S5**.

Next, we traced ^13^C_3_-labelled pyruvate to histones, and again consistently detected higher percentage of ^13^C-labelled acetylated histone H3 proteins in cells with high ALDH1A3 activity (**Figure 4B, right panel)**. This observation supports a model in which pyruvate can serve as the source of acetaldehyde and subsequently acetate for acetyl-CoA.

### ALDH1A3 alters selective Histone H3 acetylation in the genome

To better understand the significance of ALDH1A3 dependent acetyl-histone H3, we considered that previous work reported that selective histone acetylation mechanisms can regulate specific gene expression, including glucose metabolism enzymes^60,61^, and that global histone repositories serve as reservoirs for acetyl groups^62^. Extrapolated to our data, we hypothesized that pyruvate-derived acetaldehyde serves as a local source of acetyl-CoA for histone acetylation at specific gene loci to generate a cancer stem cell-like state. Conversely, in the low ALDH1A3 state, acetate derived acetyl-CoA is deposited on chromatin as a reservoir^62^, serving as a rapid source for acetyl-coA and as a buffer for intracellular pH^63,64^.

To determine how ALDH1A3-dependent histone H3 acetylation is deposited on chromatin, we first performed quantitative acetyl-histone H3 chromatin immunoprecipitation (ChIP)-seq using an antibody against pan-histone H3 acetylated sites (K9 + K14 + K18 + K23 + K27) (**Supplementary extended data; Figure S3, S4)**. Notably, in control cells, the enriched acetyl-histone H3 peaks were clustering around transcription start sites (TSSs), especially within 1kb of promoters, whereas in *ALDH1A3* KO cells, the enriched acetyl-histone H3 peaks were broadly dispersed throughout the genome, and particularly spreading into the distal intergenic region and intronic regions **(Figure S3D, E)**. Next, we found that acetyl-histone H3 sites enriched in the ALDH1A3^High^ state were present in NCSC and glucose metabolism pathway genes, while conversely, the loss of *ALDH1A3* was associated with a broadly dispersed acetyl-histone H3 landscape, in which subsets of elevated acetyl-histone H3 peaks were associated with proinflammatory genes **(Figure S4G-H; Table S4)**.

While consistent with the RNA expression patterns that we identified earlier, the use of the pan-histone acetylation antibody that recognizes five different marks could be masking the effect of site selective acetylation events. To address this, we performed CUT&TAG analysis in MPD002 control and *ALDH1A3* KD cells using individual antibodies selective in H3K23ac (a mark dependent on ALDH1A3 from 13C-labelled acetaldehyde and pyruvate tracing) and H3K27ac (a mark not dependent on ALDH1A3 from 13C-labelled acetaldehyde and pyruvate tracing). We found limited differential peaks between WT and ALDH1A3 KD groups for H3K27ac, with much stronger differential effects for H3K23ac (**Figure 4C, D)**. More importantly, by mapping the H3K23ac enriched peaks in the control comparing to ALDH1A3 knockdown cells, we again identified NCSC genes, including TFAP2B and SOX10 (**Figure 4E**). Further, by comparing the total enriched H3 acetylation peak target genes in A375 (control versus *ALDH1A3* KO) with the enriched H3K23Ac peak target genes in MPD002 H3K23 (control versus *ALDH1A3* KD), we found the overlapping target genes are over-represented for TFAP2 transcription factor (AP-2) motif (**Figure 4F, G; Table S5)**.

The discovery of the TFAP2 motif supports a likely TFAP2B feed-forward loop, acting both as a direct target and mediator of ALDH1A3-dependent gene expression, and thereby establishing and sustaining the ALDH1A3^High^ neural crest identity. The feed-forward loop is supported by i) the high similarity of binding motif across all TFAP2 members and the exclusive association of *ALDH1A3-TFAP2B* **(Figure S1J)**, ii), the fact that TFAP2B can bind chromatin regulatory regions upstream of its own gene locus^65^, and iii) our work in zebrafish proposing that Tfap2b regulates a set of genes that defines the melanocyte stem cell, including *tfap2b* itself^46^. To test our prediction, we employed gene regulatory network inference (SCENIC)^66^ and identified a melanoma derived *Tfap2b* regulon **(Table S5)**. When comparing AUCell expression scores of the *Tfap2b* regulon with our ALDH^High^ enriched gene set across 16,700 single melanoma cells, originating from 5 primary mouse murine lesions^43^, we detected significant co-expression of both transcriptional programs (**Figure 4H**). When we intersected Tfap2b regulon genes with ALDH^High^ enriched genes, we found a striking overlap that included both NC and stem cell genes such as *SOX2*, *SOX10*, *SPRY4* and *RXRG* (**Figure 4I, Table S5)**.

### ALDH1A3 forms a predicted complex with ACSS2

Nutrition, cellular metabolism, and transcription are intimately linked with epigenetic control of gene expression as exemplified in yeast where oxidation of ethanol to generate acetyl-CoA is tightly co-ordinated with histone acetylation and expression of selective growth genes^67^. Following our discoveries supporting ALDH1A3-acetaldehyde to selectively regulate targeted histone H3 acetylation, we wanted to further understand the mechanism mediating such selective acetyl-group deposit. When we analysed the proteomic database STRING^68^ to identify protein interactions potentially forming regulatory networks with ALDH1A3, we noted that acetyl-CoA synthetase 2 (Acs2) and pyruvate decarboxylase (Pdc) are interacting partners of aldehyde dehydrogenase (Ald6; ortholog of ALDH1) in yeast (*Saccharomyces cerevisiae*)^69^ (**Figure 5A**). The Ald6 and Acs2 interaction in yeast was intriguing given studies of neuronal stem cell differentiation and memory formation in mammals in which acetyl-CoA synthase (ACSS2) was reported to accumulate in the nucleus and generate acetyl-CoA “on-demand” from chromatin-bound acetate associated with selective histone acetylation and gene expression^70,71^.

**Figure 5:**
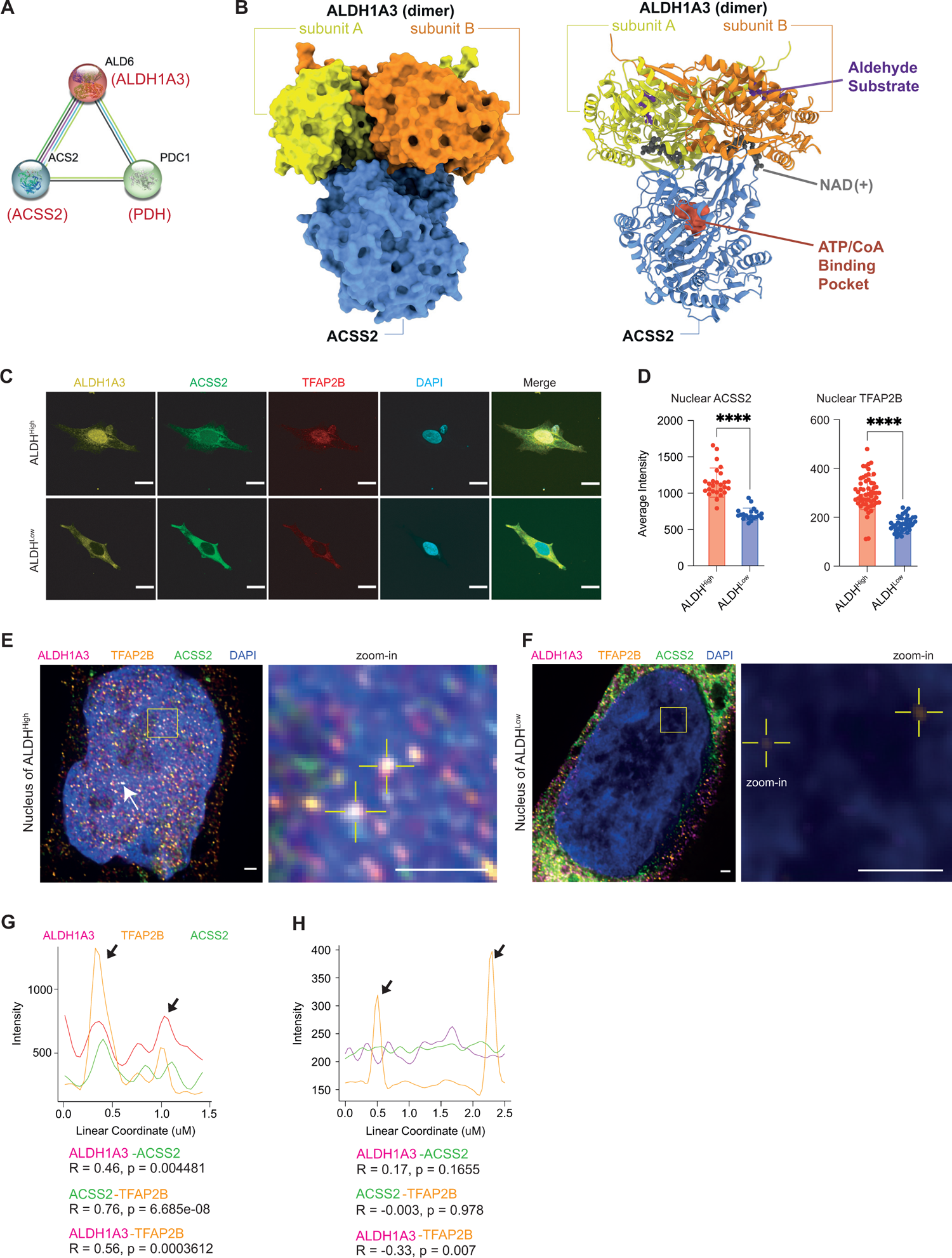
ALDH1A3 forms a predicted complex with ACSS2s. **A.** ALDH1A3 functional association. STRING functional protein association analysis between yeast Ald6, Acs2, and Pdc1, orthologous to human ALDH1A3, ACSS2, and PDH. **B.** AlphaFold multimer modelling of human ALDH1A3 and ACSS2. Proteins visualised as a complex in surface model (left) and ribbon model (right). The binding sites of an aldehyde substrate (retinaldehyde) and co-factor (NAD+) were created by structural alignment of the AlphaFold-Monomer predicted structure to PDB ID 5fhz, a published tetramer ALDH1A3 structure. The ATP/CoA binding pocket of ACSS2 was visualised by structural alignment of AlphaFill-optimized protein-ligand complexes (ATP donor: PDB ID 5k8f; CoA donor: PDB ID 3gpc). **C-D.** Subcellular expression of ALDH1A3, ACSS2, and TFAP2B. **(C)** ICC staining by fluorescence antibody probing ALDH1A3 (yellow), ACSS2 (green), TFAP2B (red) in sorted A375 ALDH^High^ and ALDH^Low^ cells. DAPI (blue). Scale bar = 10 μm. **(D)** Fluorescence signal intensity quantification of nuclear TFAP2B and ACSS2 in ICC images. N=2 biological repeat experiments, n = 55 quantified single cells for ALDH^High^ TFAP2B, n= 46 single cells for ALDH^Low^ TFAP2B; n = 26 single cells for ALDH^High^ ACSS2, n= 20 single cells for ALDH^Low^ ACSS2, (represented as individual dots), mean±s.d., unpaired non-parametric Kolmogorov-Smirnov test. ****P<0.0001. **E-H.** ALDH1A3, ACSS2 and TFAP2B co-localise in the nucleus. Structured illumination microscopy (SIM) of ALDH1A3 (magenta), TFAP2B (orange), ACSS2 (green), and DAPI (blue) in A375 melanoma FACS sorted **(E)** ALDH^High^ cells and **(F)** ALDH^Low^ cells. Scale bar = 1 μm. Complex co-localization signals were abundant (arrows) and are indicated as highlighted dots in the zoomed image in **E**. Low signals of nuclear TFAP2B are highlighted in zoomed image in **F**. **(G)** Intensity plot profiles of the line scan across two co-localisation hotspots in zoomed image of **E**. Signal overlap peaks are indicated by arrows. **(H)** Intensity plot profile of the line scan across the two TFAP2B signal spots in zoomed image of **F**. TFAP2B signal peaks are indicated by arrows, where no ALDH1A3 or ACSS2 signals are present.

Intrigued by these relationships, we used AlphaFold-Multimer, an extension of AlphaFold2 designed to predict protein-protein interactions^72^, to predict interactions between yeast Acs2 and Ald6 and between their human homologues ACSS2 and ALDH1A3 (**Figure 5B, Figure S5A)**. This approach yielded complexes for both the yeast and human pairs with moderate confidence (45% and 47%, respectively). In addition, we identified strong interface conservation between Acs2 and ACSS2 (Pearson correlation of 0.61 between buried surface area values of homologous residues, p = 1.7 × 10^-66^). In these models, Acs2/ACSS2 binds at the perimeter of the Ald6/ALDH1A3 dimerization interface in a 1:2 stoichiometry. This binding mode ensures accessibility of the Acs2/ACSS2 binding pockets for CoA and ATP and the Ald6/ALDH1A3 substrate pockets for NAD and aldehyde. Thus, both the predicted yeast and human complexes are consistent with an ACSS2-ALDH1A3 enzymatic partnership in the nucleus.

When we performed immunofluorescence on sorted A375 ALDH1A3^High^ and ALDH1A3^Low^ melanoma cells, we found that ACSS2, along with ALDH1A3 and TFAP2B, was localized in the nucleus of sorted ALDH1A3^High^ cells but not in ALDH1A3^Low^ cells (**Figure 5C, D)**. In the presence of an ACSS2 inhibitor, we found no change in nuclear ALDH1A3 whereas nuclear TFAP2B levels were significantly reduced **(Figure S5B-C)**. We propose that the ALDH1A3-ACCS2 enzymatic partnership is not required for ALDH1A3 nuclear localisation, however we suggest ALDH1A3 - dependent TFAP2B expression and nuclear localisation is mediated through ACSS2.

To investigate potential co-localisation of ACSS2 and ALDH1A3 in cells, we sorted ALDH^High^ and ALDH^Low^ cells and performed super-resolution imaging (**Figure 5E, F)**. We detected abundant nuclear co-localization signals of ALDH1A3-ACSS2-TFAP2B and a significant positive correlation between the linear signal distribution of any two of the three target proteins (**Figure 5G**). In contrast, we saw minimal nuclear signals of ALDH1A3, ACSS2, or TFAP2B in sorted ALDH^Low^ cells and the nuclear signals captured by super-resolution imaging were not overlapping (**Figure 5H**). Finally, we tested the potential for interaction *in vivo* by co-immunoprecipitation and confirmed the interaction of ACSS2 with ALDH1A3 in both A375 whole cell and nuclear department **(Figure S5D)**.

### ALDH1A3 determines ACSS2 binding to NCSC gene loci

To identify genomic loci associated with ACSS2 dependent acetylated histone peaks and increased gene expression, we designed primers targeting the promoter regions of NCSC genes and IRF1 genes and performed ACSS2 ChIP-qPCR in both A375 and MAPKi-resistant MPD002 cells (**Figure 6A**). ACSS2 ChIP-qPCR revealed higher levels of ACSS2 binding to multiple NC gene promoter regions in control cells comparing to *ALDH1A3* KO/KD cells, while in contrast ACSS2 binding was enriched at the *IRF1* promoter in *ALDH1A3* KO/KD cells. This effect was restored upon *ALDH1A3* overexpression in A375 cells. This data shows that ALDH1A3 is required for ACSS2 binding to specific loci to shape the genomic landscape.

**Figure 6:**
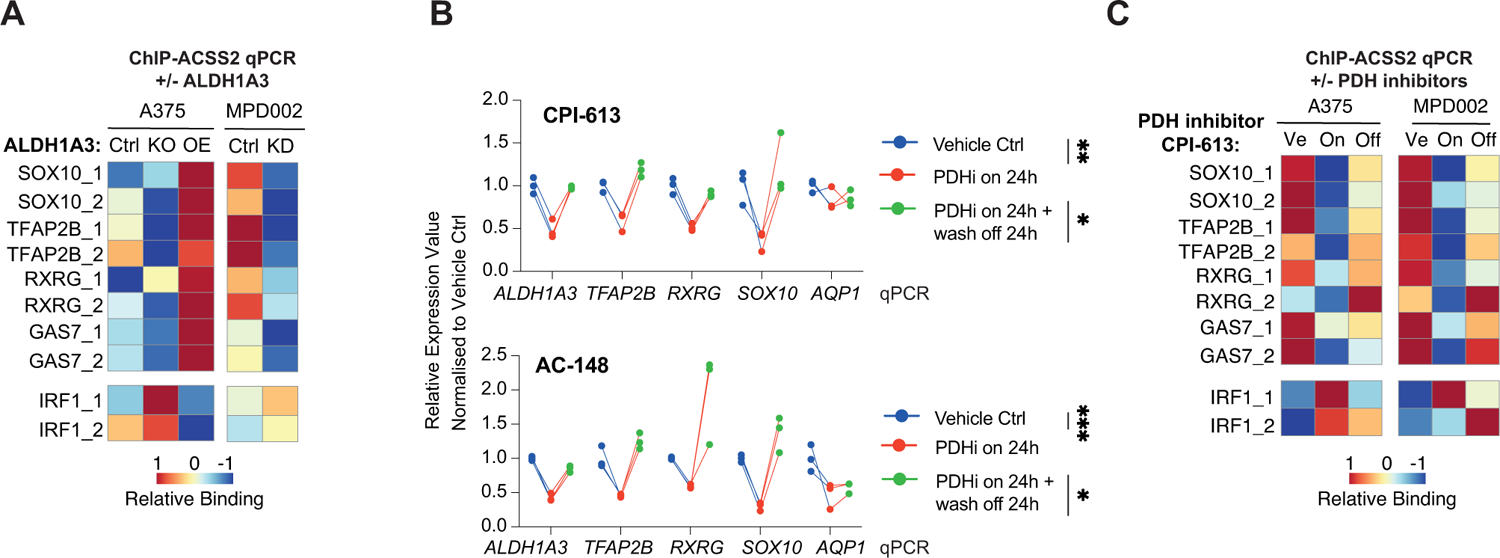
Acetaldehyde serves as an acetyl source for histone H3-acetylation. **A.** ALDH1A3 directs gene-specific ACSS2 chromatin binding. ACSS2 ChIP-qPCR results in (left panel) A375 WT control versus *ALDH1A3* KO and *ALDH1A3* OE cells, as well as in (right panel) MPD002 control versus *ALDH1A3* KD cells (siALDH1A3). N=3 biological repeat experiments in both A375 and MPD002 cells; each with n=3 technical replicates and normalised to IgG control before heatmap scaling and plotting. **B.** RT-qPCR measured NCSC gene expression change in A375 and MPD002 in response to pyruvate dehydrogenase (PDH) activity change induced by PDH inhibitors (PDHi) CPI-613 (upper panel) and AC-148 (lower panel). *P<0.05; **P<0.01; ***P<0.001; One-way ANOVA with Sidak’s correction. **C.** ALDH1A3 directs gene-specific ACSS2 chromatin binding relies on PDH activity. ACSS2 ChIP-qPCR results in A375 and MPD002 cells with vehicle control (Ve) versus PDH inhibitor CPI-613 treated samples (On, 24h) and PDH inhibitor wash off samples (24h treatment followed by additional 24h wash off recovery, Off). N=2 biological repeat experiments in both A375 and MPD002 cells; each with n=3 technical replicates and normalised to IgG control before heatmap scaling and plotting.

### PDH inhibitors alter ACSS2 binding to selective genomic loci

To strengthen our understanding of the role of pyruvate in the ALDH1A3-ACSS2 mechanism, we used two independent PDH inhibitors, the thiamine analogue AC-148^73^ **(Supplementary extended data)** and a commercially available lipoate related PDH inhibitor, CPI-613^74^, and determined the effect on gene expression of NCSC genes. Addition of either of the PDH inhibitors for 24 hr led to decreased expression in NCSC genes, followed by restored NCSC gene expression (except *AQP1*) 24hr following wash-off (**Figure 6B**). Next, we addressed if PDH inhibition altered the ACSS2 binding to multiple neural crest gene loci by ChIP-qPCR, in both A375 and MPD002 cells. We found that ACSS2 binding to these regions in control cells was reduced with PDHi treatment and restored upon inhibitor washout (**Figure 6C**).

Based on our findings thus far, we propose a model in which ALDH1A3 and ACSS2 cooperate in a metabolic cascade from pyruvate to acetaldehyde to generate a regional chromatin source of acetyl-CoA from nuclear acetaldehyde. The acetyl-CoA source is used to selectively deposit localised histone H3 acetylation and is associated with increased expression of NCSC genes (including *TFAP2B* itself) and glucose metabolism genes.

### ALDH1A3 cells promote drug resistance and disease recurrence *in vivo*

We next sought to evaluate functional relationships between ALDH1A3^High^ cells, melanoma disease progression and therapy resistance. To this end, we employed the widely used BRAF^V600E^ p53 mutant zebrafish melanoma model, in which BRAF inhibitors initially generate a response that is followed by drug resistance and recurrent melanoma growth^75–77^. Here, we dissected melanomas from BRAF^V600E^ p53 mutant zebrafish that also express GFP in the melanocyte lineage *(mitfa:GFP)*, and performed AldeRed analysis (which is similar to the Aldefluor assay but uses a red fluorescent substrate) (**Figure 7A-B**). Through FACS analysis, we identified a distinct zebrafish cell population with ALDH^High^ activity that was also low for *mitfa:GFP* signal (**Figure 7B**), indicating that both zebrafish and human melanoma ALDH^High^ activity cells express low levels of *MITF*. When we sorted zebrafish melanoma cells with the highest and lowest ALDH activity and compared RNA expression of key marker genes (**Figure 7C-D**), we found that *aldh1a3* was the most enriched *ALDH* family isoform in the ALDH^High^ cells **(Figure S6A)**, in agreement with our findings in human melanoma. Furthermore, in zebrafish, melanoma cells with ALDH^High^ activity were enriched with *aldh1a3, tfap2b, sox2,* and *sox10* while melanoma cells with ALDH^Low^ activity expressed *irf1b* (**Figure 7D**).

**Figure 7.**
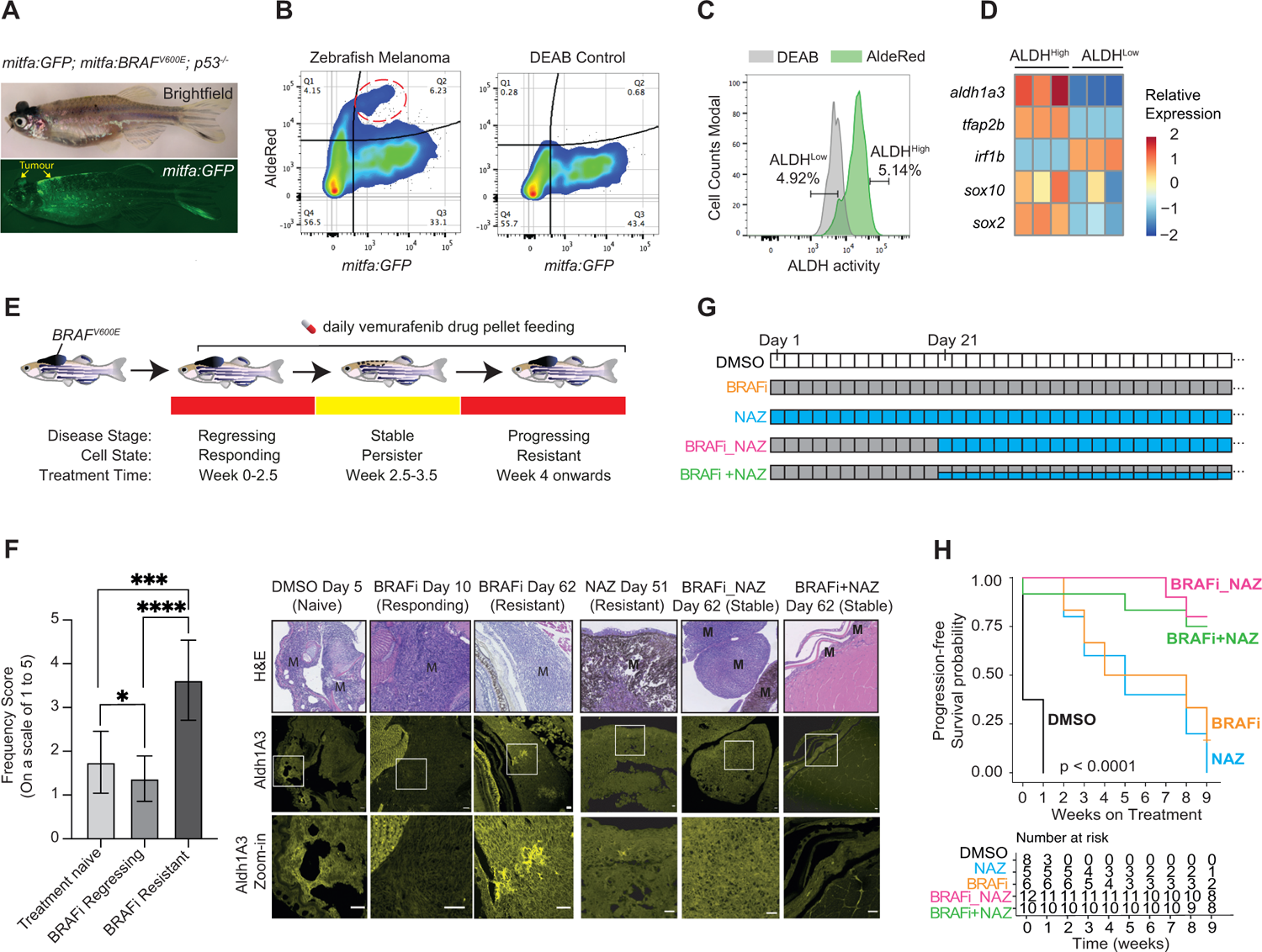
ALDH1A3^High^ subpopulations promote melanoma drug resistance *in vivo*. **A.** Zebrafish BRAF^V600E^ melanoma model. Mesoscopic images of a zebrafish with melanoma (arrow highlighted). *BRAF^V600E^* and *GFP* are expressed in the melanocyte lineage, by the *mitfa* promoter, and the zebrafish is mutant for p53. **B-D.** ALDH^High^ cells in zebrafish melanoma. Dissected zebrafish melanoma tissues were dissociated into single cell suspensions and processed with the AldeRed assay. A cell population with high ALDH activity and correspondingly low levels of *mitfa:GFP* is indicated (red circle). DEAB is an ALDH inhibitor and serves as a negative control. (**C**) Flow cytometry sorting of ALDH^High^ and ALDH^Low^ zebrafish melanoma cells. (**D**) RT-qPCR quantification showing significantly higher *sox2, sox10 and tfap2b* expression in sorted ALDH^High^ versus ALDH^Low^ zebrafish melanoma cells. (n=3 bio-replicates each with 3 technical replicates, Multiple paired t-test corrected with Holm-Sidak’s method). **E-F.** Aldh1a3 in zebrafish model of BRAF-inhibitor regression and recurrent disease. **(E)** Schematic of *BRAF^V600E^ p53* mutant zebrafish with melanomas were non-invasively fed with 200 mg/kg/day vemurafenib-containing food pellets leads to melanoma regression, drug resistance and disease recurrence. **(F)** Aldh1a3 immunohistochemistry in treatment naïve (DMSO), BRAFi and/or NAZ treatment responding disease, and in drug-resistant recurrent disease. Aldh1a3 IHC in melanoma recurrent disease treated with indicated NAZ alone or in combination with BRAFi shows Aldh1a3 on-target efficacy of nifuroxazide. One-way ANOVA with Tukey’s test correction for multiple comparisons. *P<0.05; ***P<0.001; ****P<0.0001. **G-H.** Zebrafish combination drug trial to target ALDH^High^ cells. **(G)** Long-term drug pellet treatment design. **(H)** Kaplan Meier survival curves of zebrafish melanomas under different drug treatment shown in **G**: DMSO (Black): daily DMSO control treatment. BRAFi (Orange): 200 mg/kg/day vemurafenib treatment. NAZ (Blue): daily 150 mg/kg/day Nifuroxazide treatment. BRAFi_NAZ (Magenta): 3-week treatment of 200 mg/kg/day vemurafenib followed by daily 150 mg/kg/day Nifuroxazide treatment. BRAFi+NAZ (Green): 3-week treatment of 200 mg/kg/day vemurafenib followed by the combination of 200 mg/kg/day vemurafenib and 150 mg/kg/day Nifuroxazide treatment. The weekly tumour size change of each fish was calculated and an event of tumour size increase over 20% is considered as the end of progression free survival period. Log-rank tests, p < 0.001.

Next, we administered vemurafenib (BRAF inhibitor) drug pellets to adult zebrafish with melanomas^77^, and upon assessing Aldh1a3 expression by IHC through disease stages (melanoma responding, stable, and progressive disease stages) (**Figure 7E, F)**, we found that clusters of Aldh1a3 cell populations increased in drug-resistant and progressing disease. Together, these data indicate that the zebrafish melanoma model faithfully models the transcriptional states of human ALDH^High^ melanoma cells and the therapy-associated increase in levels of ALDH1A3 identified in **Figures 1 and S1.**

Finally, we designed a zebrafish drug-trial to test whether killing the Aldh1a3^High^ cells that emerge during BRAF inhibitor (BRAFi) drug resistance would impact upon disease outcome (**Figure 7F-G)**. We made use of our drug pellet method to achieve consistent dosing in zebrafish^77^ and administered nifuroxazide (NAZ), a 5-nitrofuran pro-drug that specifically kills ALDH1^High^ cells in human cell lines and mouse xenograft melanoma models^39^. As expected, zebrafish melanoma responded rapidly to BRAFi treatment, but then grew back quickly despite ongoing treatment **(Figure S6B)**. When we administered NAZ treatment alone, we observed a period of stable disease followed by progressive melanoma growth **(Figure S6B)**.

We then tested the two drugs in combinations, using the BRAF inhibitors to target the bulk of the tumour and NAZ to target ALDH1^High^ cells: (1) BRAFi followed by NAZ (BRAFi_NAZ) at day 21, and (2) BRAFi treatment alone until day 21, followed by both BRAFi + NAZ (**Figure 7G**). Our data demonstrate that both combinations of BRAFi plus NAZ significantly improved disease outcomes and probability of survival (**Figure 7H, Figure S6B)**. To investigate whether the improved outcomes are achieved via on-target effect of NAZ, we examined Aldh1a3 levels in NAZ-treated zebrafish melanomas in progressive, drug resistant disease and found that Aldh1a3 clusters were absent (**Figure 7F**). Together, these results support a combination therapeutic strategy using BRAF inhibitors to target the bulk of the tumour and NAZ to eradicate the ALDH1A3-melanoma stem cell pool in the residual disease.

## Discussion

Cell state heterogeneity and plasticity endow genetically identical cancer cells with the capacity to respond differently to treatment. As a result, diverse resistant cell states emerge in patients, challenging clinical strategies that target limited static states^78^. Identifying actionable core regulators of cell heterogeneity and plasticity programmes, through which multi-dimensional information and fluctuating signalling pathways converge to permit cell state dynamics, is therefore a crucial challenge of high clinical relevance^79^. Here, we uncover that nuclear ALDH1A3, functioning as a master coordinator of metabolic and transcriptional cell states, fosters stem-like gene expression programs in melanoma by using pyruvate-derived acetaldehyde as an acetyl donor for histones.

We refer to this coordination of metabolism and developmental lineages in melanoma as a metabolism-neural crest stem cell state axis, based on the present work and evidence from developmental biology suggesting they may be more intimately linked than previously appreciated. In embryonic mouse development, the delaminating neural crest undergoes metabolic reprogramming towards aerobic glycolysis (the Warburg effect) through a mechanism known to initiate expression of transcription factors to promote the epithelial-to-mesenchymal transition (EMT)^45^. More recent evidence in the avian neural crest supports that enhanced glycolysis, mitochondrial respiration, and the pentose phosphate pathway jointly mobilize to achieve coordination of EMT and migration^80^. At later stages, during melanocyte specification, the melanocyte (hair follicle) stem cell transcription factor Yin Yang 1 (YY1) couples MITF with metabolism gene expression and is essential for melanoma development^81,82^. However, we lack an understanding of whether these metabolic states are relevant to neural crest and stem cell states in melanoma and how they crosstalk to induce melanoma drug resistance and disease recurrence.

We identified a nuclear ALDH1A3-ACSS2 mechanism that controls both a glycolytic gene expression program and a developmental NCSC program mediated by the neural crest transcriptional factor TFAP2B. Mechanistically, we provide evidence to support a model whereby nuclear ALDH1A3-ACSS2 directs a glucose-derived pyruvate-acetaldehyde-acetate-acetyl-CoA flux and selectively deposits histone H3 acetylation, including H3K23ac, at genomic loci encoding for glycolysis and TFAP2B-NCSC genes. Cells with low ALDH1A3 preferentially take up acetate to generate a nuclear acetyl-CoA flux, which activates a MITF and IRF1 gene program, but is otherwise broadly deposited on the chromatin as an acetate reservoir. To help visualize these concepts, we present a detailed schematic of this model in **Figure S7A**.

Critically, we demonstrate that endogenous acetaldehyde is a metabolite utilized by ALDH1A3 to modify chromatin. This functional relationship expands the scope of acetaldehyde metabolism from solely protecting cells from aldehyde macromolecule adducts^83,84^. Endogenous and alcohol-derived acetaldehydes, as well as other aldehydes including formaldehyde, are highly reactive towards DNA, and cells employ a robust two-tier protection mechanism to protect against aldehyde-induced DNA damage^83,85–88^. Tier 1 involves aldehyde detoxification enzymes (ALDH2, ADH5) followed by Tier 2 Fanconi anaemia DNA damage repair pathways to repair aldehyde-induced DNA damage^84^. Gene variants in these pathways in people lead to a loss of protection and directly contribute to increased cancer risks, bone marrow failure and risks associated with the alcohol-exposed developing foetus^89–94^. However, recent evidence also shows exogenous alcohol can serve as an acetyl source for histones in the brain via ACSS2, near genes involved in learning and memory, and in the liver^95,96^. Although it is not known if alcohol-derived acetylated histones are generated via local ALDH enzymes and acetaldehyde, combined with our results, these findings point to a possible broader role for acetaldehyde as a metabolite source for chromatin regulatory marks. Supporting this intimate relationship between aldehydes and chromatin modification, recent evidence shows that oxygenase mediated nucleosome demethylation releases formaldehyde^97^, and that histone deacetylase 3 suppresses endogenous formaldehyde production, the reaction products of which are used in one-carbon (formate) metabolism^98,99^. Thus, chromatin may be both modified by aldehydes to shape the regulatory landscape, as well as serve as a source of metabolites when required, including Acetyl-CoA and formate.

Our data here, along with recent discoveries that formaldehyde metabolism promotes differentiation of primed melanocyte stem cells^100^, highlights an emerging physiological role for aldehydes as essential metabolites governing stem cell function. Pioneering studies of nuclear condensates show that the spatial clustering of active gene transcription is intimately linked with nuclear protein-metabolite condensate distribution^101^, providing a possibility that subnuclear ALDH-aldehyde metabolism may exert local control of gene expression or acetate reservoirs. It will be important for future studies to identify the external signals that promote the ALDH1A3 ^High^ state following drug treatment, and address whether these contribute to the cross-resistance mechanisms triggered by targeted therapies and immune checkpoint blockade. Transcriptionally, analysis of two independent HA-tagged MITF ChIP-seq experiments show HA-MITF binds the ALDH1A3 promoter region^102,103^, and given that we find MITF activity is low in ALDH1A3^High^ cells, this suggests that MITF may be a repressor of ALDH1A3 transcription **(Figure S7B)**. While an increase in pyruvate accounts for the source of acetaldehyde in ALDH1A3^High^ cells in the models presented here, it will be important to identify if aldehydes available from the microenvironment (possibly even through diet or the microbiome) can impact upon on the metabolism-neural crest stem cell state axis in melanoma.

In conclusion, we present evidence that ALDH1A3 is a master regulator of a metabolism-NCSC state axis that partners with ACSS2 to modify histone acetylation by locally depositing acetaldehyde-derived acetyl groups. In cancer cell lines and low passage patient-derived melanoma cells, we show that ALDH1A3 expression alone is sufficient to shift cells towards NCSC states. Further, we show that targeting ALDH1A3-NCSC cells *in vivo* is a relevant therapeutic strategy given evidence from our pre-clinical melanoma models showing that ALDH1A3^High^ states emerge during BRAF inhibitor resistance, and that eliminating these cells extends disease-free survival. The conceptual framework we present here for melanoma may be broadly applicable to ALDH^High^ cancer stem cell subpopulations in other cancer types, as ALDH isoforms potentially cooperate with lineage specific master transcription factors (often from developmental lineages) that are co-opted to regulate tumour cell states^104^.

### Limitations of this study

Here, we show that ALDH1A3 is a regulator of H3K23 acetylation at NCSC genes, with little effect on H3K27. Additional CUT&TAG or ChIP-Seq experiments using independent antibodies for each of the H3K-ac marks would clarify how acetaldehyde-selective histone H3 acetylation relates to the chromatin landscape beyond H3K23 and H3K27. Testing for ACSS2 interactions with other ALDH family members would provide more insight into the specificity and wider applicability of ACSS2-ALDH interactions in other biological contexts. We use the ALDH1 suicide inhibitor pro-drug NAZ to kill ALDH1A3 cells in zebrafish models of MAPKi-treated melanoma and show on-target specificity using IHC. However, drugs often have more than a single target in vivo, and we are not able to determine the contribution of these potential other targets to the melanoma response in our experimental model. An inducible genetic model for targeted cell ablation ^105^ of *aldh1a3* expressing cells could provide supporting evidence regarding the killing ALDH1A3 cells in combination with MAPKi therapies to delay or prevent melanoma recurrence.

## STAR METHODS

### KEY RESOURCES TABLE

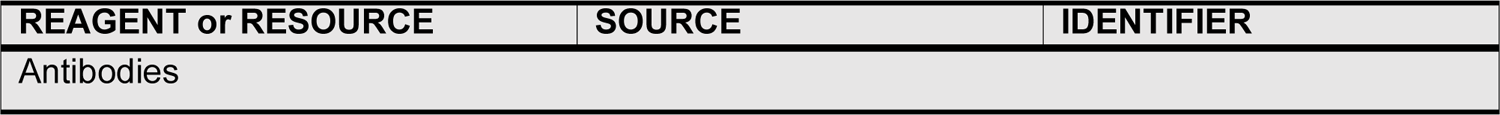
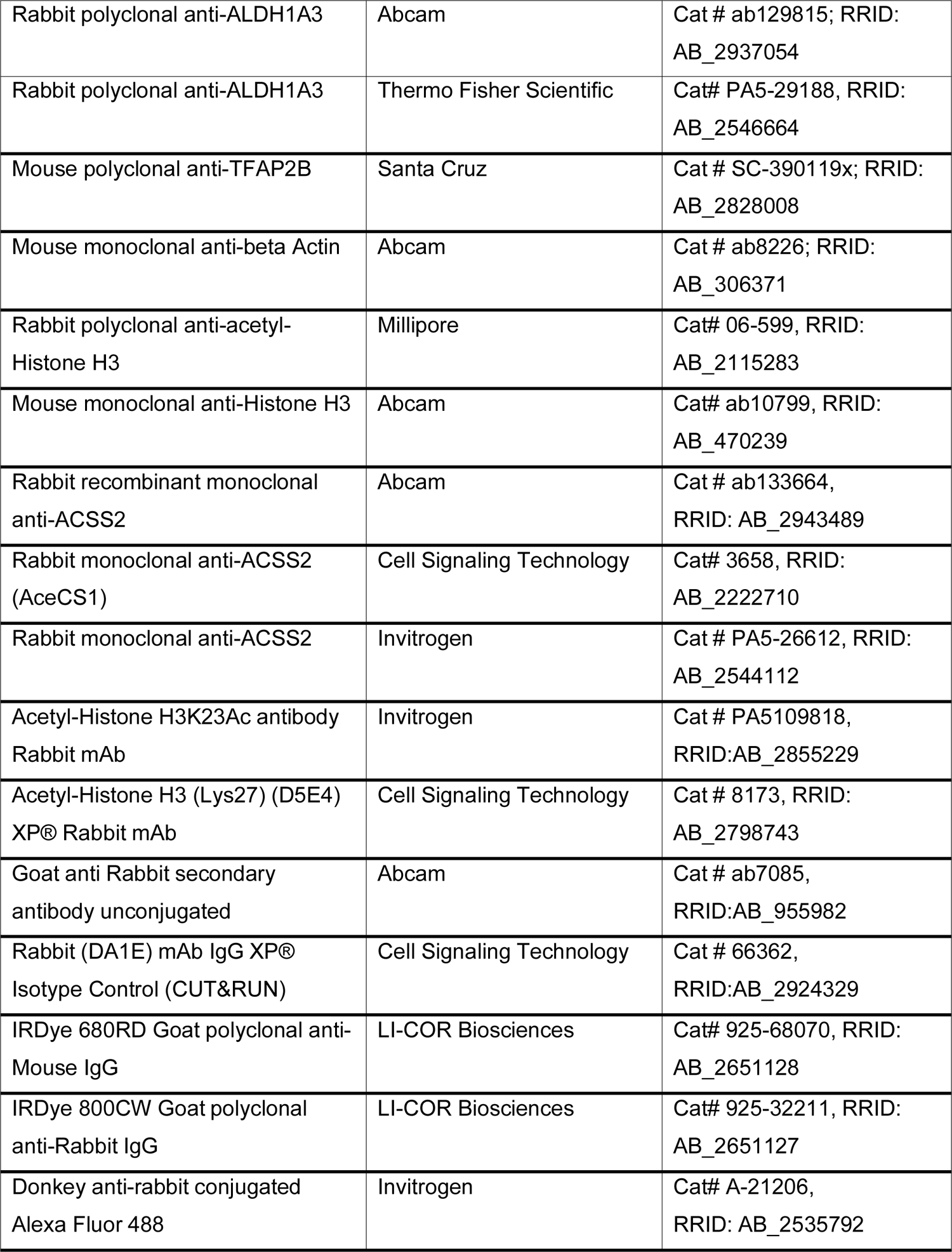
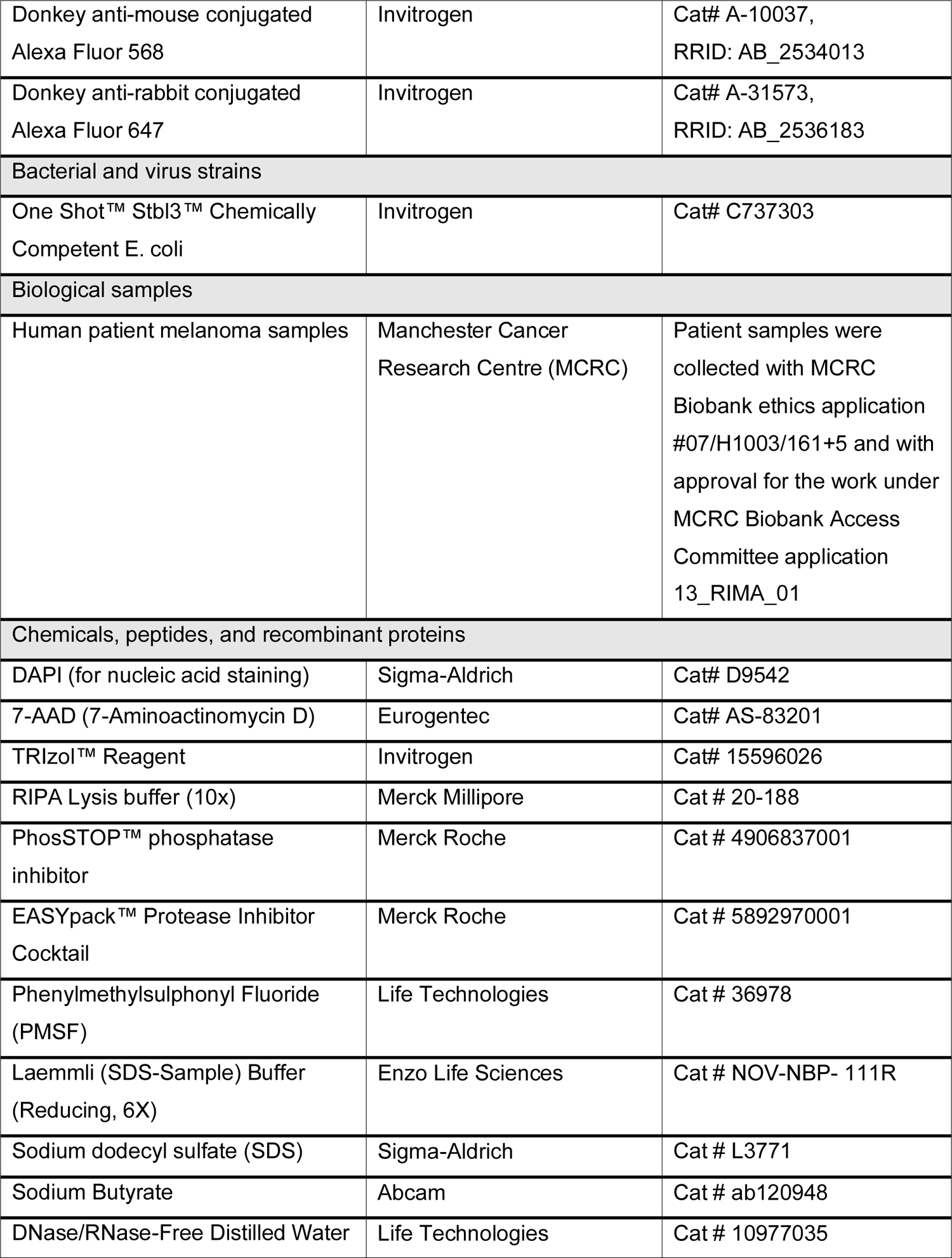
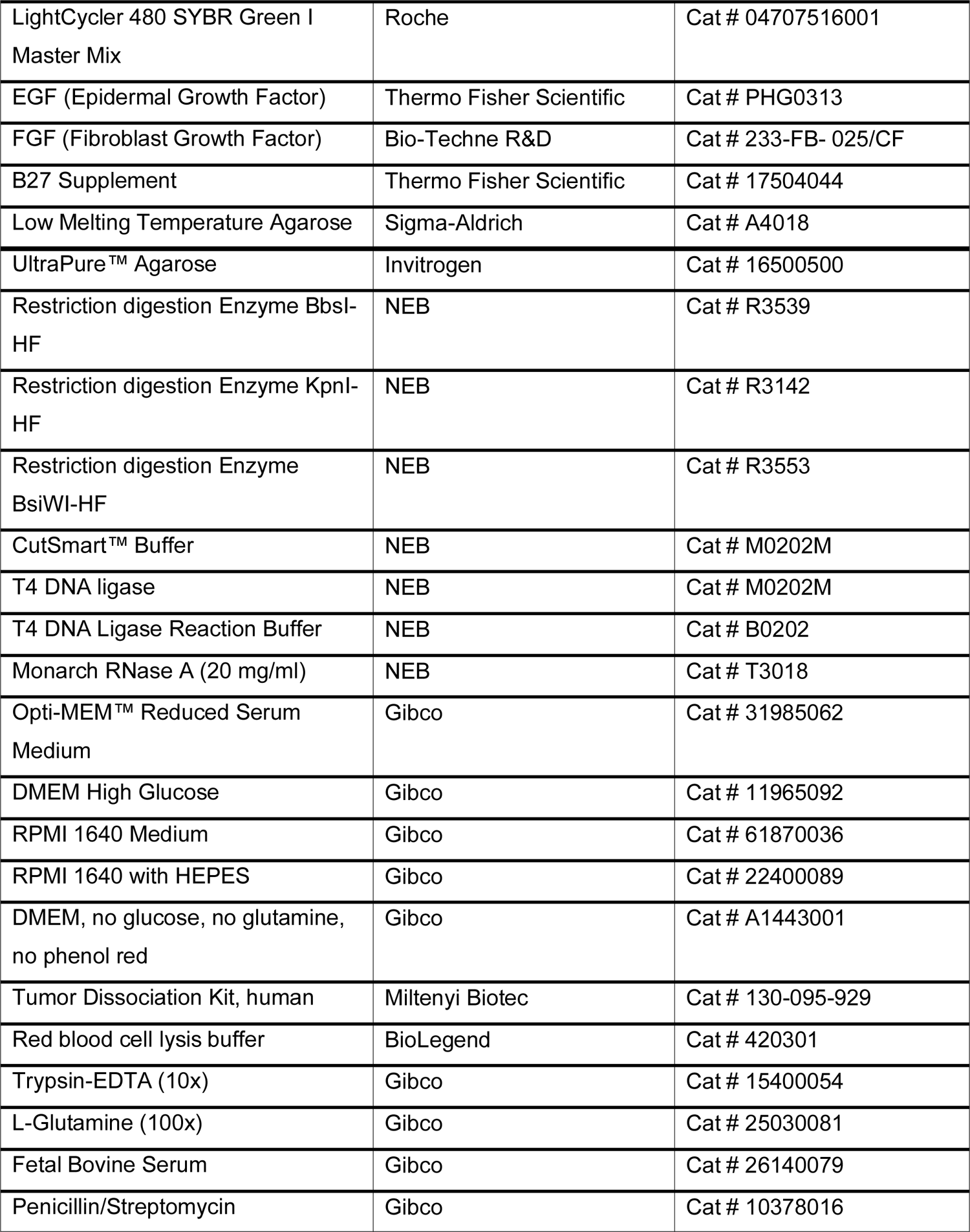
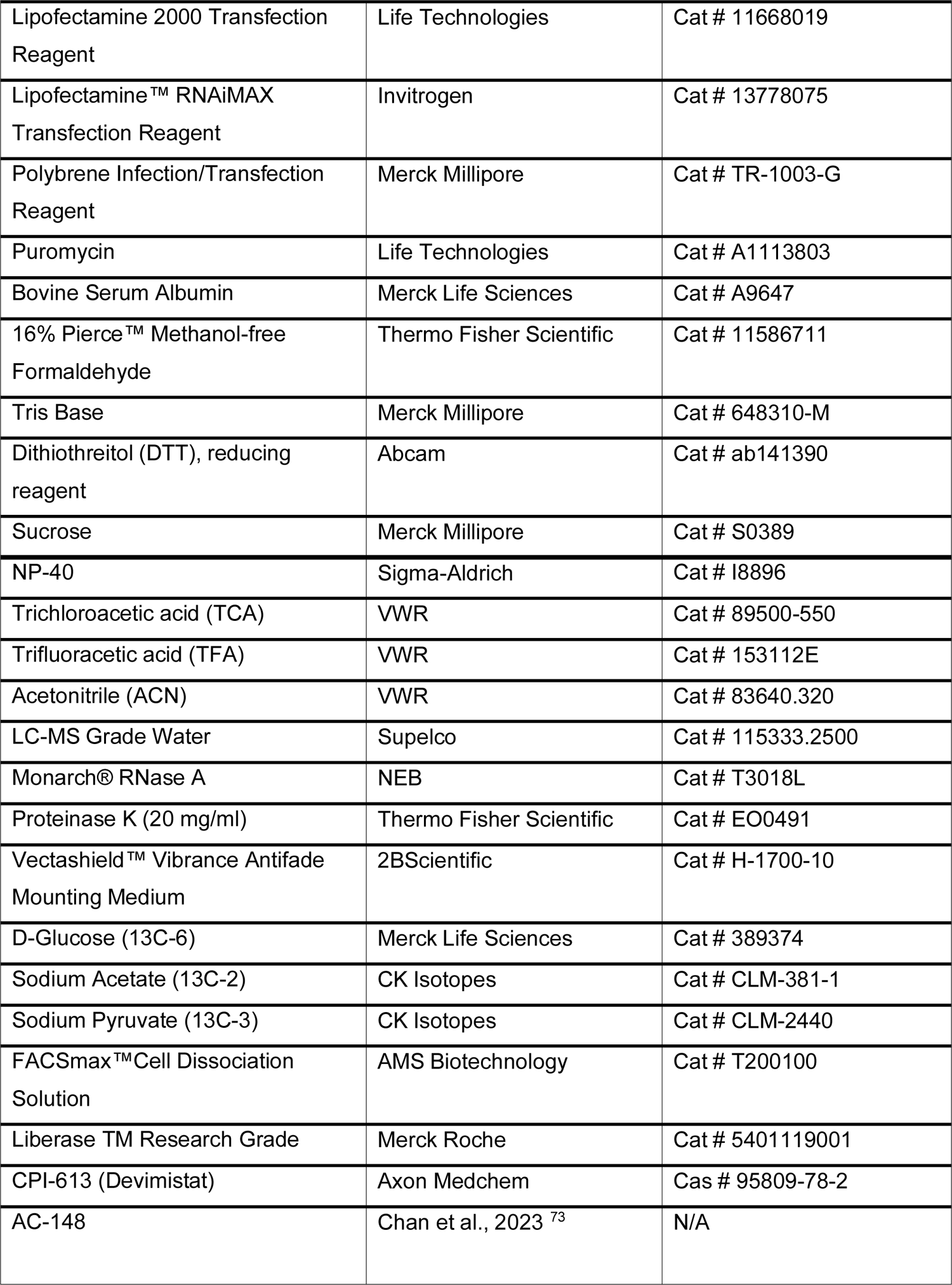
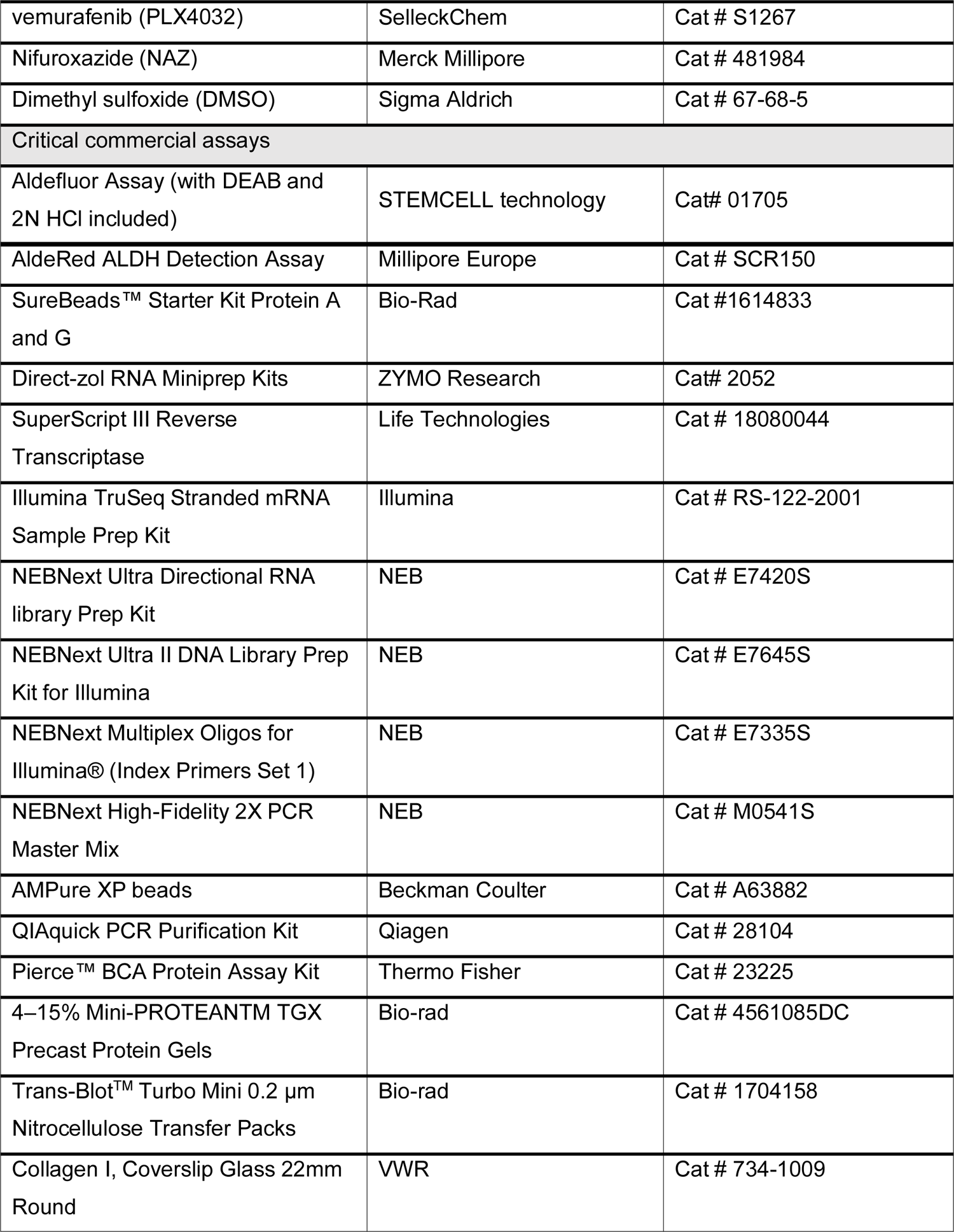
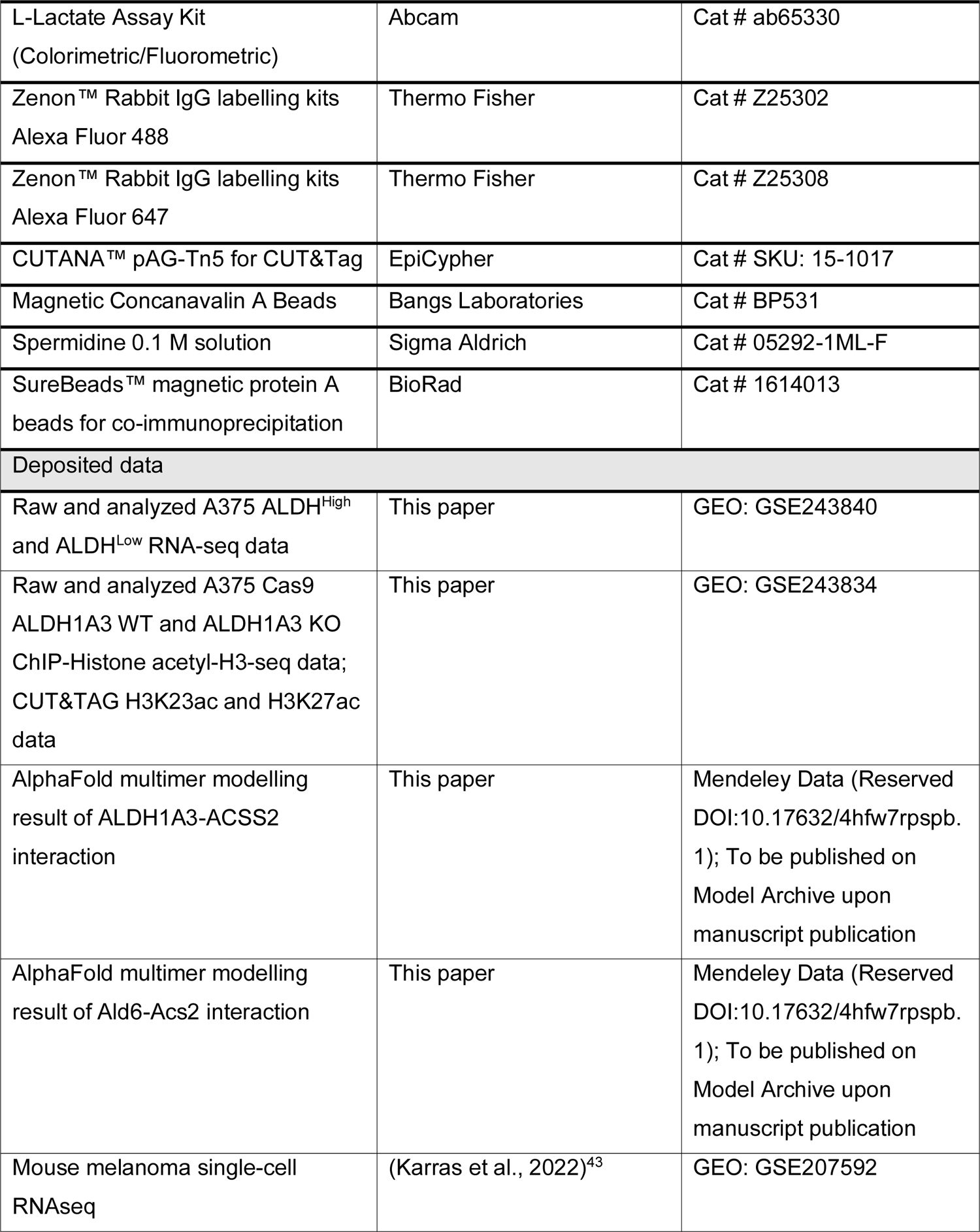
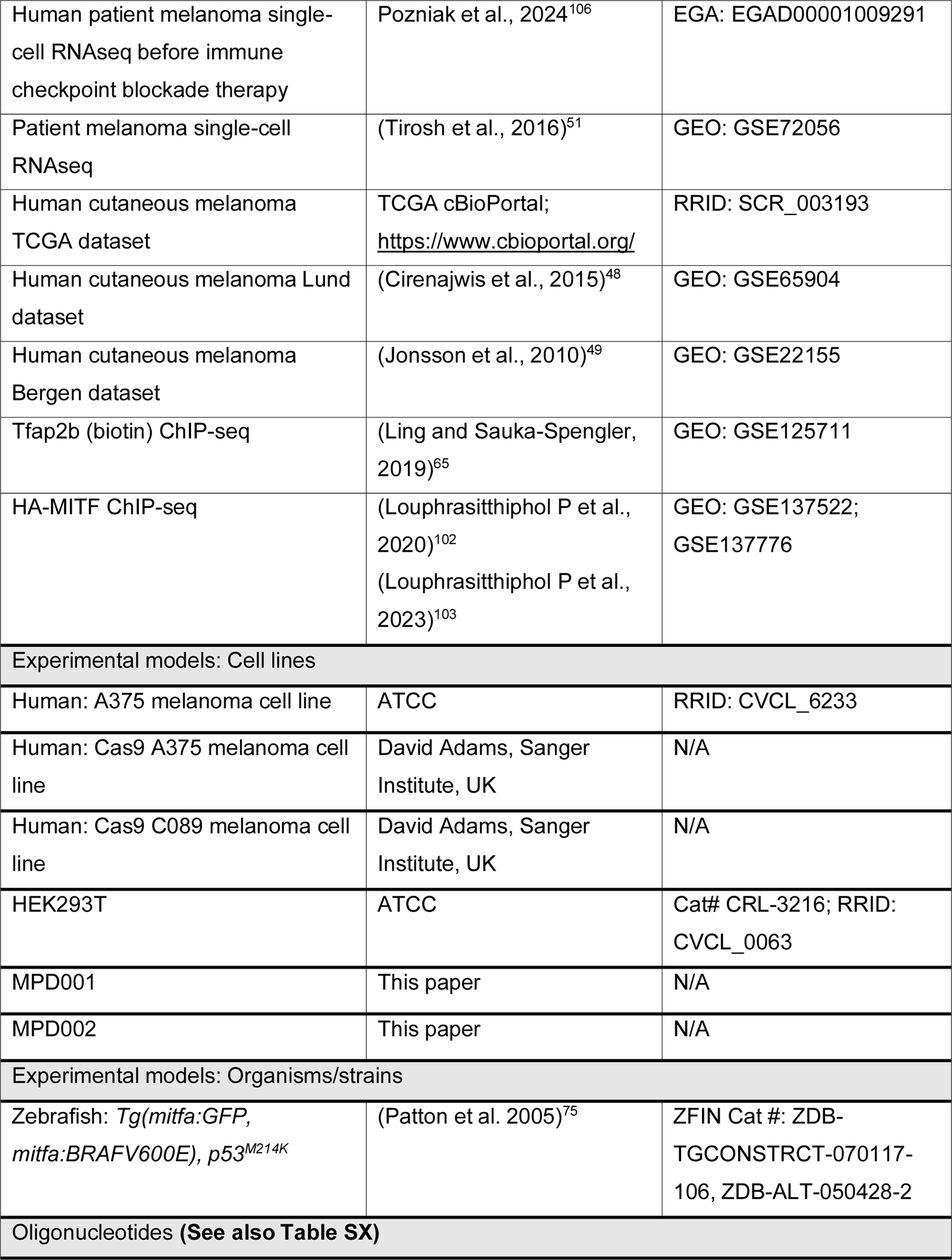
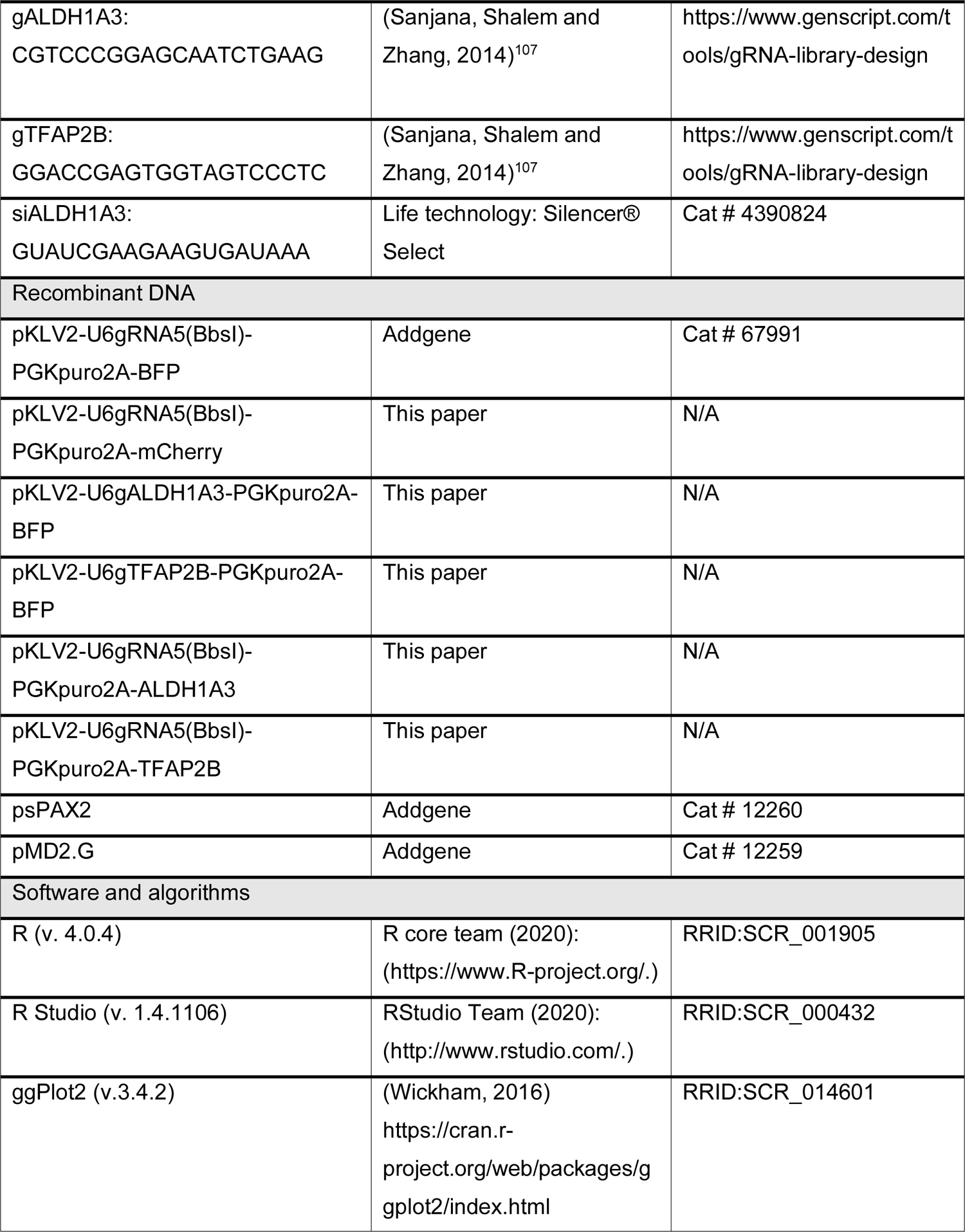
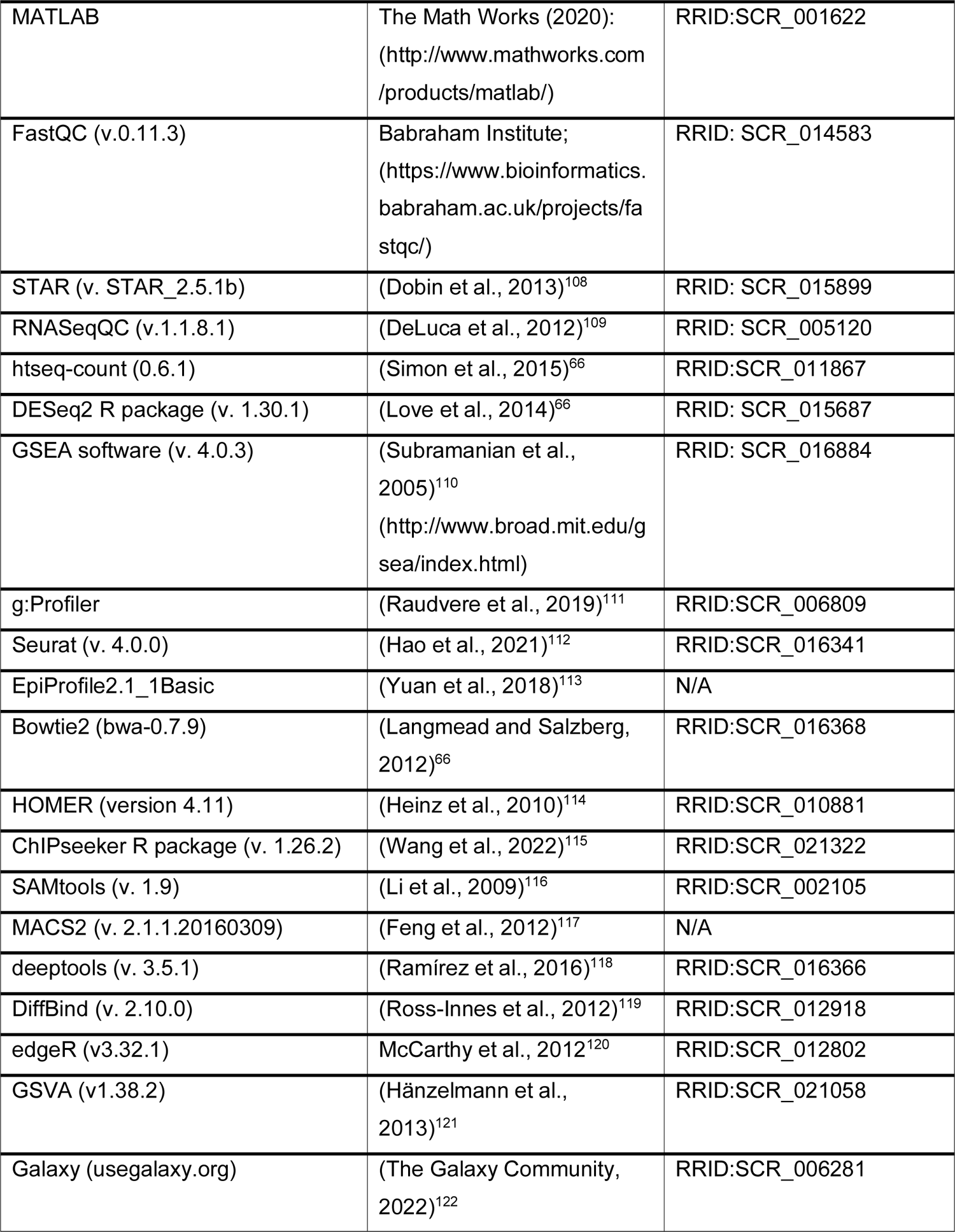
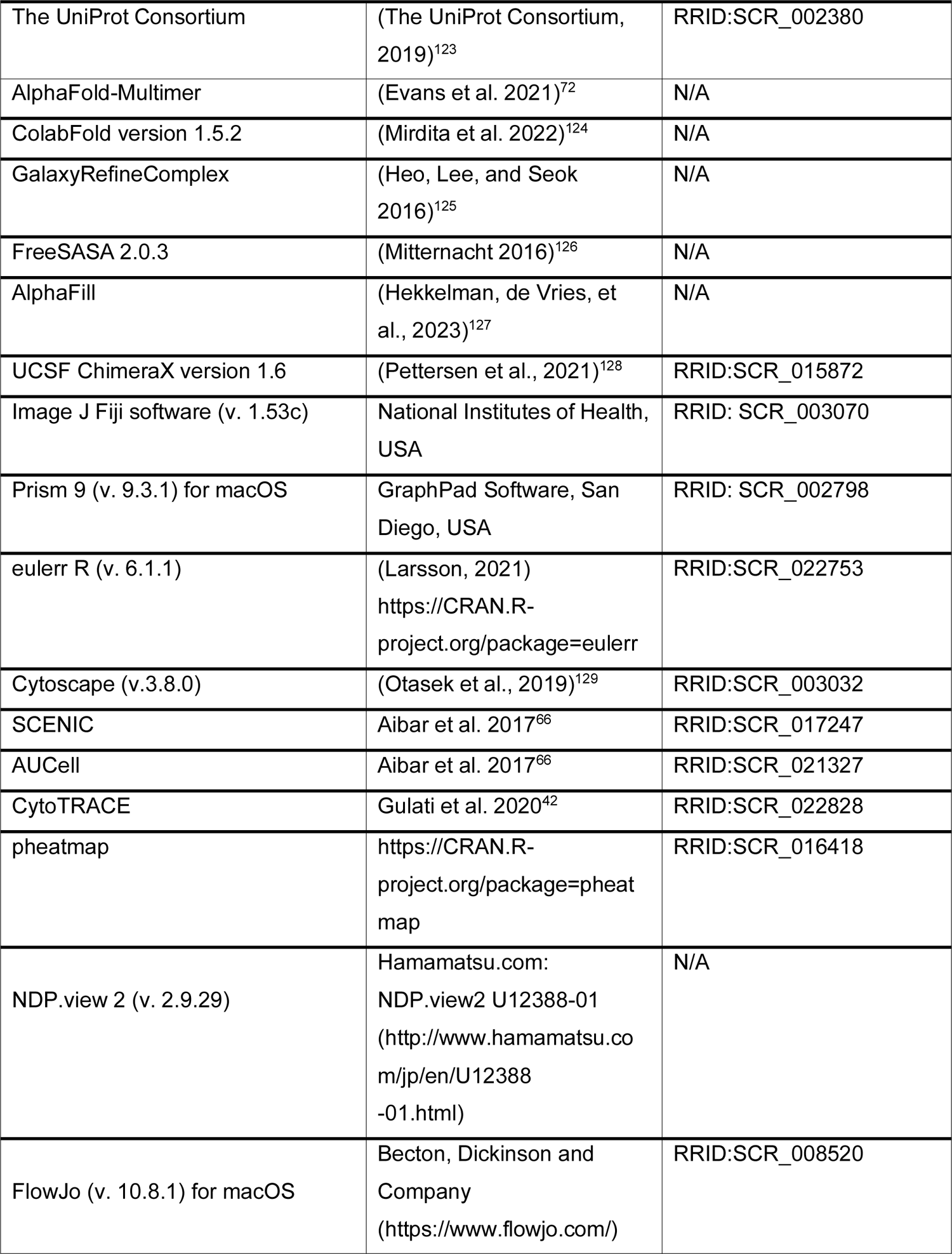

### RESOURCE AVAILABILITY

#### Lead contact

Further information and requests for resources and reagents should be directed to and will be fulfilled by the lead contact, E. Elizabeth Patton.

#### Materials availability

Plasmids generated in this study will be made available upon request made to the lead contact and will be deposited in Addgene. Patient derived low passage melanoma cells are available, upon MTA approval, upon request to.

### EXPERIMENTAL MODELS AND SUBJECT DETAILS

#### Zebrafish maintenance and husbandry

Zebrafish were maintained in accordance with UK Home Office regulations, UK Animals (Scientific Procedures) Act 1986, under project license P8F7F7E52. All experiments were approved by the Home Office and AWERB (University of Edinburgh Ethics Committee).

#### Zebrafish melanoma models

Zebrafish were genotyped using DNA extracted from tail fin clipped tissue by PCR to confirm the mutant allele status of *tp53^M^*^214^*^K^* (referred to as *p53^-/-^ or p53 mutant*) and *mitfa:BRAF^V600E^* as described in our previous publications^75^. The emergence of melanoma is usually observed in individuals aged 3- to 6-month-old. Individuals used in this study for DMSO control versus vemurafenib or Nifuroxazide drug pellets treatment were aged 5- to 6-month-old when entering the treatment scheme. Both female and male individuals were admitted into the treatment course regardless of the sex.

#### Human melanoma cell line culture

A375 cells were cultured in DMEM high glucose media, C089 cells were cultured in RPMI 1640 media, and patient sample derived cell line MPD001 were cultured in RPMI 1640 with 25 mM HEPES. All media were supplemented with 2 mM L-glutamine and 10% fetal calf serum, and all cells grown at 37 °C in a 5% CO_2_ humidified incubator. All cells have been routinely tested for mycoplasma, with the most recent test performed on June 21, 2023.

## METHOD DETAILS

### Establishment and amplification of patient derived melanoma cell lines

Melanoma Patients were managed in accordance with the ethical principles of Declaration of Helsinki and in accordance with Good Clinical Practice as defined by the International Conference on Harmonisation. All patients gave informed written consent to participate in clinical trials or EAP or EAMS. Patient samples were collected with written full-informed consent under Manchester Cancer Research Centre (MCRC) Biobank ethics application #07/H1003/161+5 and approval for the work under MCRC Biobank Access Committee application 13_RIMA_01. Tissue samples were collected from cutaneous melanoma patients at The Christie NHS Foundation Trust. For cell line MPD001 and MPD002, tumour fragments from the lymphatic melanoma and skin lesion to chest wall metastases were obtained respectively during the surgical procedure and dissociation was performed on the same day utilising the human tumour dissociation kit (Miltenyi Biotec) following manufacturer’s instructions. Briefly, after mechanical dissociation with a scalpel, tissue was resuspended in RPMI media with a mixture of Kit Enzymes (H, R and A) and placed on the gentleMACS^TM^ Dissociator, program 37°C _h_TDK_1 for 30 minutes. Next, cells were transferred to a 50 ml falcon tube and centrifuged at 300xg for 7 minutes, then resuspended and passed through a 70 µm cell strainer. This was centrifuged again (300xg for 7 minutes) and resuspended in 1X red blood cell lysis buffer (BioLegend) in deionized water, then incubated at room temperature, protected from light, for 15 minutes. After this step, the sample was centrifuged and finally resuspended and plated in a 10 cm cell culture petri dish in RMPI medium supplemented with FBS (Gibco) (10%) and Penicillin/Streptomycin (Gibco) (1%). Cells were cultured in an incubator at 37°C with 5% CO_2._ Cell lines were frozen and stored after a 2-week amplification period, freezing performed using FBS with DMSO (10%). Cryotubes were placed in an isopropanol freezing container (Nalgene® Cat. C1562-1EA) at −80°C for 24h for gentle freezing and transferred to liquid nitrogen tanks for long term storage.

### Generation of human melanoma mutant cell lines

Human melanoma cell lines A375 and C089 engineered with stable expression of *Streptococcus pyogenes* Cas9 were gifted to us and used to build *ALDH1A3* mutant cells^39^. Briefly, for *ALDH1A3* knockout, the vehicle plasmid expressing gRNA (pKLV2-U6gRNA5(BbsI)-PGKpuro2ABFP, Addgene: 67991) was engineered by Golden Gate cloning using restriction enzyme BbsI-HF (NEB) to express gRNA targeting ALDH1A3 (CGTCCCGGAGCAATCTGAAG). Lentiviral particles were produced by co-transforming the HEK293T (ATCC) cells with target plasmid, the packaging plasmid psPAX2 (Addgene: 12260), and pMD2.G (Addgene: 12259), facilitated with Lipofectamine 2000 (Invitrogen) following the manufacturer’s instructions. 48 hours post transfection, the 293T cell culture supernatant was collected and filtered (0.45 µm) to transfect the targeted recipient cell lines, supplemented with 10 µg/ml polybrene (Merck Millipore). 48-72 hours post transfection, the cells were split and seeded with the complete growth media containing 1 µg/ml puromycin (Life Technologies) to obtain clones with stable expression of the gRNA. Suitable single clones were validated by western blot to confirm full-length *ALDH1A3* knockout and Aldefluor assay to confirm the loss of ALDH activity before expanded for biological experiments. For *TFAP2B* knockout cells, the gRNA sequence was engineered similarly (GGACCGAGTGGTAGTCCCTC) using Golden Gate cloning. For vehicle control samples, the vehicle plasmid expressing empty gRNA were engineered to express mCherry instead of *BFP* sequence; for *ALDH1A3* and *TFAP2B* over-expression cells, the vehicle plasmid expressing empty gRNA were engineered to express *ALDH1A3* CDS instead of *BFP*. Briefly, restriction digestion enzyme KpnI-HF (NEB) and BsiWI-HF (NEB) were used to remove *BFP* sequence and create sticky ends matching the *mCherry*, *ALDH1A3* or *TFAP2B* CDS flanking sequence. The digested fragment of plasmid backbone and the target sequence were then ligated using T4 DNA ligase (NEB), following which a similar transfection and selection procedures were performed to acquire stable mutant lines.

### Human melanoma cell ALDH activity measurement

The ALDH enzyme activity in human melanoma cells was measured using the Aldefluor assay kit (StemCell Technologies) following the manufacturer’s instructions. In brief, melanoma cells dissociated with trypsin (Gibco) were resuspended in the Aldefluor buffer with the fluorescent bodipy-aminoacetaldehyde (BAAA) reagent included in the kits. For each experiment, a negative control vial was set up using a small aliquot out of the sample vial (100 μl out of 1 ml), supplemented with (5 μl) pan-ALDH inhibitor DEAB (diemethylaminobenzaldehyde) immediately after the resuspension. After incubation of 30 minutes at 37°C, the Aldefluor activity was measured using flow cytometry (Fortessa, BD Biosciences). For cell sorting to establish the ALDH^High^ and ALDH^Low^ cells, the stained cells were sorted by FACS Aria II (BD Biosciences) and the population with the highest and lowest 5% ALDH activity (ALDH^High^ and ALDH^Low^) were collected. All flow cytometry data were analysed using the software FlowJo. Dead cells were excluded using 7-Aminoactinomycin D (7-AAD, Eurogentec), or 4,6-Diamidine-2-phenylindole dihydrochloride (DAPI, Sigma-Aldrich).

### RNA extraction and RT-qPCR

Quantitative reverse transcription PCR (RT-qPCR) assays were performed by standard protocol suitable for LightCycler® 480 Instrument (Roche). In brief, total RNA was extracted and purified from live cells using TRIzol (Invitrogen) and Direct-Zol RNA Miniprep Kits (ZYMO Research). After quality check and measurement by NanoDrop™ (ThermoFisher), 1 µg RNA for each sample was reverse-transcribed using Superscript III reverse transcriptase (Life Tchnologies). Quantitative PCR were carried out by setting up reactions using the reverse transcribed cDNA template, primers (see also **Table S6, related to STAR Oligonucleotides**) and LightCycler ® 480 SYBR Green I Master reagent, run by the program of Standard Roche Template (System II). Reads of gene ACTB (Beta Actin) were used as the internal control to calculate the relative expression values.

### RNA-seq Pipeline

Libraries were prepared from 500 ng of each total-RNA sample using the TruSeq Stranded mRNA Library Kit (Illumina) according to the provided protocol and purified using AMPure XP beads (Beckman Coulter). Sequencing was performed using the NextSeq 500/550 High-Output v2 (150 cycle) Kit (# FC-404-2002) on the NextSeq 550 platform (Illumina Inc, #SY-415-1002). Libraries were combined in an equimolar pool based on the library quantification results and run across a single High-Output Flow Cell. Raw FASTQ sequence reads were quality checked using FastQC (v. 0.11.3) and aligned to the human genome (GrCh38) assembly using STAR (v. STAR_2.5.1b) software with default parameters. The quality of the resulting alignment to the transcriptome (Ensembl annotation version GRCh38.91) was checked using RNASeqQC (v. 1.1.8.1). Raw counts of reads covering the transcriptome (Ensembl annotation version GRCh38.91) were obtained using htseq-count (0.6.1) with the “-s reverse” option. Differential expression was analysed using the DESeq2 R package (v. 1.20.0).

### Pathway Enrichment Analysis

Gene set functional enrichment analysis (GSEA) was used to identify the enriched pathways at FDR < 0.05 (weighted Kolmogorov–Smirnov test) using GSEA software (Subramanian et al., 2005). The gene expression matrix of sorted ALDH^High^ versus ALDH^Low^ cells or selected melanoma patient ALDH1A3^High^ versus ALDH1A3^Low^ groups were used to compare against literature-based datasets **(Table S1, S2)** using gene set permutation settings. For gene over-representation analysis across literature curated signature terms, genes with differential acetyl-Histone H3 in WT compared to *ALDH1A3* knockout were selected based on FDR < 0.05 (in total 1599, see also **Table S4**) as input to the g:Profiler website^111^. The gene terms with FDR < 0.05 (hypergeometric test, BH adjusted) were considered significantly over-represented, with the term size cut-off set to 1000 (token: gp_33Kp_FaEr_Mcs). For gene over-representation analysis across the REACTOME database^130^, genes with acetyl-histone H3 peaks differentially enriched in WT or *ALDH1A3* knockout (see also **Table S4**) were separately tested and terms of FDR < 0.05 (hypergeometric test, BH adjusted) were considered significantly over-represented.

### Gene set expression correlation analysis

To assess the expression corelation between *ALDH1A3* and NCSC gene signature, gene set variation analysis (GSVA) ^120^ was used to calculate the relative expression score for the gene set containing only ALDH1A3 or the related signature gene lists. Spearman correlation analysis was then performed to evaluate the statistical correlation across all samples. Positive correlation was determined as Spearman co-efficient R>0.3 with exact critical probabilty p-value < 0.05.

### Human metastatic melanoma single cell data mining

Human metastatic melanoma single-cell RNA-seq expression matrix was accessed via supplementary data of Tirosh et al., 2016^51^. Cells classified as malignant tumour cells were extracted for UMAP clustering via Seurat (v.4.0.0), with *ALDH1A3* and *TFAP2B* visualised, clustering resolutions equals 0.5. For immune immune checkpoint blockade therapy naïve samples, single-cell RNA-seq expression matrix of human melanoma was accessed via the published study of Pozniak et al., 2024^106^ as deposited on European Genome-phenome Archive (EGA): EGAD00001009291, with the original patient tumour progression tracked and recorded for their anti-PD-1 and/or anti-CTLA-4 treatment outcomes. Cells classified as malignant tumour cells were extracted for violin plot analysis via Seurat (v.4.0.0), with *ALDH1A3* visualised and compared between the responder and non-responder groups based on the matched patient clinical records.

### *NRAS^Q61K/°^; Ink4a^−/−^* murine melanoma data mining

Single-cell RNA expression data of malignant melanoma cells, originating from 5 primary murine melanoma lesions^43^, were interrogated for *Aldh1a3*, ALDHhigh_enriched_signature and Tfap2b_regulon expression activities, measured by AUCell^66^. A Tfap2b regulon is inferred by using the pySCENIC pipeline on a mouse melanoma single-cell RNA-seq data^43^ with the genes extracted and visualized using Cytoscape (v.3.8.0)^131^. To map potential differentiation trajectories onto the single-cell UMAP space, we calculated for each melanoma cell its corresponding CytoTRACE score^42^, which is a measure of gene expression diversity and a surrogate for developmental potential (0<CytoTRACE score<1). CytoTRACE scores close to 1 are indicative of a less differentiated and close to 0 of a differentiated state. Besides the single cells, every gene was scored and correlated to either contribute to a dedifferentiated or differentiated state and ranked accordingly.

### Western Blot

Cells were detached by Trypsin (Gibco) and lysed on ice for 30 min at a cell density of 10^7^ cells per ml of RIPA lysis buffer (Merck Millipore), supplemented with phosphatase and complete protease inhibitors (Merck Roche). Debris of cells were removed by centrifuge (10,000 rpm, 10 min, 4 °C). Protein concentrations were determined using the BCA assay (Thermo Fisher) and 10-20 mg protein per lane was electrophoresed on 4-15% precast gradient gels (Bio-Rad). Based on the protein concentration and sample volumes, calculated amount of Laemmli (SDS) buffer (Enzo Life Science) were added and incubated with the samples at 95 °C for 5 min before gel loading. After the electrophorese program, gels were transferred onto Turbo™ transfer membranes (Bio-Rad) using the semi-dry Turbo Transfer system (Bio-Rad). Membranes were then blocked with 5% w/v BSA/TBS for 30 min at room temperature, subsequently probed by primary antibodies with optimised dilution factors overnight at 4°C, incubation of goat anti-mouse IRDye 680- or goat anti-rabbit 800-labelled secondary antibodies (LI-COR Biosciences) and imaged using an Odyssey infrared scanner (LI-COR Biosciences).

To probe the histone proteins using western blot, an acid extraction protocol was carried out instead of RIPA lysing protocol. In brief, the cells were washed with ice-cold PBS and lysed on ice for 10 min at a cell density of 10^7^ cells per ml of Triton Extraction Buffer (TEB: PBS, 0.5% Triton X 100 (v/v), 2 mM phenylmethylsulfonyl fluoride (PMSF, Life Technologies)), supplemented with 5 mM sodium butyrate (Abcam) to retain levels of histone acetylation. The nuclei of cells were centrifuged (650 x g, 10 min, 4 °C) and washed in half the volume of TEB and centrifuged again. Pellets were re-suspended using 0.2 N HCl at a density of 4×10^7^ nuclei per ml. The histones were extracted overnight at 4°C and the debris were removed by centrifuge (650 x g, 10 min, 4 °C). 5M NaOH were added at 1/20 of the volume of the supernatant to neutralise the samples. The same steps of protein measurement and western blot were followed as described above for probing protein in the total cell lysates.

The primary antibodies used are as following: ALDH1A3 (1:10,000, Rabbit, Invitrogen), TFAP2B (1:1000, Mouse, Santa Cruz), beta-Actin (1:1000, Mouse, Invitrogen), Histone H3 (1:1000, Mouse, Abcam), acetyl-Histone H3 (1:10,000, Rabbit, Millipore).

### Co-immunoprecipitation of ACSS2

Co-immunoprecipitation was performed using SureBeads™ Protein A Magnetic Beads (BioRad) following recommended instructions from the manufacturer. Briefly, to prepare the beads for binding target protein, for each 1 million cells, 5 µl SureBeads were washed in 0.1% TBST for 3 times and then incubated with 1:50 ACSS2 primary antibody (Rabbit, Cell Signaling) or 1:50 IgG control (Cell Signaling Technology) at room temperature for 30 minutes. Unbound antibodies were removed by another 3 times of washing using 0.1% TBST. To prepare the protein lysates, cells were cultured to 80-90% confluence before dissociated using trypsin/EDTA. Cells were pooled into a master tube before aliquoting 20 million cells per sample for whole cell co-IP or 40 million cells per sample for nuclear department co-IP.

For whole-cell co-IP, aliquoted cells were pelleted by centrifuge at 300 xg, for 3 min and resuspended in 1x RIPA buffer supplemented with Roche protease inhibitor cocktail and lysed on ice for 10 min. The lysates were then centrifuged at 12,000 xg for 10 min before the supernatant were transferred to primary antibody conjugated SureBeads Protein A Magnetic Beads for incubation overnight at 4°C. For nuclear protein co-IP, nuclei isolation were performed by adding 5 ml nuclei isolation buffer (NIB) to the cell pellets (15 mM Tris, 60 mM KCl, 15 mM NaCl, 5 mM MgCl_2_, 1 mM CaCl_2_, 250 mM sucrose, pH adjusted to 7.5, supplemented with 1 mM DTT and 0.1% NP-40) with gentle pipetting and incubation on ice for 5 minutes. Nuclei were spun down by centrifuging at 600 xg for 5 min at 4 °C and then proceed to the same RIPA lysing steps as described for the whole cell co-IP assay. For both whole-cell and nuclei co-IP, 10% of lysates were set aside as input before proceeding to adding SureBeads conjugated antibodies.

Following incubation at 4°C overnight, unbound proteins were washed off by rinsing the magnetic beads 3 times using 0.1% TBST. Captured proteins were released by adding 50 µl 1x RIPA buffer and 10 µl 6x Laemmli SDS reducing buffer to the beads slurry and being heated at 95 °C for 3 minutes. Proteins were then probed by standard Western Blot as described above.

### Colony formation assay in Soft Agar

For each sample in a 6-well plate set up, 5000 cells were suspended in serum-free growth media supplemented with 10 ng/ml EGF, 10 ng/ml FGF, B27 supplement (1x, Thermo Fisher Scientific), and 0.3% low melting agarose (Sigma-Aldrich). Cells were then layered over a solid base of 0.5% low melting agarose, cultured for 18 days (A375) or 25 days (C089), optimised by the colony growth speed. Colonies (>10 cells) from 10 separate fields of each sample were then manually counted using the images captured by a Nikon DS-L3 camera system on an Eclipse TS100 microscope (Nikon) with a 4X objective.

### Immunocytochemistry with fluorescence labelling and imaging

Cells were cultured on the collagen I coated coverslip glasses (VWR) for ICC assays. Samples were washed twice with PBS and fixed with 4% paraformaldehyde (PFA, Thermo Fisher Scientific) at room temperature for 15 min before permeabilized by PBST (0.1% triton x-100/PBS) for 10 min. Coverslips were then incubated in 3% BSA/PBST (w/v) for 30 min before probed with primary antibodies at appropriate dilution for 4 hours at room temperature or 4°C overnight (ALDH1A3, 1:300, Abcam; TFAP2B, 1:100; ACSS2, 1:100, Abcam). After being washed 3 times in PBST for 5 min each, samples were incubated with fluorochrome-conjugated secondary antibodies (Donkey anti-Rabbit 488, Donkey anti-Mouse 568, and Donkey anti-Rabbit 647, all 1:1000, Invitrogen) for 30 min in the dark. After 3 times of wash in PBST for 5 min each, DAPI (Sigma-

Aldrich) was added to the final wash of PBST to stain the nucleus, and the coverslips were mounted with antifade mounting medium (Vectashield, 2BScientific) before fluorescent microscope imaging using multimodal Imaging Platform Dragonfly (Andor technologies, Belfast UK). Images were acquired using a 20X or 40X lens equipped with 405, 488, 561, and 640 nm lasers built on a Nikon Eclipse Ti-E inverted microscope body with Perfect focus system (Nikon Instruments, Japan). Data were collected in Spinning Disk 25 μm pinhole mode on the Zyla 4.2 sCMOS camera using a Bin of 1 and no frame averaging using Andor Fusion acquisition software.

For ICC on flow cytometry sorted ALDH^High^ and ALDH^Low^ cells, Aldefluor-sorted cells were resuspended in normal culture media and allow to attach on the collagen I coated coverslip glasses (VWR) in 6-well plates for 4-6 hours before fixed to ensure the ALDH activity states.

For interrogating ALDH1A3-TFAP2B-ACSS2 co-localisation, primary antibodies raised in the same species, *i.e.* anti-ALDH1A3 and anti-ACSS2 Rabbit IgG, were pre-conjugated to different fluorescence dye (Alexa Fluor 488 and Alexa Fluor 647) separately following the manufacturer’s instructions (Zenon Rabbit IgG labelling kits, Thermo Fisher) before incubation with the samples together with Mouse anti-TFAP2B primary antibody for 2 hours at room temperature. The cells were then washed three times in PBS and incubated with Donkey-anti-mouse conjugated Alexa Fluor 568 antibodies for 0.5 hour at room temperature (Invitrogen). Cells were washed, nuclear stained with DAPI (Sigma) and mounted in Vectashield® mounting media before imaging. Super-resolution images were acquired using instant Structured Illumination Microscopy (SIM)^132^ with Nikon SoRa™ system. Imaging was carried out using an SR HP Plan Apo λS 100x 1.35NA Silicone lens (Nikon Instruments). The CMOS cameras used for acquisition were Teledyne Photometrics Prime 95B (Teledyne Photometrics 3440 E.Britannia Drive, Tucson AZ) and 405 / 488 / 514 / 561 / 640nm laser lines. Z-step size for Z stacks was set to 0.120 μm as required by manufacturers software. Acquisition of images and deconvolution was carried out using the (3D algorithm) Nikon NIS Elements Advanced Research software. Settings for acquisition and reconstruction were identical in all images. Line scanning signal correlation analysis were carried out in Fiji by taking the intensity from each channel across the scanning line as an individual variable and conducting Pearson correlation test.

### ^13^C_6_-glucose tracing via targeted UPLC-MRM/MS

For ^13^C_6_-glucose tracing experiment, DMEM with no glucose, no glutamine, no phenol red (Gibco) was purchased and then supplemented with 4.5 g/L ^13^C6-glucose (Merck Life Sciences), 2 mM L-glutamine, and 10% FCS (referred to from now on as the ^13^C_6_-glucose DMEM). To trace ^13^C_6_-glucose in A375 ALDH^High^ and ALDH^Low^ cells, A375 melanoma cells were cultured in T175 flasks (Corning) until reaching 70% confluence, then incubated in ^13^C_6_-glucose DMEM for 12 hours before dissociated for Aldefluor staining and live sorted by FACS Aria II (BD Biosciences). Every sample contains ∼0.5 million sorted cells, which was immediately snap frozen in liquid nitrogen upon sorting. To trace ^13^C_6_-glucose in A375 Cas9 control and *ALDH1A3* knockout cells, cells were seeded in 6-well plates until reaching 50% confluence, then incubated in ^13^C_6_-glucose DMEM for 24 hours before trypsin dissociation and liquid nitrogen snap frozen. Metabolites extraction, derivation, and targeted UPLC-MRM/MS profiling the central carbon metabolites were performed as described in Han et al., 2013^133^.

### Histone acetylation profiling using bottom-up mass spectrometry

Histone extraction and derivatization workflow was optimised based on Sidoli et al., protocol^134^. Briefly, nuclei from live attached melanoma cells were isolated by 5-minute incubation on ice with NIB buffer (15 mM Tris, 60 mM KCl, 15 mM NaCl, 5 mM MgCl_2_, 1 mM CaCl_2_, 250 mM sucrose, pH adjusted to 7.5, supplemented with 1 mM DTT, 0.5 mM PMSF, 0.05 µM NaF, 0.05 µM NaVO_4_ and 10 mM sodium butyrate, 0.1% NP-40) followed by scraping. Nuclei were spin down by centrifuging at 1000 rcf for 5 min at 4 °C. The nuclei pellets were washed twice by NIB without NP-40. Histone proteins were extracted by adding 0.2 M H_2_SO_4_ in a 1:4 ratio of nuclear pellet to H_2_SO_4_ and incubated for 4 hours at 4 °C. Histones were then precipitated by adding 1:3 v/v Trichloroacetic acid (TCA) and incubated overnight at 4 °C. For chemical derivatization, precipitated histone proteins were air-dried by vacuum centrifuge and rinsed with ice-cold acetone prior to four rounds of propionylation, with the last two rounds of propionylation carried out on histone peptides post trypsin digestion. Propionlated peptides were transferred to C18 staging tips for desalting and eluted using 80% acetonitrile (ACN) with 0.1% trifluoracetic acid (TFA) before LC-MS analysis.

Peptides resulting from all digestions were separated by nanoscale C18 reverse-phase liquid chromatography using an UltiMate 3000 RSLCnano system coupled online to an Orbitrap Fusion Lumos Tribrid mass spectrometer (Lumos) (all Thermo Fisher Scientific). HPLC buffers-0.1% formic acid in HPLC-grade water (buffer A); 0.1% formic acid in HPLC-grade acetonitrile (buffer B) were prepared. HPLC method was programmed as follows: from 0 to 30% buffer B in 30 min, from 30 to 100% B for the next 5 min and at isocratic 100% B for 8 min. the flow rate was set to 250-300 nl/min.

Acquisitions were carried out in data independent acquisition mode (DIA) using Tune application 3.5.3890 (Thermo Scientific). A nanoelectrospray ion source (Sonation) was used for ionisation in positive mode. Chromatography was carried out at a flow rate of 250-300 nl/min using 50 cm fused silica emitters (CoAnn Technologies) packed in house with reverse phase Reprosil Pur Basic 1.9 µm (Dr. Maisch GmbH). The emitter was heated to 50°C using a column oven (Sonation), and an Active Background Ion Reduction Device (ABIRD) was used to decrease air contaminants signal level. Peptides were eluted with a 60-minute two-step gradient, over a total run time of 90 minutes. A full scan was acquired at a resolution of 60,000 at 200 m/z, over mass range of 300-1100 m/z, followed by a DIA scan. All precursors were fragmented using 15 consecutive windows with 50 Da width, allowing for a 1 m/z overlap, covering a mass range from 349.5 to1100.5 m/z. Higher energy collisional dissociation fragmentation spectra were recorded at 15,000 resolution at 200 m/z. All ions were fragmented using normalised collision energy of 28%, for a maximum injection time of 54 ms, or a normalised AGC target of 1000%.

Data analysis was performed on MATLAB using the EpiProfile2.1_1Basic package (DOI: 10.1021/acs.jproteome.8b00133) using label-free settings (nsource=1) or for histone H3 acetylation and 13C2-acetyl incorporation analysis the C13 on acetylation group (nsource=3). Isotopic correction was performed by MATLAB as implemented in EpiProfile.

### Acetyl-CoA extraction and LC-MS analysis

To extract acetyl-CoA from whole cell lysates and nuclear department, samples were collected as described by Trefely et al., protocol^135^. Briefly, live attached melanoma cells were isolated by 5-minute incubation on ice with NIB buffer (15 mM Tris, 60 mM KCl, 15 mM NaCl, 5 mM MgCl_2_, 1 mM CaCl_2_, 250 mM sucrose, pH adjusted to 7.5, supplemented with 1 mM DTT and 0.1% NP-40) followed by scraping. 10% of each sample were removed and quenched in 1 ml ice-cold 10% TCA as total lysates. Nuclei were spun down by centrifuging at 600 rcf for 5 min at 4 °C. The nuclei pellets were washed twice by NIB without NP-40 and then quenched in 1 ml 10% TCA. Acetyl-CoA was extracted using solid phase extraction to remove TCA. Extraction cartridges (Oasis^®^ HLB 1cc (30 mg)) were conditioned by running 1000 µL of methanol (MeOH) following by 1000 µL of water (H_2_O). On the cartridges, 100 µL of samples were then loaded, washed with 1000 µL of H_2_O and eluted with 500 µL of MeOH. The extracts were then dried under nitrogen and reconstituted in 80% ACN: 20% H_2_O (20 mM ammonium carbonate 0.1% ammonium hydroxide solution 25%).

Samples were analysed on a Dionex UltiMate 3000 LC System (Thermo Scientific, Waltham, Massachusetts, EUA) coupled to a Q Exactive Orbitrap Mass Spectrometer (Thermo Scientific, Waltham, Massachusetts, EUA) operating in negative polarity with scan range from 806 to 815 m/z. Chromatographic separation was achieved using a ZIC®-pHILIC 150 x 2.1 mm column (Merck Millipore Sigma, Burlington, Massachusetts, EUA) at 45°C using a gradient starting from 20% buffer A (20 mM ammonium carbonate 0.1% ammonium hydroxide solution 25%), and 80% B (acetonitrile) to 80% buffer A, 20% buffer B at 9.5 minutes and reconditioning the column to the initial condition until 14.5 minutes. Mass spectrometry data were processed using Skyline^136^ on a targeted fashion by matching accurate mass and retention time with standard.

### A375 acetyl-Histone H3 ChIP-Seq

The control and *ALDH1A3* KO melanoma cells were cultured to 80% confluency and harvested by dissociation with trypsin in PBS/EDTA. Cells were resuspended in PBS and immediately fixed in 1% formaldehyde in PBS for 10 minutes at room temperature. The fixation was terminated by adding glycine and incubated for an additional 5 minutes. After being washed in cold PBS, cell pellets were resuspended in 150 μl chilled lysis buffer (1% SDS, 10mM EDTA, 50 mM Tris-HCl pH8.1, 1x protease inhibitor cocktail, 1x PhosSTOP phosphatase inhibitors (Roche), 5 mM sodium butyrate (Sigma) and fresh 1mM DTT) and supplemented with 850 μl 1% Triton X IP dilution buffer (1% Triton X, 20mM Tris-HCl pH8.1, 150mM NaCl, 2mM EDTA, 1x protease inhibitor cocktail (Roche), 1x PhosSTOP phosphatase inhibitors (Roche), 1mM DTT, 5 mM sodium butyrate and 1 mM PMSF) and incubated on ice for 10 min. Lysed cells were sonicated on ice for 8x 30 sec on / 30 sec off burst cycles with a probe sonicator (SoniPrep150) in a chilled ice-water bath (12 Amplitude) to yield chromatin fragments ranging between 200 – 800 bp in length. Sheared chromatin was centrifuged at 16,000xg for 10 minutes at 4°C and the soluble supernatant transferred to new tubes. Each 500 μl chromatin samples were supplemented with 5 μl (5mg/ml) BSA. 10% of the input was stored and the rest was used for the immunoprecipitation.

Rabbit anti-acetyl-Histone H3 Antibody (Millipore) were pre-bound to magnetic SureBeads (BioRad) in 10% w/v BSA in PBS according to the manufacturer’s instructions for approximately 1h at 4°C with rotation, following which free antibody was removed with 3 washes of cold 10% w/v BSA in PBS. Chromatin and proteinG beads were combined and incubated over night at 4°C (at a ratio of 500 ng of bead bound antibody per 1 million cell equivalents of chromatin). Samples were washes at 4°C with rotation through the following series: 2 times in 1% Triton X IP dilution buffer, 2 times with ChIP wash A (50mM HEPES ph7.9, 500mM NaCl, 1mM EDTA, 1% Triton X-100, 0.1% Na-deoxycholate, 0.1% SDS. 1x protease inhibitor cocktail, 1x PhosSTOP phosphatase inhibitors (Roche) and fresh 1mM DTT) and 2 times with ChIP wash B (20mM Tris pH 8.0, 1mM EDTA, 250mM LiCl, 1% NP-40, 0.1% Na-deoxycholate, 1x protease inhibitor cocktail, 1x PhosSTOP phosphatase inhibitors (Roche) and fresh 1mM DTT). Finally, the samples were washed with TE (1mM EDTA, 10mM Tris pH8.0). The samples were resuspended in TE and supplemented with preheated 37°C Extraction buffer (0.1M NaHCO3 and 1% SDS), vortexed and incubated for 15 minutes at 37°C on a vibrating platform.

The pH of the extracted chromatin was adjusted by adding 6μl 2M Tris-HCl pH6.8 following which both the ChIP and input samples were incubated with 20μg RNAse A (NEB) at 65°C for 1 hour. Cross-links were reversed and the protein degraded by the addition of 20μg Proteinase K and incubation at 65°C for 6-8 h. Following removal of the magnetic SureBeads from the ChIP samples, DNA was purified using a Qiagen PCR cleanup kit following manufacturer’s instructions. DNA libraries were prepared using NEBNext® Ultra™ II DNA Library Prep Kit for Illumina® and NEBNext® Multiplex Oligos for Illumina® (Index Primers Set 1) following manufacturer’s instructions. Sequencing was performed using the NextSeq 500/550 High-Output v2.5 (150 cycle) Kit (#20024907) on the NextSeq 550 platform (Illumina Inc, #SY-415-1002). PhiX Control v3 (Illumina, #FC-110-3001) was spiked into the library pool at a concentration of ∼1% to enable troubleshooting in the event of any issues with the run.

### ChIP-seq data analysis pipeline

Basecall data produced by the NextSeq 550 is automatically uploaded to BaseSpace, a cloud-based data management and analysis service provided by Illumina. Here it is converted into FASTQ files and mapped to was mapped to the human genome (GRCh19) using bowtie2 (bwa-0.7.9) applications directly accessible through BaseSpace. BAM files were uploaded to the open access bioinformatic community platform Galaxy (the public server at usegalaxy.org) for downstream analysis^122^. All datasets and analysis history can be accessed (via https://usegalaxy.org/u/yuting_lu/h/a375-acetyl-histone-h3-chipseq-wt-vs-aldh1a3-knockout).

Briefly, mapped regions (due to fragment processing) extended beyond the end of the chromosomes were removed using SAMtools. MACS2 calling broadpeak algorithm was used to identify histone H3 acetylation regions, with each ChIP BAM file paired with the input control, cut off --mfold 5 50, band width --bw 500, FDR --qvalue 0.05. Call peak results (gapped peaks) were used for differential binding (histone acetylation in this case) analysis between A375 WT and ALDH1A3 knockout cells using the DiffBind^119^. R package ChIPseeker^137^ was used for adjacent gene annotation and genomic distribution analysis. To visualise the histone H3 acetylation sites on the genome browser UCSC^138^ mapped reads were converted to bigWig files (.bw) using the bamCoverage (deepTools2) algorithm^118^, with replicants for each condition merged.

### ACSS2 ChIP qPCR

ACSS2 bound chromatin were pulled down using rabbit anti-ACSS2 antibody (Cell Signaling Technology) with the same ChIP protocol described above in A375 acetyl-Histone H3 ChIP-seq, ChIP-ACSS2 DNA and input control were purified using the QIAquick PCR Purification Kit (Qiagen) and used for qPCR with the same LightCycler 480 SYBR Green (Roche) system as described above in the qPCR method for RT-qPCR. The ChIP qPCR primer sequences are listed in **Table S6**.

### MPD002 ALDH1A3 knockdown

ALDH1A3 knockdown was achieved using siRNA Transfection. In brief, MPD002 cells were seeded in 6-well plate at a density of 5 × 10^4^ cells/well and incubated overnight for attachment. Cells were then incubated with 10 nM ALDH1A3 siRNA or scrambled control (Life Technology) in 0.7 ml Opti-MEM™ media with Lipofectamine RNAiMAX (4.5μl/well)(Invitrogen) for 6 hours. Medium was then changed to normal culture condition for another 42-64 hours (48-72h in total post transfection) and the cells were collected for downstream analysis.

### MPD002 acetyl-Histone H3K23 and H3K27 CUT&TAG

The CUT&TAG experiments were performed following the protocol established by the Henikoff lab^139^ in principle. In brief, the control and *ALDH1A3* knockdown melanoma cells as well as spike-in control mouse embryonic stem cells (mESCs) were cultured to 90% confluency in 6-well plate and harvested by dissociation with trypsin in PBS/EDTA. Cells were resuspended in PBS supplemented with 5 mM sodium butyrate (Sigma) and counted. For each sample, 50,000 target cells (MPD002) and 5,000 spike-in control cells (mESCs) were combined and aliquoted for nuclei extraction (50 µl per sample, 20 mM HEPES pH7.5; 10 mM KCl; 0.5 mM Spermidine; 0.1% Triton; 1× Protease inhibitor cocktail; 20% v/v glycerol; 10 min on ice). Nuclei pellets were then collected by centrifuge for 4 min at 1,300 × g at 4°C and washed once in PBS supplemented with 5 mM sodium butyrate (Sigma) before binding to prepared concanavalin A coated magnetic beads (Bangs Laboratories, add 3.5 ul per sample to 50 µl binding buffer: 20 mM HEPES pH 8.0; 10 mM KCl; 1 mM CaCl2; 1 mM MnCl2; incubate 20 min at RT). The bead-bound nuclei were resuspended in 25 µl Wash Buffer (20 mM HEPES pH 7.5; 150 mM NaCl; 0.5 mM Spermidine; 1×

Protease inhibitor cocktail) and the appropriate primary antibodies were added at 1:50 dilution ratio (Rabbit anti-H3K23ac, Invitorgen; Rabbit anti-H3K27ac, Cell Signaling Technology; CUT&TAG IgG control, Cell Signaling Technology). Primary antibody incubation was performed on a rotating platform overnight at 4 °C. Unbound primary antibodies were removed by placing the sample tubes to magnet stand and discarding all supernatant liquid. Next, 25 µl of 1:100 diluted goat anti-rabbit unconjugated secondary antibodies (Abcam) were added to each sample to increase the number of Protein A binding sites, with samples incubated on a rotating platform at RT for 1 hour. Samples were then rinsed in Wash Buffer for 2-3 times before 1:200 dilution of pA-Tn5 adapter complex (EpiCypher) were prepared in high salt wash buffer (20 mM HEPES, pH 7.5, 300 mM NaCl, 0.5 mM Spermidine, 1× Protease inhibitor cocktail) and added to each sample (1.25 ul for each 25 ul reaction volume). pA-Tn5 incubation was performed at RT for 1 h on a rotating platform, before the unbound pA-Tn5 enzymes were removed by washing the beads using Wash Buffer on a magnet stand. Finally, tagmentation reaction were performed by resuspending the pA-Tn5 bound samples in 50 µl tagmentation buffer (High Salt Wash Buffer plus 10 mM MgCl2) and incubate at 37 °C for 1 h. Tagmentation was terminated by removing the tagmentation buffer using magnet stand and wash once in TAPS wash buffer (10 mM TAPS, 0.2 mM EDTA). To release the DNA, 5 µl SDS release buffer was added to each sample (0.1% SDS in 10 mM TAPS, 0.2 mM EDTA) with samples incubated at 58°C for 1 h in a PCR cycler.

To prepare the libraries for sequencing, 15 μl of 0.67% triton water solution were added directly to the bead slurry, plus 2.5 μl each of 10 μM uniquely barcoded i5 primer and i7 primers, using a different barcode combination for each sample. Next, 25 µl NEBNext HiFi 2× PCR Master mix (NEB) was immediately added and mixed before proceeding to PCR cycles (58 °C for 5 min (gap filling); 72 °C for 5 min (gap filling); 98 °C for 30 s; 14 cycles of 98 °C for 10 s and 60 °C for 10 s; final extension at 72 °C for 1 min and hold at 10 °C). Post-PCR clean-up was performed by adding 1.3× volume (65 µl) of Ampure XP beads (Beckman Counter) to the PCR reaction (50 µl) and incubating for 15 min at RT. The beads were then washed twice in 80% ethanol, with the final DNA libraries eluted in 22 µl 10 mM Tris pH 8.0. Library quality and fragment sizes were examined by high sensitivity TapeStation (Agilent) before sequencing.

### CUT&TAG data analysis pipeline

Basecall data produced by the NextSeq 550 is automatically uploaded to BaseSpace. The FASTQ files were mapped to the human genome (GRCh38) and the spike-in control genome (mm9) using bowtie2 (bwa-0.7.9). The mapped read counts splitting between target species (human) and spike-in control (mm9) were then used for calculating the scaling factors. Next, mapped regions were filtered using SAMtools for the flag of PCR duplication. Filtered BAM files were used for MACS2 calling broadpeak algorithm to identify histone H3K23 and H3K27 acetylation regions respectively, with each BAM file paired with the IgG control from the matching experiment group, (cut off -- mfold 2 50, band width --bw 300, FDR --qvalue 0.05). Peak calling results (gapped peaks) were used for differential peak analysis by applying edgeR differential analysis (fold change >1, FDR-q <0.05) between the control and ALDH1A3 knockdown master list peak reads (n=2), which were compiled by retrieving the filtered BAM reads count from the gapped peak bed coordinates, normalised by sequencing depth, and calibrated using the corresponding scaling factor. R package ChIPseeker^137^ was used for adjacent gene annotation and genomic distribution analysis. Representative peak on gene tracks were visualised using UCSC genome browser following bigwigAverage combining the replicate sample bigwig output from filtered BAM alignment files, which were individually normalised by genome coverage and calibrated using scale factors (bamCoverage --normalizeUsing RPGC --scaleFactor).

### AlphaFold Multimer modelling of ALD6-ACS2 and ALDH1A3-ACSS2 interaction

Canonical sequences of yeast ALD6 (UniProt accession: P54115) and ACS2 (P52910) and human ALDH1A3 (P47895) and ACSS2 (Q9NR19) were retrieved from UniProt^123^. Disordered N-termini of the human proteins, the first 24 residues of each, were removed to facilitate complex prediction. AlphaFold-Multimer^72^ predictions were performed with LocalColabFold (https://github.com/YoshitakaMo/localcolabfold), running ColabFold version 1.5.2^124^ on a single 350GB NVIDIA A100 GPU. Both yeast and human sequences were run as dimers of ALD6/ALDH1A3 and monomers of ACS2/ACSS2 because of sequence length limitations. We used templates available in the Protein Data Bank^140^ and the “mmseqs2_uniref” option for the – msa-mode flag. For both yeast and human complexes, we generated 3 models with 5 recycles and excluded models that were incompatible with the tetrameric structure of ALDH1A3. Confidence is 45% for yeast and 47% for the human complexes, calculated as 0.2 × pTM + 0.8 × ipTM^72^. The highest ranking models of each complex were further refined with GalaxyRefineComplex^125^, using default settings of protocol 2. Solvent accessible surface was calculated at residue level with FreeSASA 2.0.3^126^ with the buried surface area defined as the difference in solvent accessible surface area between the monomer and the complex. Homologous residues were determined via sequence alignment with MUSCLE^141^. Conservation of ACS2/ACSS2 interface with ALD6/ALDH1A3 was determined by calculating the Pearson correlation between the buried surface area values of homologous residues. ATP/CoA binding pocket of ACSS2 was visualised by structural alignment of AlphaFill-optimized^127^ protein-ligand complexes (ATP donor: PDB ID 5k8f; CoA donor: PDB ID 3gpc) to the AlphaFold-Multimer predicted model. Visualisation of protein structures was performed with UCSF ChimeraX version 1.6^128^. The PDB models of ACSS2-ALDH1A3 and Acs2-Ald6 will be released on ModelArchive upon the publication of this manuscript and is currently available for reviewers via Mendeley Data (Reserved DOI:10.17632/4hfw7rpspb.1).

### Zebrafish melanoma ALDH activity measurement

Zebrafish melanoma live cells were dissociated from freshly dissected tumour samples as described in Travnickova et al., 2019^10^. Cell suspensions were sized to 1 million cell count per ml and incubated with AldeRed (ThermoFisher Technologies) following the instruction by the manufacturer. After incubation of 1 hour at 28°C, the AldeRed activity was measured using flow cytometry (Fortessa, BD Biosciences). For cell sorting to establish the zebrafish ALDH^High^ and ALDH^Low^ cells, the stained cells were sorted by FACS Aria II (BD Biosciences) and the population with the highest and lowest 5% ALDH activity (ALDH^High^ and ALDH^Low^) were collected, as is described for AldeFluor guided ALDH subpopulation selection.

### Zebrafish drug pellet treatment

To perform drug treatment on adult zebrafish bearing melanoma, we produced fish bite-size drug pellets as described in our previous publication^77^ and fed single-housed individual fish with fish food agar pellets containing DMSO (Sigma Aldrich), vemurafenib (SelleckChem), and/or Nifuroxazide (Merck Millipore) once per day. Zebrafish under the drug treatment procedure were fed daily in the AM and early PM with artemia, and then fed the drug pellets in the late PM (6-8 PM). Zebrafish actively sought for and consumed the drug pellets voluntarily without any handling. Zebrafish under continuous drug treatment were imaged one day pre-treatment and once every week to track tumour size change.

### Imaging of adult zebrafish tumour and size measurement

Zebrafish were briefly anesthetised (Tricaine in PBS 1:10,000 concentration) for no longer than 10 min per session and fully recovered in fresh system water. Brightfield images were taken for each fish positioned on both sides. Images of fish lesions were captured at the same magnification scale every week using a Nikon COOLPIX5400 camera attached to a brightfield microscope (Nikon SMZ1500). The size of each lesion was quantified by using the manual field selection in Fiji on each tumour image, then compared to the matching pre-treatment lesion to calculate the relative percentage change. Lesions that could be observed from both sides of the fish were measured by combining the area number averaged from both sides.

### Zebrafish histology and IHC quantification

Zebrafish melanoma samples were collected, fixed, and processed as described in our earlier publications^10,17,142^. The slides of Haematoxylin and Eosin staining were imaged using a Hamamatsu NanoZoomer SlideScanner, and the images were processed using NDP.3 software. Aldh1a3 expression was assessed using Rabbit polyclonal anti-ALDH1A3 primary antibody (1:200, Abcam); Cd79a expression was probed using Rabbit monoclonal anti-CD79a (1:200, Abcam). Following secondary fluorescent antibody incubation (Donkey anti-mouse conjugated Alexa Fluor 568, Donkey anti-Rabbit conjugated Alexa Fluor 488 or 647, Invitrogen), nuclei were stained with DAPI dye (1:1000, Life Technologies). Stained tissue slides were mounted with antifade mounting medium (Vectashield, 2BScientific) before fluorescent microscope imaging using multimodal Imaging Platform Dragonfly (Andor technologies, Belfast UK). Similar to ICC imaging, images were acquired using a 20X lens equipped with 405, 488, 561, and 640 nm lasers built on a Nikon Eclipse Ti-E inverted microscope body with Perfect focus system (Nikon Instruments, Japan). Data were collected in Spinning Disk 25 μm pinhole mode on the Zyla 4.2 sCMOS camera using a Bin of 1 and no frame averaging using Andor Fusion acquisition software. Standard deviation intensity (STD) projection of a confocal z-stack was performed in Fiji to allow intensity quantification and cell subpopulation assessment.

### Quantification and Statistical Analysis

All statistical methods used in the paper are described in the figure legends and, where indicated, additional details are provided in the method details. Definitions of sample size, measures of centre and dispersion, and precision measures are also indicated in figure legends. Statistics were computed using R and GraphPad Prism. When appropriate, corrections for multiple comparisons were implemented as indicated in the figure legends.

### CRedit Author contributions

Conceptualization: YL, AC, EEP

Data curation: YL, ZK, JAM, AvK

Formal Analysis: YL, MB, FR, PL

Funding acquisition: FR, AvK, JAM, OS, RI, EEP

Investigation: YL, MB, FR, AC, ZK

Methodology: YL, JT, MB, ZK, JAM, LM, AvK

Project administration: YL, EEP

Resources: YL, JT, PGM, AHC, CJS, RM, VSP, OS, EEP

Supervision: AvK, CJS, JAM, VSP, RI, EEP

Validation: YL, RI

Visualization: YL, MB, FR, EEP

Writing – original draft: YL, MB, RI, EEP

Writing – review & editing: YL, JT, AvK, CJS, RI, EEP

## Inclusion and diversity

One or more of the authors of this paper self-identifies as an underrepresented ethnic minority in their field of research or within their geographical location. One or more of the authors of this paper self-identifies as a gender minority in their field of research.

## Acknowledgements

We thank James Chen (Stanford University), Kevin Myant (CRUK Scotland, Edinburgh) and Wendy Bickmore (MRC Human Genetics Unit) for helpful discussions. We thank Sergio Lilla (Cancer Research UK Beatson Institute, CRUK Scotland Centre) for MS proteomic analysis (funded by Cancer Research UK Beatson Institute Advanced Technology Facilities grant no. A17196) related with **Figure 6C**; Richard Clark and the Wellcome Trust Clinical Research Facility (WTCRF) for the next generation sequencing service; Matthew Pearson (MRC Human Genetics Unit) for assistance with super resolution imaging; Graeme Grimes (MRC Human Genetics Unit) and the IGC bioinformatic core for the RNA-seq analysis pipeline related with **Table S1**. We thank Faith Robison (University of Washington in St. Louis) for helping us to establish histone mass spectrometry related with **Figure S3**; Jun Han (Metabolomics Innovation Centre, University of Victoria, Canada) for ^13^C_6_-glucose traced central carbon metabolites mass spectrometry assays related with **Table S3**; David Adams (Sanger Institute) for the human melanoma cell lines A375 and C089 engineered with stable expression of Cas9. We thank Craig Nicol and Uta Mackensen for the graphical assistance; Christina Lilliehook for editing assistance; the MRC HGU zebrafish facility; the MRC HGU Flow Cytometry facility and Technical Service staff; the MRC HGU imaging facility and Edinburgh Super-Resolution Imaging Consortium.

RSI is supported by an MRC Career Development award (MR/S007644/1) and the Simons Initiative for the Developing Brain (SFARI - 529085); AVK is funded by Wellcome Trust (Multiuser Equipment 208402/Z/17); ZK is funded by Melville Trust for Cancer Research (Studentship M00109.0001/TZH/MHR); FR is funded by Melanoma Research Alliance and the Wolfgang & Gertrud Boettcher Foundation; JAM is funded by the European Research Council (101001169). CJS thanks Cancer Research UK (C8717/A18245; C8717/A28285) and the Wellcome Trust (106244/Z/14/Z). The generation of patient derived cell lines was supported by funds from the Wellcome Trust (100282/Z/12/Z) to RM. EEP is funded by the Medical Research Council (MC_UU_00035/13), the European Research Council (ZF-MEL-CHEMBIO-648489), Melanoma Research Alliance and Rosetrees Trust (MRA Awards 687306, 917226). This work was supported by the Cancer Ressarch UK Scotland Centre (CTRQQR-2021\100006).

## Supplementary Figure Legends

**Supplementary Figure 1.**
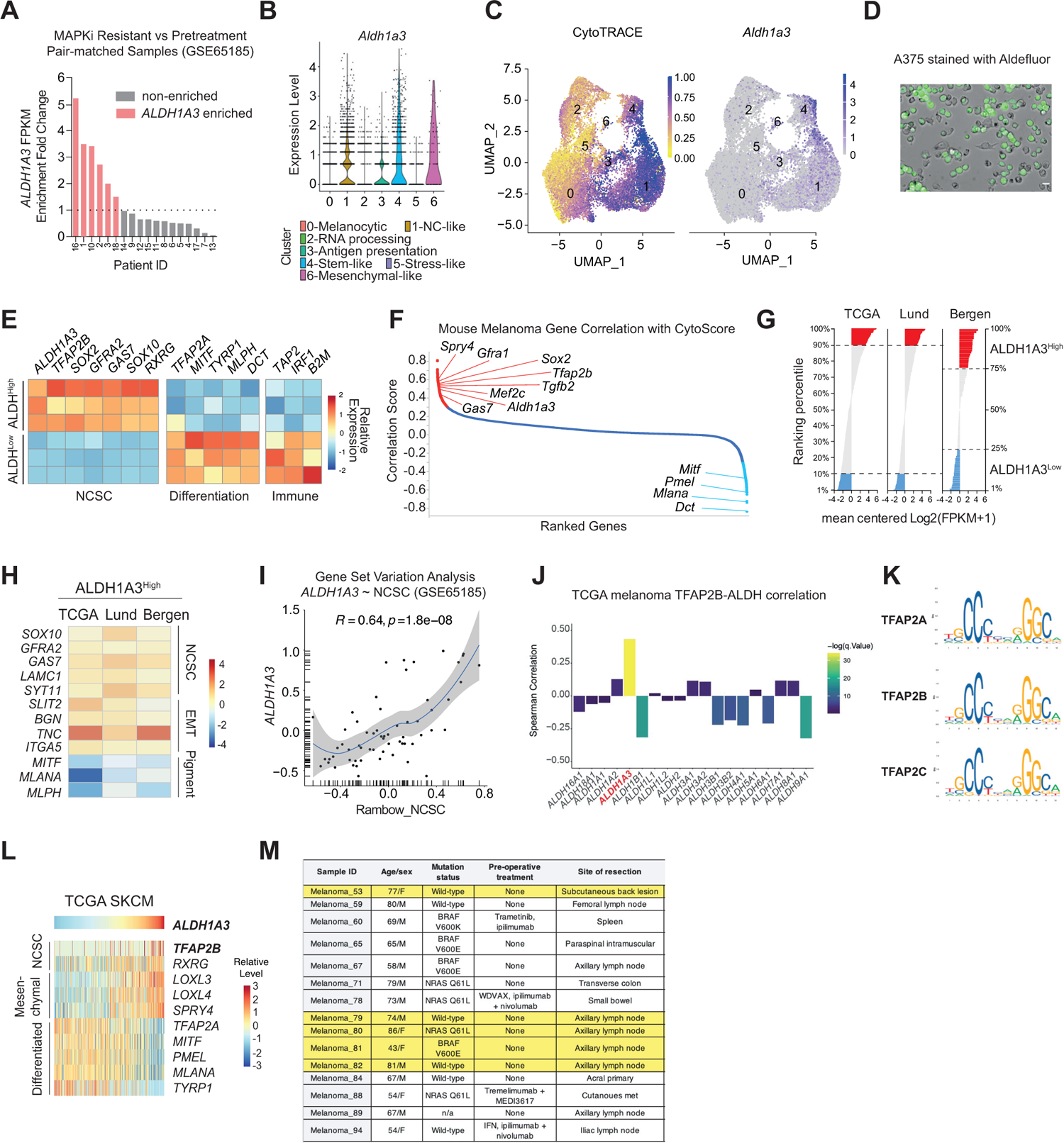
**A.** Pair-matched patient samples (GSE65185)^143^ ranked by *ALDH1A3* fold enrichment comparing post MAPKi resistance to pre-treatment biopsies. The average of *ALDH1A3* was used for comparison when multiple MAPKi-resistant tumour biopsies were taken from the same patient. **B.** Violin plot depicts *Aldh1a3* RNA expression in different murine melanoma cell states, including neural crest (NC)-like (cluster 1) and stem-like states (cluster 4)^43^. **C.** UMAP projection of murine melanoma cells colored by CytoTRACE score (gene expression diversity score, less differentiated = score close to 1 and more differentiated = score close to 0) and Aldh1a3 expression. Cluster numbers that relate to **B** are indicated. **D.** Microscope image overlaying brightfield and GFP channel shows the heterogeneous ALDH activity among A375 melanoma cells as measured by the Aldefluor assay on (adhered) live cells. Scale bar = 20 μm. **E.** Heatmap showing RT-qPCR results validating differentially expressed genes identified from RNA-seq on 3 bio-replicants of independently sorted A375 ALDH^High^ and ALDH^Low^ cells. Relative expression of each gene was averaged across 3 technical replicants and normalised to beta-actin loading control before heatmap plotting. For heatmap visualisation, each row (each gene) was scaled for the colour index filling. **F.** Murine melanoma genes from **B, C** ranked by decreasing correlation coefficient with CytoTRACE score (genes conferring a less differentiated phenotype close to 1 and genes conferring a differentiated phenotype close to −1). **G.** Patient samples ranked by *ALDH1A3* expression, with the top and bottom 10% ranking samples of TCGA and Lund datasets defined as ALDH1A3^High^ and ALDH1A3^Low^ group respectively, while the top and bottom 25% ranking samples of Bergen datasets were defined as ALDH1A3^High^ and ALDH1A3^Low^ groups due to fewer patient numbers in the Bergen cohort. **H.** Heatmap showing gene expression fold enrichment levels comparing the ALDH1A3^High^ to ALDH1A3^Low^ patient groups. Colorimetric values normalized by column (each cohort). **I.** Scatter plot showing the correlation of *ALDH1A3* and NCSC gene set expression (measured by gene set variation analysis) over different human melanoma patient samples (GSE65185). Spearman’s rank correlation score (*R*) and probability (p) value. **J.** Bar plot showing spearman correlation value of each *ALDH* isoform with *TFAP2B* in TCGA dataset. Colour filling scaled by FDR.q.value. Only *ALDH1A3* shows significant positive correlation with TFAP2B. **K.** DNA Binding motif sequences of TFAP2 family members from JASPAR database. **L.** *ALDH1A3* gene network heatmap. Heatmap of TCGA melanoma patient samples with RNA expression of *TFAP2B*, *RXRG* and mesenchymal genes, as well as *MITF*, *TFAP2A*, and their target genes ranked by *ALDH1A3* level. Colour index filling scaled by row (each gene). **M.** Mutation and clinical background patient samples table from Tirosh et al., 2016^51^. The cluster of *ALDH1A3-TFAP2B* consists of cells from highlighted patient entries.

**Supplementary Figure 2.**
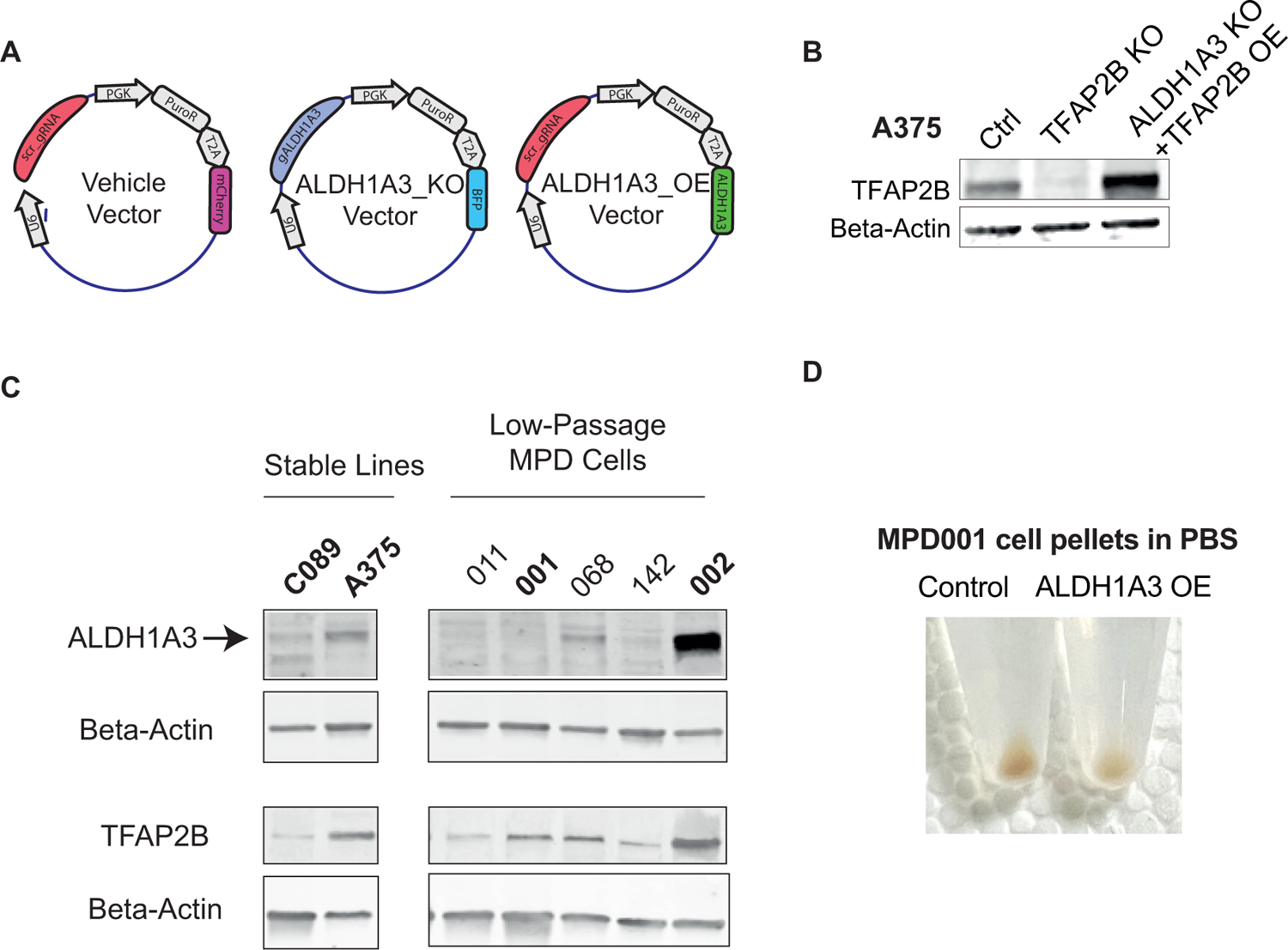
**A.** Schematics of plasmids used to generate wildtype (WT), *ALDH1A3* knockout (KO), and *ALDH1A3* overexpression (OE) cell lines. **B.** Western blot of TFAP2B levels in *TFAP2B* knockout cells and *ALDH1A3* knockout cells overexpressing *TFAP2B.* Beta-Actin is presented as a loading control. **C.** Western blot of ALDH1A3 and TFAP2B levels across established stable melanoma cell lines used for this work (C089 and A375) as well as a panel of MPD low passage cells. **D.** ALDH1A3 overexpression leads to less pigmentation. Cell pellets showing brown melanin in control and ALDH1A3 overexpressing MPD001 cells.

**Supplementary Figure 3.**
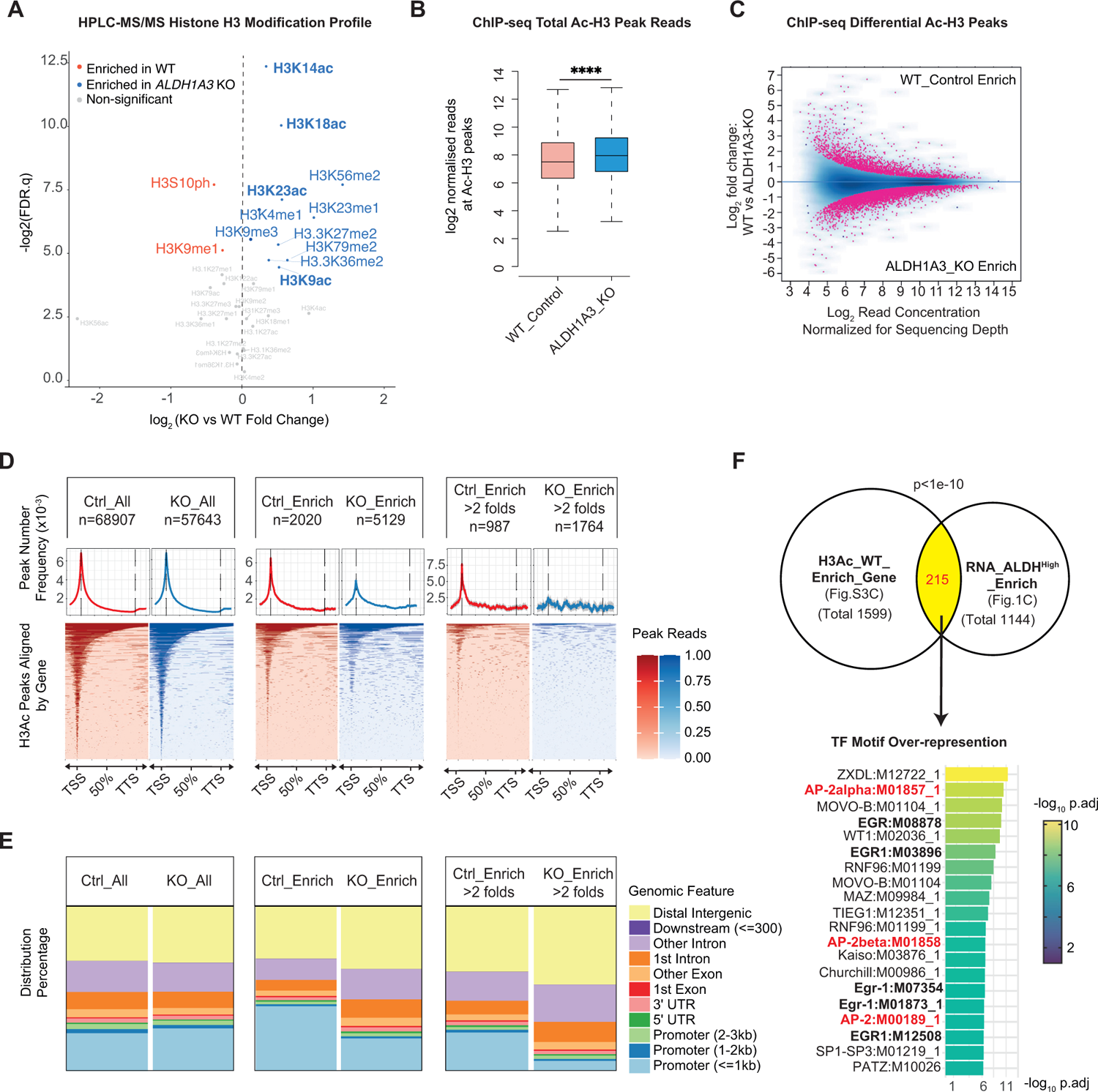
**A.** Volcano plot of histone H3 peptide modification quantification measured by bottom-up histone HPLC-MS/MS. (n=5, FDR q values by two-way ANOVA corrected with Sidak’s multiple comparison method.) Acetyl-histone H3 lysine residuals with significant upregulation in *ALDH1A3 KO* group comparing to the control are highlighted in **Bold**. **B.** Collective acetyl-histone H3 reads abundance at peak regions measured by acetyl-histone H3 ChIP-seq. (Wilcoxon signed-rank test. ****P<0.0001). **C.** Volcano plot of differentially acetylated histone H3 peaks in WT control and ALDH1A3 knockout (highlighted by magenta, fold change >1, FDR.q < 0.05). **D. E. (D)** Heatmap of ChIP-seq histone pan-acetyl-histone H3 peaks aligned by gene body extended by 20% upstream transcription starting site (TSS) and downstream transcription termination site (TTS) (bottom panels), with average profile showing read count frequency distribution (upper panels) and **(E)** genomic annotations by bar plot (top panels) among A375 control (Ctrl_All), ALDH1A3 knockout (KO_All), significantly enriched peaks in A375 control (Ctrl_Enrich, Ctrl_Enrich>2 folds), and significantly enriched peaks in ALDH1A3 knockout (KO_Enrich, KO_Enrich>2 folds). **F.** (Upper panel) Venn diagram of overlapping genes between upregulated genes by RNA-Seq in sorted ALDH^High^ cells (Figure 1C), and genes enriched in control cells versus *ALDH1A3* knockout cells by histone H3 acetylation in **B**. P values by Fisher’s exact test. (lower panel) Transcription factor binding motif over-representation analysis of the Venn diagram overlapping genes ranked by enrichment score calculated from g:Profiler (version e109_eg56_p17_1d3191d) with g:SCS multiple testing correction method applying statistical significance threshold of 0.05 ^111^.

**Supplementary Figure 4.**
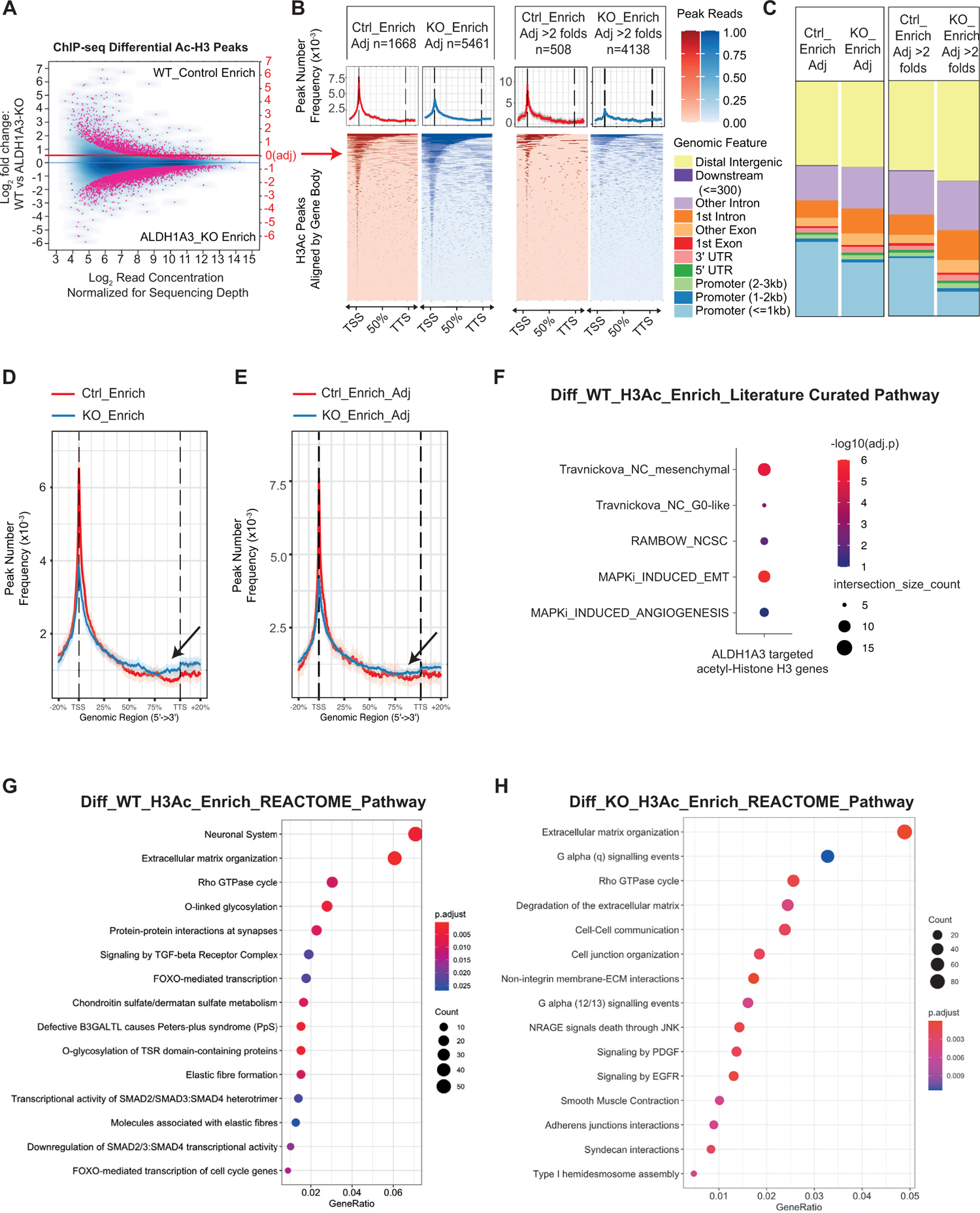
**A.**Schematics of adjusted (right Y axis, highlighted in red) differential peak analysis on acetylated histone H3 peaks in WT control compared to *ALDH1A3* knockout cells. Here, we have manually increased log base 2 peak score by +0.5 in *ALDH1A3* KO samples to account for the potential biased loss of peak score during normalisation. **B.**Heatmap of adjusted differentially acetylated histone H3 peaks aligned by the gene body and extended by 20% upstream transcription starting site (TSS) and downstream of the transcription termination site (TTS) (bottom panels). Read count frequency distribution is shown in an average profile (upper panels). **C.**Genomic annotations by bar plot among adjusted enriched peaks in A375 control (Ctrl_Enrich_Adj, Ctrl_Enrich_Adj>2 folds), and in *ALDH1A3* knockout cells (KO_Enrich_Adj, KO_Enrich_Adj>2 folds). **D-E.** Overlay of the differentially enriched acetyl-histone H3 peaks aligned by gene body from Figure 4C and **(E)** from **Figure S4B.** Both showed reduced proportion of TSS-centring peaks in *ALDH1A3* KO cells, coupled with increased proportion of peaks towards the TTS and extended gene downstream regions as highlighted by arrows. **F-H.** Significant over-representation gene terms with literature curated gene lists **(F).** Reactome gene pathway datasets enriched in control **(G)** and *ALDH1A3* knockout **(H)** cells. FDR <0.05, Benjamini-Hochberg test.

**Supplementary Figure 5.**
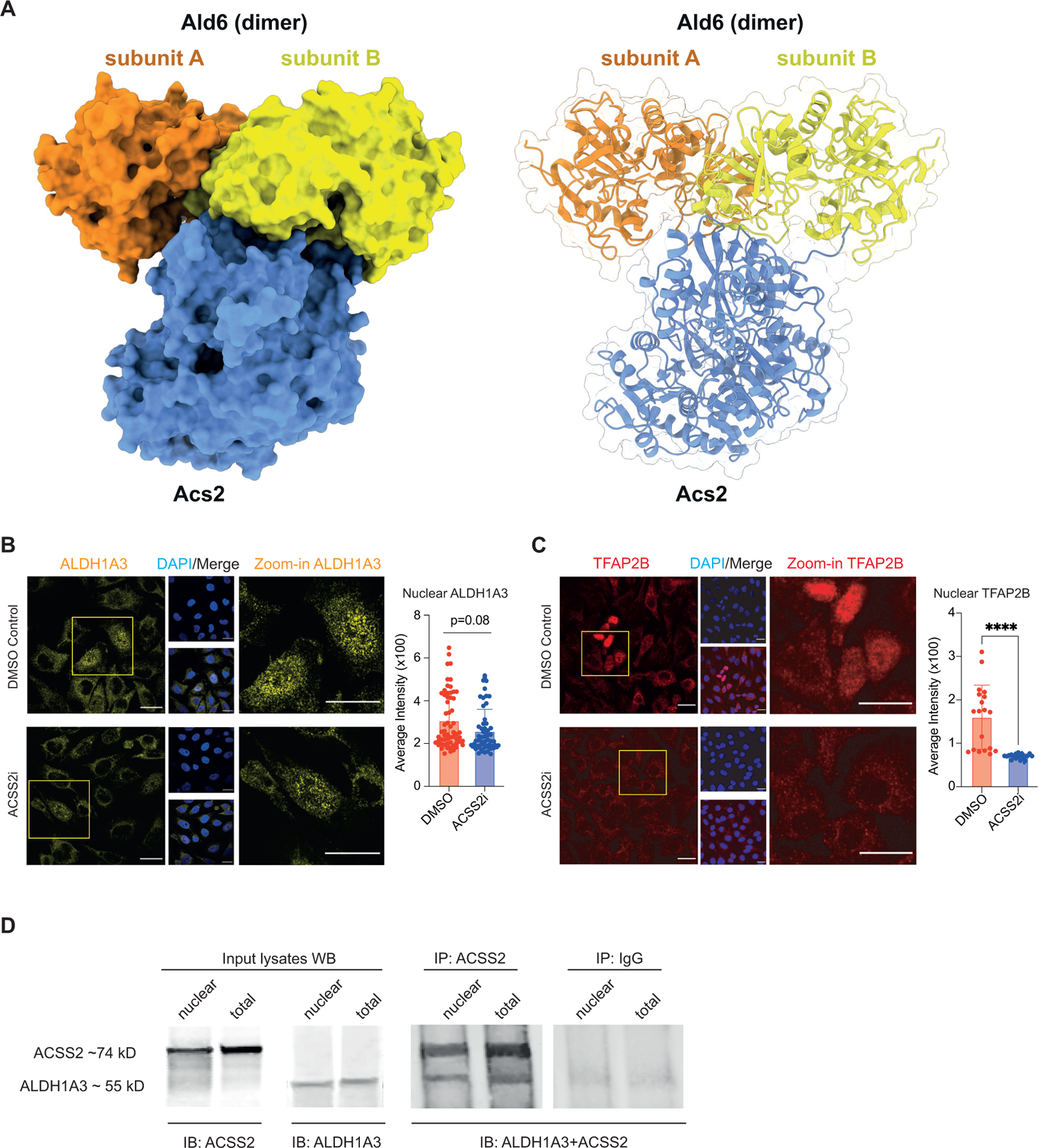
**A.** AlphaFold multimer modelling of yeast Ald6 and Acs2 proteins visualised in surface model (left) and ribbon model (right). **B-C**. ICC staining by fluorescence antibody for ALDH1A3 (yellow, in **B**), and TFAP2B (red, in **C**) in sorted A375 ALDH^High^ and ALDH^Low^ cells. DAPI (blue). Scale bar = 20 μm. Fluorescence signal intensity quantification of nuclear **(B)** ALDH1A3 and **(C)** TFAP2B in ICC images. n = 60 single cells for ALDH1A3 and n=19 single cells for TFAP2B quantification (represented as individual dots), mean±s.d., unpaired non-parametric Kolmogorov-Smirnov test. ****P<0.0001. **D.** Western blot of nuclear or total A375 cell lysates probing either ACSS2 or ALDH1A3 levels of the input materials used for co-immunoprecipitation of ACSS2 (IP: ACSS2 and IP: IgG). The co-immunoprecipitated materials were examined by immunoblotting ALDH1A3 as well as ACSS2. Immunoprecipitation of ACSS2 successfully captured ALDH1A3, and this was not detected in the IgG control. For each condition (isolated nuclei versus whole cell department), the input, IP: ACSS2, and IP: IgG materials were divided from the same vial of harvested A375 cells.

**Supplementary Figure 6.**
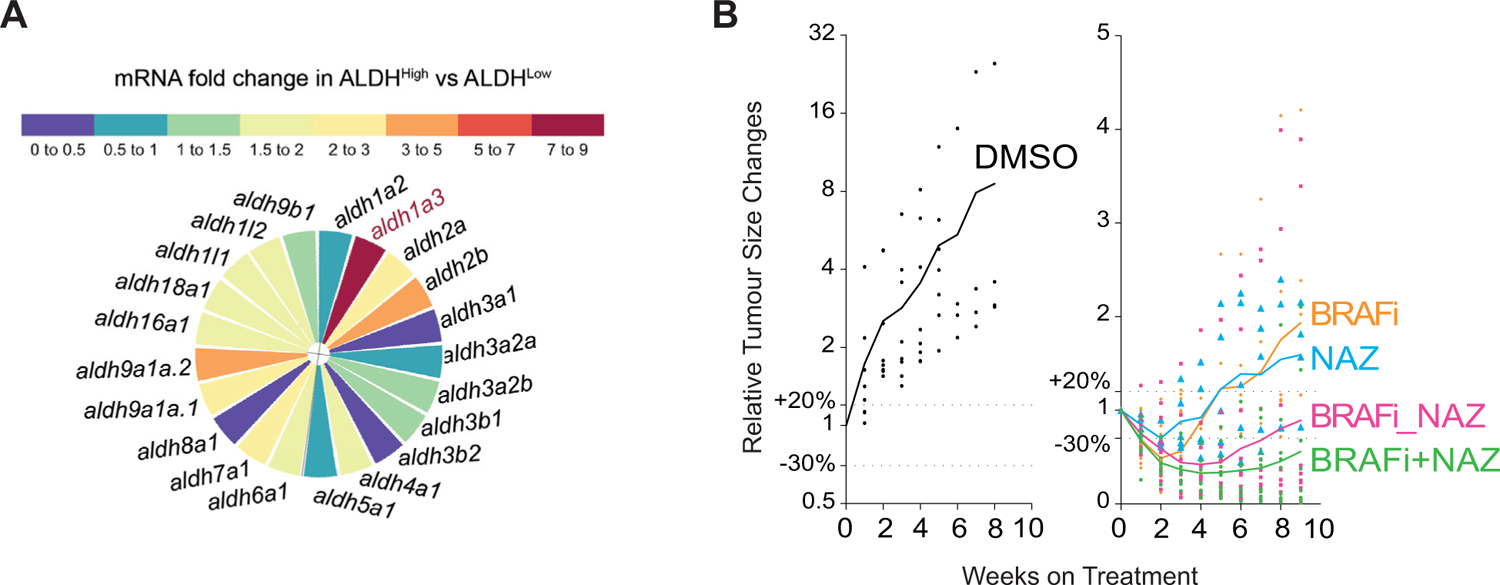
**A.**RT-qPCR quantification shown by a heat rose map of *aldh1a3* expression enriched with the highest folds in zebrafish ALDH^High^ versus ALDH^Low^ melanoma cells. (n=3 bio-replicates each with 3 technical replicates, Multiple paired t-test corrected with Holm-Sidak’s method). **B.** Spaghetti summary plot related to Figure 7H, showing tumour volume changes during drug trial design as listed in Figure 7G. DMSO: melanomas n=8, fish N=5; BRAFi: melanomas n=5, fish N=4; NAZ: melanomas n=6, fish N=4. BRAF_NAZ: melanomas n=12, fish N=5; BRAFi + NAZ: melanomas n=10, fish N=4. Each dot on the plot represents one melanoma lesion. Drug pellet treatment is colour coded as shown in **G**: DMSO (Black): daily DMSO control treatment. BRAFi (Orange): 200 mg/kg/day vemurafenib treatment. NAZ (Blue): daily 150 mg/kg/day Nifuroxazide treatment. BRAFi_NAZ (Magenta): 3-week treatment of 200 mg/kg/day vemurafenib followed by daily 150 mg/kg/day Nifuroxazide treatment. BRAFi+NAZ (Green): 3-week treatment of 200 mg/kg/day vemurafenib followed by the combination of 200 mg/kg/day vemurafenib and 150 mg/kg/day Nifuroxazide treatment.

**Supplementary Figure 7.**
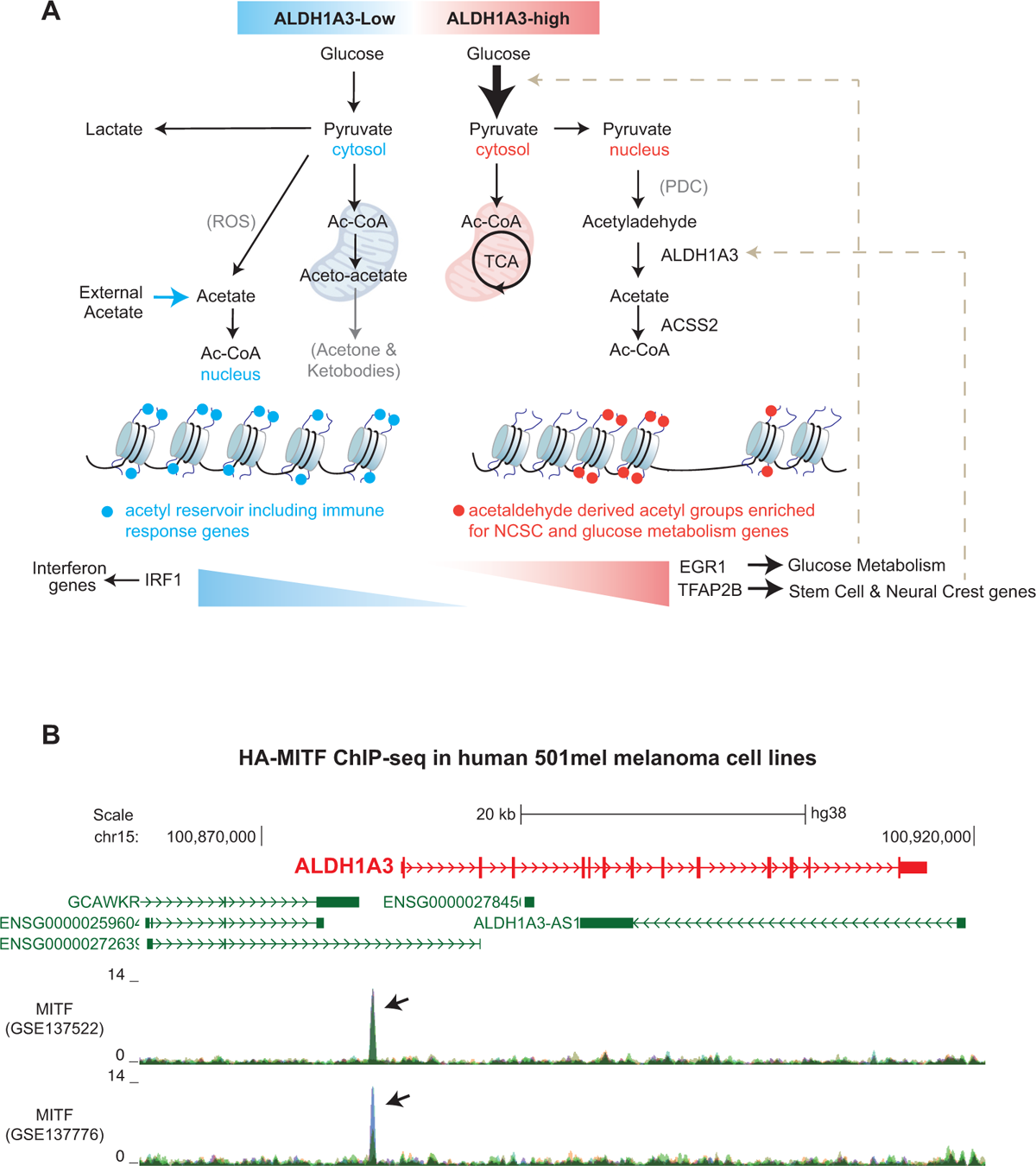
**A.** Schematic model of ALDH1A3 orchestrating the metabolic, epigenetic, and transcriptional cell states of melanoma. **B.** UCSC genome browser snapshot of MITF ChIP-seq multiWig tracks at the ALDH1A3 locus from two experiments (Top, GSE137522; Bottom, GSE137776). Each track is an overlay of an input control and 2 independent HA-MITF ChIP-seq experiments in the human 501mel melanoma cell line engineered to inducibly express HA-tagged MITF. Inputs are shown in Green Mist colour RGB (200,200,150).

## Supplementary Extended Data and Text

### Supplementary extended text for Figure S3 and S4

To determine how ALDH1A3-dependent histone H3 acetylation is deposited on chromatin, we first performed quantitative acetyl-histone H3 chromatin immunoprecipitation (ChIP)-seq using an antibody against pan-histone H3 acetylated sites (K9 + K14 + K18 + K23 + K27). We found that the average acetyl-histone H3 signal increased by ∼1.4 fold across H3-ac peak regions in *ALDH1A3* KO cells **(Figure S3B;** consistent with our western blotting and mass spectrometry analysis, Figure 3K**, L; Figure S3A),** and identified 5129 acetyl-histone H3 peaks enriched in *ALDH1A3* KO cells, versus 2020 peaks enriched in *ALDH1A3* control cells (fold change >1, FDRq < 0.05) **(Figure S3C, D).** Notably, in control cells, the enriched acetyl-histone H3 peaks were clustering around transcription start sites (TSSs), especially within 1kb of promoters, whereas in *ALDH1A3* KO cells, the enriched acetyl-histone H3 peaks were broadly dispersed throughout the genome, and particularly spreading into the distal intergenic region and intronic regions **(Figure S3D, E)**. This distribution effect is even more prominent when comparing the top fold-change enriched acetyl-histone H3 peaks (>2 fold) between control and *ALDH1A3* KO cells **(Figure S3D, E)**. To test the robustness of our differential acetyl-histone H3 analysis, we adjusted the average peak score of *ALDH1A3* KO samples by manually increasing the log base 2 value by +0.5, ∼ log2(1.4) **(Figure S4A)**, which is the fold change determined by western blot and mass spec (Figure 3K**, L; Figure S3A).** With the adjusted peak score (a more stringent analysis), enriched acetyl-histone H3 remained distributed at distal intergenic and intronic regions in the *ALDH1A3* KO, in contrast with the enriched acetyl-histone H3 in promoter areas in control cells **(Figure S4B-E).**

To determine the mechanistic basis of information flow from acetyl-histone H3 to transcription in cells with high ALDH1A3, we compared genes with promoters marked by high acetyl-histone H3 (total 1499) with the RNA-seq from Figure 1C. We found that ∼19% of genes with enriched expression in ALDH1A3^High^ (total 1144) have ALDH1A3-driven acetyl-histone H3 in their promoters (215, p<1e-10) **(Figure S3F)**.

In addition, and consistent with our prior observations (Figure 1, Figure 3), pathway enrichment analysis revealed that neural crest and stem cell state marker genes, neuronal signalling, TGF-beta signalling, and O-glycosylation pathways were over-represented in ALDH1A3^High^ specific acetyl-histone H3 states **(Figure S4F, G).** Pathways enriched in acetyl-histone H3 sites in ALDH1A3 KO cells also included cell-cell junction terms **(Figure S4H)**, possibly reflecting the difference in cell morphology between the rounded ALDH^High^ cells and the flat, elongated ALDH^Low^ cells (Figure 1B). Together, these observations support that the high-glucose flux and NCSC transcriptional states promoted by high ALDH activity arise from selective histone H3 acetylation.

### Supplementary extended data for the synthesis and NMR spectrum for AC-148

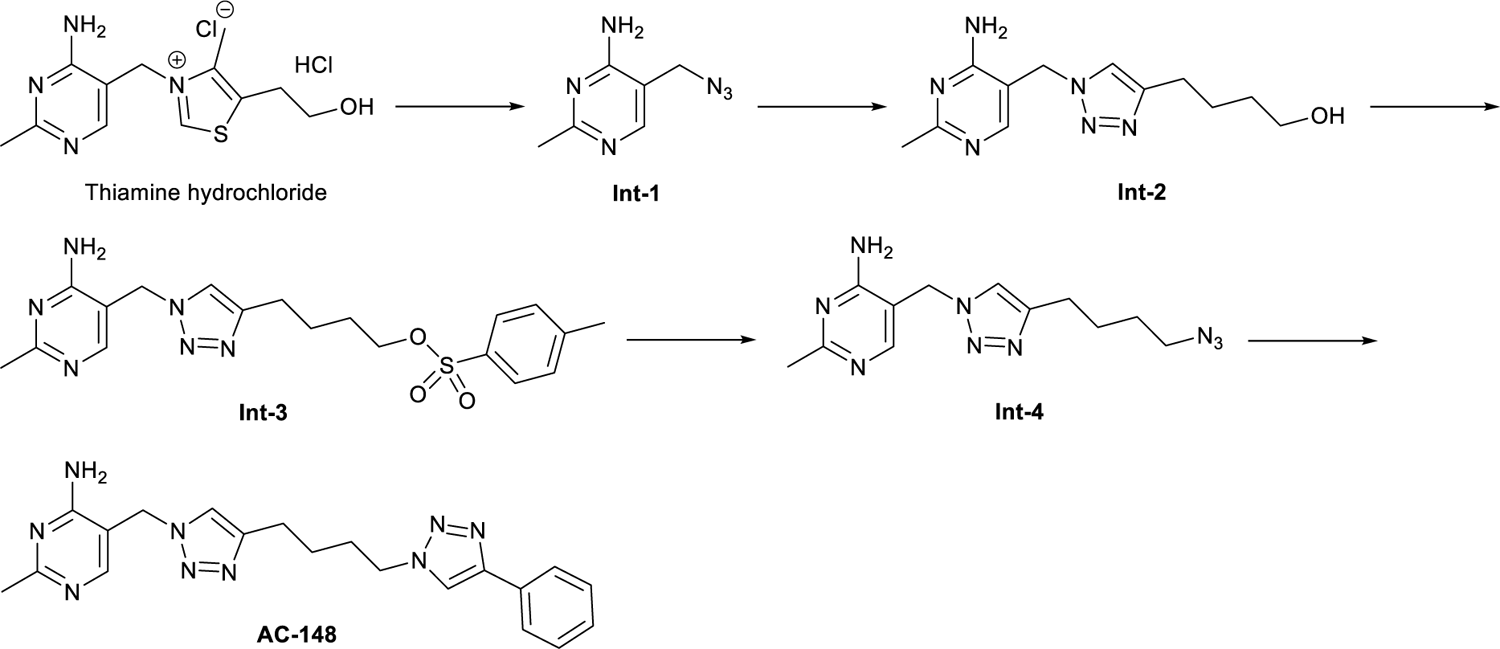

#### 5-(Azidomethyl)-2-methylpyrimidin-4-amine Int-1

To a stirred solution of thiamine hydrochloride (10.0 g, 29.7 mmol) and NaN_3_ (5.0 g, 76.9 mmol) in water (90 mL, 0.33 M) was added Na_2_SO_3_ (0.4 g, 3.2 mmol). The resultant mixture was stirred at 70 °C for 18 hours, then acidified with citric acid monohydrate (10.5 g) to pH 4-5, washed with DCM (200 mL), and basified with K_2_CO_3_ to pH 8-10. Upon product precipitation, the suspension was filtered under reduced pressure. The residue was rinsed with cold water and dried under reduced pressure to yield **Int-1** as a white solid (3.4 g, 70%). **^1^H NMR** (400 MHz, CD_3_OD) δ 8.02 (s, 1H), 4.32 (s, 2H), 2.43 (s, 3H). Analytical data are consistent with those reported (Chan et al., 2023).

#### 4-{1-[(4-Amino-2-methylpyrimidin-5-yl)methyl]-1H-1,2,3-triazol-4-yl}butan-1-ol Int-2

To a stirred solution of **Int-1** (1322 mg, 8.0 mmol) and 5-hexyn-1-ol (1020 mg, 10.4 mmol) in *t*-BuOH and water (24 + 8 mL, 0.25 M) was added CuSO_4_.5H_2_O (60 mg, 0.24 mmol) and sodium ascorbate (475 mg, 2.4 mmol). The resultant mixture was stirred at 40 °C for 40 hours, concentrated under reduced pressure, diluted with CHCl_3_/*i*-PrOH (3:1, 50 mL), washed with 0.1 M K_2_CO_3_ (50 mL), dried over anhydrous Na_2_SO_4_, filtered, and evaporated under reduced pressure. The residue was purified by silica flash chromatography (10% MeOH in DCM) to yield **Int-2** as a white solid (1366 mg, 65%). **^1^H NMR** (400 MHz, CD_3_OD) δ 8.03 (s, 1H), 7.80 (s, 1H), 5.47 (s, 2H), 3.56 (t, 2H, *J* = 6.5 Hz), 2.71 (t, 2H, *J* = 7.7 Hz), 2.42 (s, 3H), 1.73 (m, 2H), 1.57 (m, 2H). Analytical data are consistent with those reported (Chan et al., 2023).

#### 4-{1-[(4-Amino-2-methylpyrimidin-5-yl)methyl]-1H-1,2,3-triazol-4-yl}butyl 4-methylbenzene-1-sulfonate Int-3

To a stirred solution of **Int 2** (1337 mg, 5.1 mmol) in dry pyridine (25.5 mL, 0.2 M) under nitrogen at 0 °C was added *p*-TsCl (4850 mg, 25.5 mmol) in three portions. The resultant mixture was stirred at 25 °C for 4 hours, quenched with cold 1 M HCl (20 mL), diluted with water (10 mL), neutralised with NaHCO_3_ to pH 7, and extracted with DCM (150 mL). The organic phase was washed with sat. aq. Cu_2_SO_4_ (100 mL), dried over anhydrous Na_2_SO_4_, filtered, and evaporated under reduced pressure. The residue was purified by silica flash chromatography (10% MeOH in DCM) to yield **Int-3** as a pale-yellow semi-solid (1063 mg, 50%). **^1^H NMR** (400 MHz, CD_3_OD) δ 8.04 (s, 1H), 7.78 (d, 2H, *J* = 7.8 Hz), 7.75 (s, 1H), 7.43 (d, 2H, *J* = 7.8 Hz), 5.46 (s, 2H), 4.04 (t, 2H, *J* = 5.5 Hz), 2.63 (t, 2H, *J* = 6.5 Hz), 2.45 (s, 3H), 2.42 (s, 3H), 1.65 (m, 4H). Analytical data are consistent with those reported (Chan et al., 2023).

#### 5-{[4-(4-Azidobutyl)-1H-1,2,3-triazol-1-yl]methyl}-2-methylpyrimidin-4-amine Int-4

To a stirred solution of **Int 3** (1041 mg, 2.5 mmol) in dry DMF (2.5 mL, 1 M) under nitrogen was added NaN_3_ (325 mg, 5.0 mmol). The resultant mixture was stirred at 25 °C for 40 hours, quenched with 0.1 M K_2_CO_3_ (50 mL), and extracted with CHCl_3_/*i*-PrOH (3:1, 50 mL). The organic phase was dried over anhydrous Na_2_SO_4_, filtered, and evaporated under reduced pressure. The residue was purified by silica flash chromatography (10% MeOH in DCM) to yield **Int-4** as a white foam (445 mg, 62%). **^1^H NMR** (400 MHz, CD_3_OD) δ 8.04 (s, 1H), 7.81 (s, 1H), 5.47 (s, 2H), 3.33 (t, 2H, *J* = 6.7 Hz), 2.74 (t, 2H, *J* = 7.6 Hz), 2.42 (s, 3H), 1.74 (m, 2H), 1.62 (m, 2H). Analytical data are consistent with those reported (Chan et al., 2023).

#### 2-Methyl-5-({4-[4-(4-phenyl-1H-1,2,3-triazol-1-yl)butyl]-1H-1,2,3-triazol-1-yl}methyl)pyrimidin-4-amine AC-148

To a stirred solution of Int-4 (86 mg, 0.3 mmol) and phenylacetylene (31 mg, 0.3 mmol) in *t*-BuOH and water (0.9 + 0.3 mL, 0.25 M) was added CuSO_4_.5H_2_O (2.5 mg, 0.01 mmol) and sodium ascorbate (20 mg, 0.1 mmol). The resultant mixture was stirred at 40 °C for 72 hours, concentrated under reduced pressure, diluted with EtOAc (50 mL), washed with 0.1 M K_2_CO_3_ (50 mL), dried over anhydrous Na_2_SO_4_, filtered, and evaporated under reduced pressure. The residue was purified by silica flash chromatography (10% MeOH in DCM) to yield **AC-148** as a white solid (41 mg, 35%). **^1^H NMR** (400 MHz, CD_3_SOCD_3_) δ 8.57 (s, 1H), 7.94 (s, 1H), 7.82-7.86 (m, 3H), 7.41-7.47 (m, 2H), 7.30-7.35 (m, 1H), 6.87 (br, 2H, NH_2_), 5.37 (s, 2H), 4.42 (t, 2H, *J* = 6.9 Hz), 2.65 (t, 2H, *J* = 7.8 Hz), 2.30 (s, 3H), 1.86-1.96 (m, 2H), 1.54-1.63 (m, 2H). **HRMS** (ESI) m/z: [M+H^+^] calculated for C20H24N9: 390.2149; found: 390.2146. Analytical data are consistent with those reported (Chan et al., 2023).

^1^H NMR of **AC-148** in CD_3_SOCD_3_:

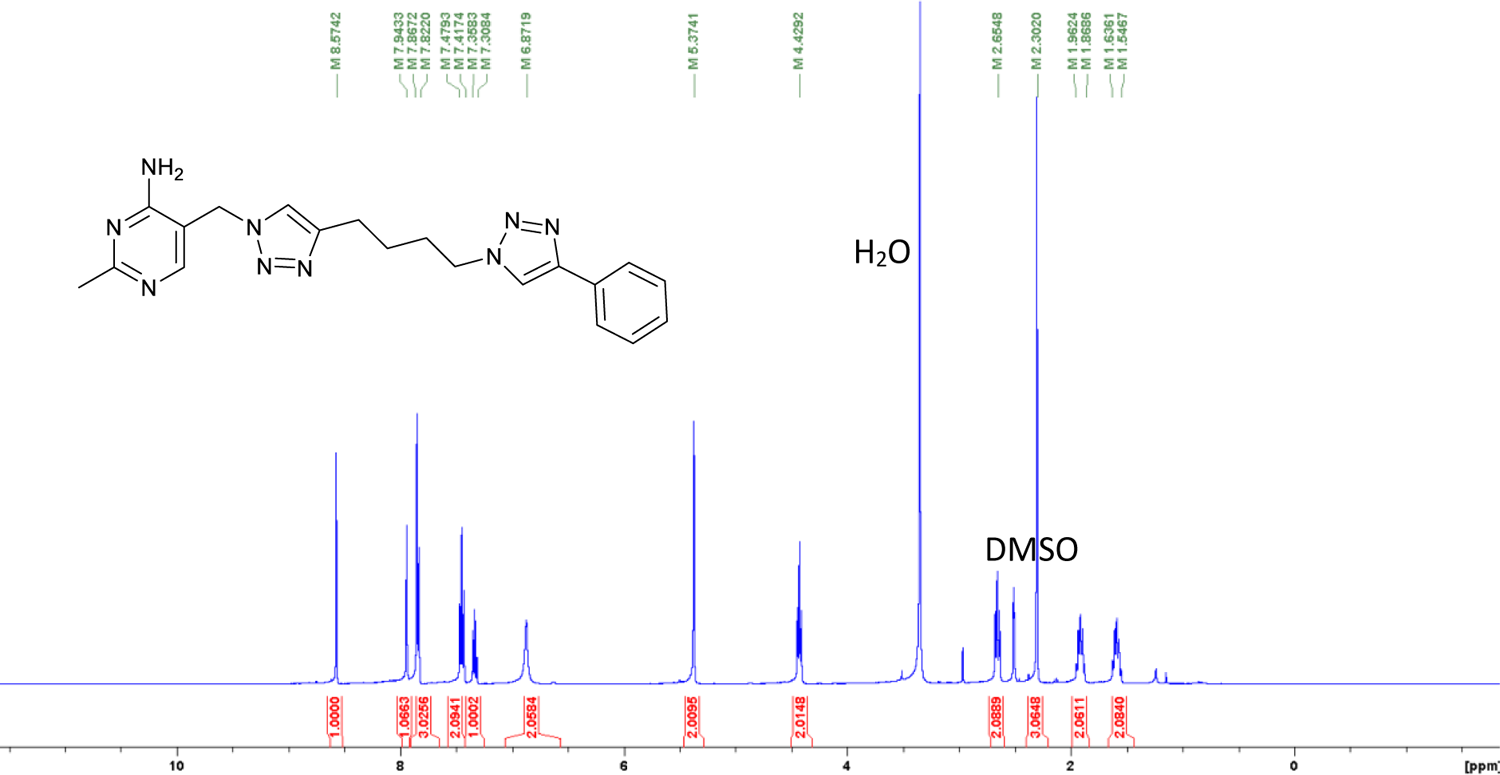

